# An Integrated Single-Cell Atlas Reveals Hepatic Stellate Cell Heterogeneity and Spatiotemporal Dynamics after Liver Injury

**DOI:** 10.1101/2025.11.25.690307

**Authors:** Jiongliang Wang, Jiawang Tao, Cuicui Xia, Miaoxiu Tang, Andrei-Florian Stoica, Shengxian Yuan, Yinxiong Li, Lin Guo, Xiangqian Kong, Jie Wang

## Abstract

**Background:** Hepatic stellate cells (HSCs) orchestrate fibrosis-free repair after acute liver injury (ALI) and sustain fibrogenesis in chronic liver injury (CLI). However, the heterogeneity and spatiotemporal dynamics of HSCs across these different injury models remain elusive. This study sought to construct a cross-etiological HSC atlas to delineate HSC states and transitions during liver injury.

**Methods:** We integrated 86,072 single-cell transcriptomes from 84 mouse samples across four etiologies and validated findings using multi-omics data from 277 mouse and 798 human samples. Cellular dynamics were characterized through clustering, trajectory inference, spatial analysis, and multicellular coordination network analysis. Experimental validation included liver injury models and gain-of-functional assays in primary mouse HSCs and LX-2 cells.

**Results:** We established a cross-etiological, spatiotemporally resolved HSC atlas comprising 11 subpopulations. Trajectory analysis delineated a continuous quiescence-activation-attenuation (QAA) trajectory, recapitulating the *in vivo* full spectrum of state transitions and being supported by sequential pathway activation validated *in vitro*. In ALI, HSCs spatiotemporally completed the QAA trajectory around injury zones, whereas collagen-producing *S100a6*⁺ HSCs pathologically accumulated in mouse and human CLI due to trajectory dysregulation. Notably, the atlas identified a previously unrecognized apoptosis-prone *Mrc2*⁺ HSC subtype strongly co-localized with p53 in both mice and humans. Overexpression of transcription factors confirmed that *Hbp1*, *Tbx20*, *Atoh8*, and *Plagl1* enriched in *Mrc2*^+^ HSCs promoted HSC apoptosis. Finally, we revealed the microenvironment of distinct cellular modules which coordinated HSC progression along the QAA trajectory. A 531-gene signature derived from the inflammatory-fibrotic cellular module significantly correlated with fibrosis stage and hepatocellular carcinoma risk in human cohorts.

**Conclusions:** We established a comprehensive HSC atlas and delineates HSC heterogeneity and spatiotemporal dynamic across etiologies. Dysregulation of the QAA trajectory underlies fibrotic progression, providing a resource for identifying antifibrotic targets.

## Introduction

Liver fibrosis, a progressive scarring response to chronic liver disease, represents a major global health burden with limited therapeutic options [1–3]. Hepatic stellate cells (HSCs) are recognized as a key driver of liver fibrogenesis [4–6]. The quiescent HSCs (qHSCs) maintain metabolic homeostasis [6, 7], but upon injury they become activated (aHSCs) and produce extracellular matrix, promoting fibrosis. Notably, HSCs exhibit remarkable plasticity. Following acute liver injury (ALI), HSCs undergo transient activation and subsequently resolve through apoptosis or become inactivated HSCs (iHSCs), enabling fibrosis-free repair[8–12]. In contrast, chronic liver injury (CLI) leads to persistent activation and pathological accumulation of fibrogenic aHSCs[3, 5, 6, 9]. The mechanisms that enable complete resolution after ALI but disrupted during CLI remain largely elusive.

Recent studies using single-cell or single-nucleus RNA sequencing (sc/snRNA-seq) have revealed heterogeneous HSC subpopulations, including spatially distinct quiescent cells and diverse types of activated cells characterized by inflammatory, contractile, and collagen-producing functions[4–6, 9, 13–15]. Although dozens of HSC subpopulations have been annotated for different injury contexts, a systematic identification and comparison of these subtypes within a unified, cross-etiological atlas is still lacking. Most previous studies have focused on single etiologies or discrete stages[13–22], leaving no continuous model of HSC transitions from quiescence through activation to inactivation. Although apoptosis contributes to fibrosis regression[1, 23], the molecular identity, regulatory mechanisms, and *in vivo* transitions of apoptosis-related HSCs remain poorly understood. Furthermore, how intrinsic HSC programs are coordinated with multicellular microenvironments to determine fibrosis resolution or persistent fibrosis is largely unknown.

Here, we systematically compared the fibrosis-free repair model of ALI and CLI to delineate how HSC programs become dysregulated and maladaptive in CLI. By integrating over 80,000 mouse HSCs from seven independent scRNA-seq datasets, along with spatial RNA sequencing (spRNA-seq) data, single-nucleus ATAC-seq (snATAC-seq) data, and validation in human HSCs, we constructed a cross-etiological, spatiotemporally resolved HSC atlas and delineated a conserved quiescence-activation-attenuation (QAA) trajectory. Within this landscape, we identified a previously underappreciated apoptosis-prone *Mrc2*^high^ HSC subtype likely critical for fibrosis regression. CLI disrupted the QAA trajectory, leading to maladaptive accumulation of collagen-producing HSCs, while multicellular coordination modules aligned spatiotemporally with HSC dynamics. Together, this study provides a comprehensive landscape of HSC plasticity and suggests that liver fibrosis arises not only from persistent activation but also from the dysfunction of QAA trajectory. It also offers a comparative and integrative perspective to guide further mechanistic and translational studies.

## Results

Cross-Etiological Single-Cell Integration Reveals 11 HSC Subtypes including an Apoptosis-prone *Mrc2*^high^ Subtype To comprehensively characterize HSC heterogeneity and dynamics across diverse pathological contexts, we implemented a strategy combining integrative analysis, multi-omics cross-validation and experimental verification (**Figure 1A; Table S1**).

**Figure 1.**
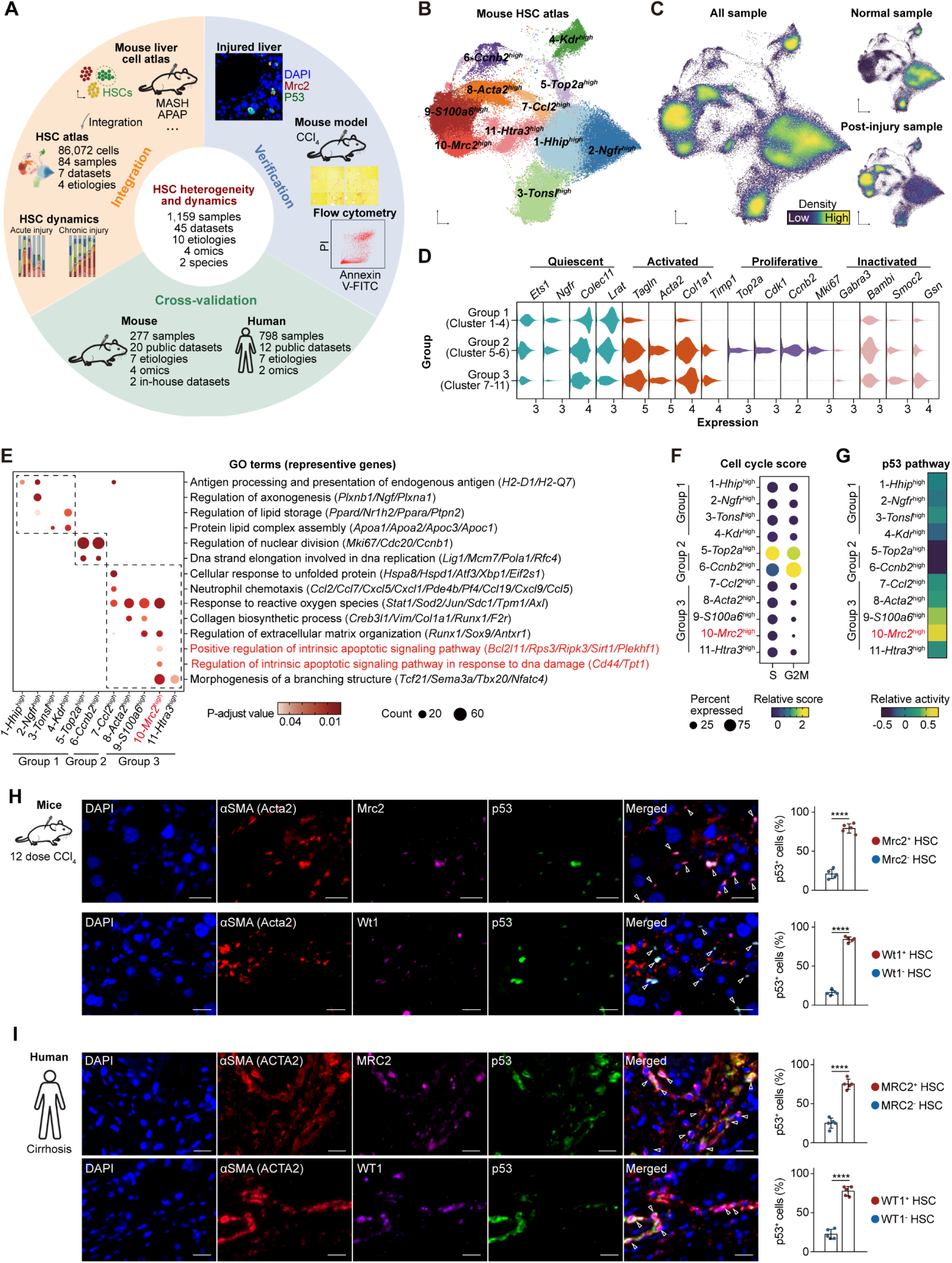
Landscape of HSC states across diverse liver injury models. **(A)** Schematic overview of the study design, illustrating the integrative framework combining mouse and human liver datasets through integration, cross-validation, and experimental verification. **(B)** Uniform manifold approximation and projection (UMAP) visualization of the integrated mouse HSC atlas encompassing multiple etiologies and injury stages. **(C)** UMAP density plots showing the spatial distribution of HSCs in the entire dataset, and in normal versus post-injury conditions. **(D)** Violin plots showing the expression of representative markers defining quiescent, activated, proliferative, and inactivated HSC states across the quiescent (Group 1: clusters 1-4), proliferative (Group 2: clusters 5-6), and injury-responsive (Group 3: clusters 7-11) groups. **(E)** Dot plot of significantly enriched gene ontology (GO) terms with representative genes. Apoptosis-related pathways are highlighted in red. **(F)** Bubble plot showing cell-cycle scores across HSC subtypes. **(G)** Heatmap depicting PROGENy-inferred p53 pathway activity among HSC subtypes. **(H)** Representative immunofluorescence images (left) and quantification (right, n = 5) showing that αSMA⁺Mrc2⁺ (top) and αSMA⁺Wt1⁺ (bottom) HSCs display high co-expression with p53 protein in CCl₄-injured mouse livers (12 doses). Data represent mean ± SD. Scale bars, 20 μm. **(I)** Representative immunofluorescence images (left) and quantification (right, n = 5) showing that αSMA⁺MRC2⁺ (top) and αSMA⁺WT1⁺ (bottom) HSCs also display robust co-expression with p53 protein in cirrhotic human livers. Data represent mean ± SD. Scale bars, 20 μm. Statistical significance is denoted as: *, p < 0.05; **, p < 0.01 (two-sided Wilcoxon test).

This framework incorporated 1,159 samples of 45 datasets from multiple technologies (RNA-seq, sn/scRNA-seq, spRNA-seq, and snATAC-seq) across 10 etiologies and two species (human and mouse), aiming to ensure the robustness and translational relevance of our findings.

We first integrated seven published mouse scRNA-seq datasets[13, 14, 16, 19, 20, 24, 25], spanning ALI and CLI from diverse liver injury etiologies (e.g., MASH, metabolic dysfunction-associated steatohepatitis; CCl_4_, carbon tetrachloride; APAP, acetaminophen) (**Figure S1A; Table S1**). Using the Harmony algorithm [26], 84 samples comprising 86,072 HSC cells were merged into a reference atlas. The analysis effectively corrected for batch effects while preserving biological variation (**Figure S1B-S1C**). HSC identity was validated by high expression of canonical markers such as *Dcn* and *Des* (**Figure S2A**).

Unsupervised clustering revealed 11 transcriptionally distinct HSC subtypes, which were classified into qHSCs, proliferative HSCs (pHSCs), and injury-responsive HSCs based on their distribution across liver samples (**Figure 1B-1C and S2B-S2C**). Clusters 1-4 belonged to qHSCs, which reside in the normal liver and highly express quiescence-associated genes (e.g., *Lrat* and *Ets1*) as well as pathways related to lipid storage and homeostasis (**Figure 1D-1E; Table S2**). Clusters 5-6 were annotated as pHSCs, characterized by specific expression of proliferative markers and cell cycle signatures (**Figure 1D-1F**). Clusters 7-11 were designated as injury-responsive HSCs, which were enriched in post-injury samples **(Figure 1C)**. These HSCs comprised activated and inactivated HSCs, expressing classic activated (e.g., *Acta2* and *Col1a1*) and inactivated HSC markers (e.g., *Gsn* and *Gata6*; **Figure 1D**), along with enriched pathways like collagen biosynthesis and immune regulation **(Figure 1E**). Mapping annotations from the original studies [14, 16, 20, 24] supported the identities of quiescent, proliferative, and activated HSCs (**Figure S2D-S2E**).

Based on enriched functions and previously reported molecular features [1, 4, 7, 9], we further established a consistent nomenclature, categorizing the 11 subtypes into four functional states, qHSCs, pHSCs, aHSCs, and attenuated HSCs (atHSCs) (**Figure S3A; Table S3**). Three qHSC subtypes (central vein *Hhip*^high^, portal vein *Ngfr*^high^, and immune *Kdr*^high^ HSCs) display corresponding features, whereas the features of *Tonsl*^high^ qHSCs remain undefined. Two pHSC subtypes include S-phase *Top2a*^high^ and G2M-phase *Ccnb2*^high^ HSCs. Three aHSC subtypes are inflammatory *Ccl2*^high^, contractile-migratory *Acta2*^high^, and collagen-producing *S100a6*^high^ HSCs. We also found two atHSC subtypes (apoptosis-prone *Mrc2*^high^ and inactivated *Htra3*^high^ HSCs). The *Htra3*^high^ subtype was characterized by high expression of known inactivated HSC marker *Bambi* but low expression of quiescent HSC marker *Gfap*. However, *Mrc2*^high^ HSCs (cluster 10) was previously underappreciated, which highly expressed *Mrc2* (mannose receptor, C-type 2) and *Wt1* (WT1 transcription factor) (**Figure S3B**).

Interestingly, *Mrc2*^high^ HSCs showed enriched intrinsic apoptotic signaling pathway and increased activity of the p53 signaling pathway (**Figure 1E and 1G**), along with reduced cell cycle activity (**Figure 1F**). Immunofluorescence staining confirmed that Mrc2-positive or Wt1-positive HSCs highly co-expressed with p53 in injured mouse liver tissue (**Figure 1H**). These results supported that *Mrc2*^high^ HSCs were identified as an apoptosis-prone HSC subpopulation. Furthermore, this *Mrc2*^high^ subtype exhibited elevated expression of *Mmp* genes, which encode extracellular matrix-degrading enzymes, along with known suppressors of HSC activation or liver fibrosis, including *Wt1*, *Gata6*, *Nr4a1*, and *Nr4a2* [27–30] (**Figure S3C-S3D**). These observations collectively suggest a protective role against liver fibrosis.

To assess whether apoptosis-prone *Mrc2*^high^ HSCs are conserved between mouse and human, we performed integrative analysis of human snRNA-seq data (**Figure S4A-S4C**). A *MRC2*^high^ HSC subtype was identified in human (**Figure S4D-S4F**). *MRC2*^high^ HSCs also exhibited enrichment of apoptosis-related terms, p53 signaling pathway activation, and high expression of orthologous activation suppressors (*WT1*, *GATA6*, *NR4A1*, and *NR4A2*) (**Figure S4G-S4I**). Immunofluorescence staining of human liver tissue confirmed MRC2-positive or WT1-positive HSCs significantly co-expressed p53 (**Figure 1I**), paralleling the apoptosis-prone subpopulation observed in mouse liver. These findings collectively indicated a previously unrecognized, species-conserved apoptosis-prone *Mrc2*^high^ HSC subtype.

We further revealed distinct subtype distributions between acute and chronic injuries (**Figure S4J-S4K**). Acute injury was enriched in inflammatory *Ccl2*^high^ and contractile-migratory *Acta2*^high^ subtypes alongside proliferating and some quiescent (*Tonsl*^high^ and *Kdr*^high^) HSC populations. In contrast, the HSC landscape in chronic injury was enriched in *S100a6*^high^ and *Mrc2*^high^ subtypes, as well as *Ccnb2*^high^, *Acta2*^high^, and *Htra3*^high^ subtypes. These observations highlight distinct subtype compositions closely associated with different injury states.

A Quiescence-Activation-Attenuation Trajectory Orchestrates HSC Transitions *In Vivo* and is Recapitulated *In Vitro* Building on the above comprehensive HSC atlas, we next sought to explore whether the 11 subtypes represent distinct states along a continuous spectrum of quiescence, activation, and inactivation. To infer potential state transitions, we performed trajectory analysis using Dynamo[31] (**Figure 2A**), which was corroborated by an independent Monocle3 analysis [32] (**Figure 2B**). Notably, we identified a primary path, termed the quiescence-activation-attenuation (QAA) trajectory (Path-1), representing a sequential progression from qHSCs (*Hhip*^high^) through activated states (inflammatory *Ccl2*^high^, contractile-migratory *Acta2*^high^, and collagen-producing *S100a6*^high^) to an attenuation phase characterized by apoptosis-prone *Mrc2*^high^ and inactivated *Htra3*^high^ atHSCs (**Figure S5A**). This trajectory was accompanied by sequential expression of quiescent, activated, and inactivated HSC markers (**Figure S5B**). Notably, *Mrc2*^high^ atHSCs bridged the activated and inactivated states, suggesting a critical transitional role. The Path-2 trajectory represents an alternative path, in which HSC transition from proliferative G2M-phase *Ccnb2*^high^ pHSCs to *Acta2*^high^ aHSCs, and merging with the main QAA trajectory (**Figure 2A-2B**). Path-3 extends from *Ccnb2*^high^ pHSCs toward quiescent states, whereas Path-4 originates from *Tonsl*^high^ qHSCs toward central vein *Hhip*^high^ or portal vein *Ngfr*^high^ qHSCs. Collectively, these analyses underscore that *in vivo* HSC state transitions are multilineage rather than a simple binary switch between quiescence and activation.

**Figure 2.**
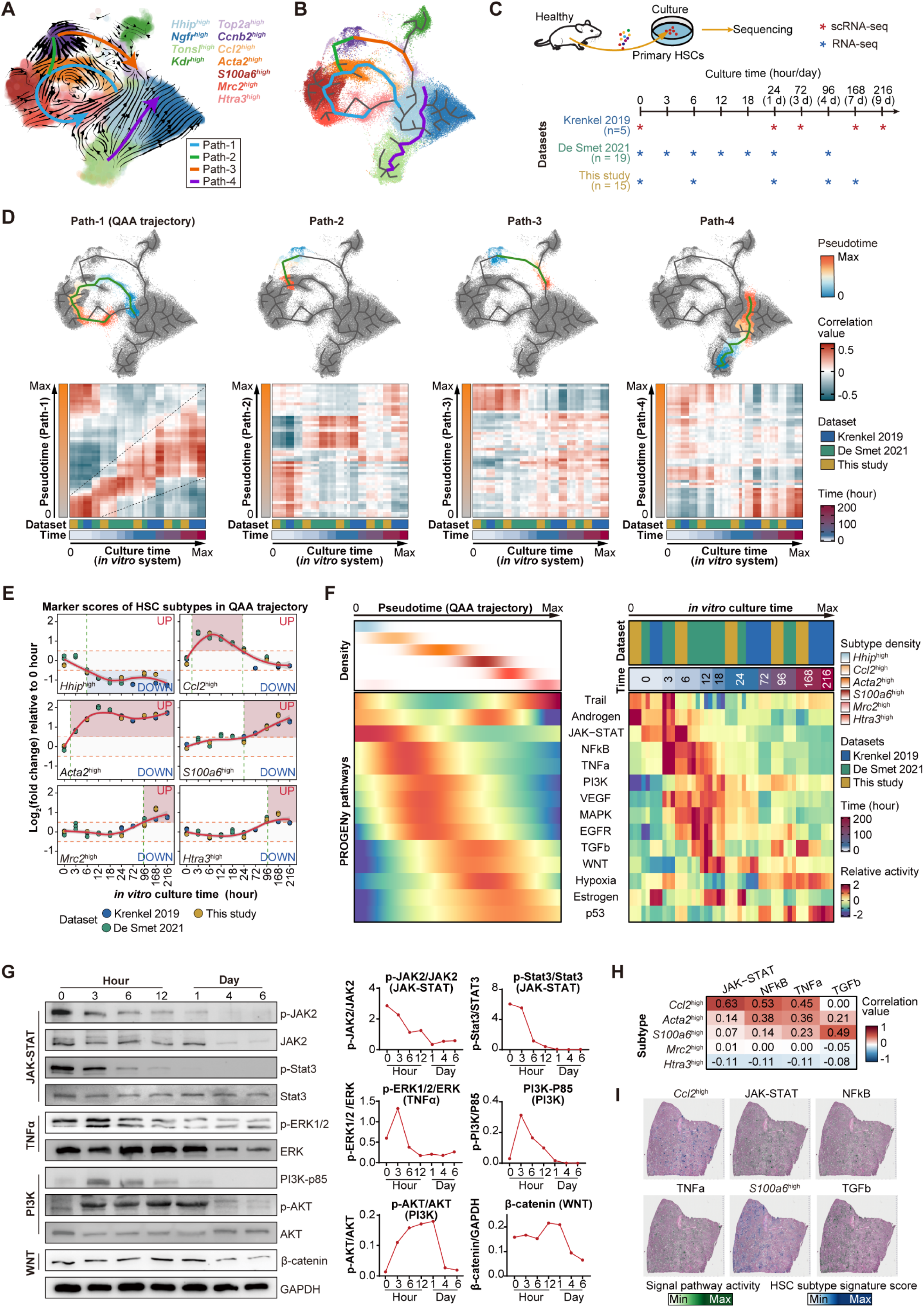
Reconstruction of HSC state transitions. **(A-B)** UMAP visualization of HSC state transitions. Four distinct paths (Path-1 to Path-4) are indicated, with Path-1 representing the quiescence-activation-attenuation (QAA) trajectory. Cell trajectories were inferred by Dynamo (A) and Monocle3 (B). **(C)** Experimental design for primary HSC activation *in vitro*. Three time-series independent datasets (this study and two publicly available datasets of Krenkel 2019 and De Smet 2021) were generated from cultured primary mouse HSCs. **(D)** Temporal transcriptome correlation analysis between *in vivo* pseudotime trajectories and *in vitro* HSC activation. Top panels: UMAP visualization of four paths colored by pseudotime progression. Bottom panels: Heatmap showing gene expression correlation between the *in vivo* trajectory and the *in vitro* HSC culture course. **(E)** Changes of average expression relative to 0 hours for the top 400 markers (ranked by fold change) in each HSC subtype along the QAA trajectory. Red and blue backgrounds were identified as periods of stimulation and repression, respectively. **(F)** Heatmap showing PROGENy-inferred pathway activities along pseudotime (*in vivo*) and culture time (*in vitro*). Top left: Cell density distribution of HSC subtypes across pseudotime. **(G)** Western blot (left) assay demonstrating the sequential activation of key proteins in JAK-STAT, TNFα/NFκB, PI3K and WNT, as well as their quantification (right) during primary HSC activation *in vitro*. **(H-I)** Spatial validation of relationships between pathways and subtypes in MASH mouse liver tissue (GSM5764420 from Guilliams 2022). **(H)** Correlation matrix between PROGENy-inferred pathway activities (JAK-STAT, NFκB, TNFα, and TGFb) and HSC subtype signature scores (*Ccl2*^high^ and *S100a6*^high^). **(I)** Spatial distribution of pathway activities and HSC subtype signatures in spRNA-seq data, demonstrating co-localization of specific pathways with HSC subtypes.

Next, we compared pseudotime-derived *in vivo* transcriptomes with our own RNA-seq data and two publicly available datasets, which comprised 39 time-series transcriptomic profiles (0-216 hours) from cultured primary mouse HSCs (**Figure 2C and S5C**). Remarkably, the QAA trajectory exhibited considerable temporal alignment with *in vitro* activation (**Figure 2D**), highlighting that the *in vivo* transcriptional dynamics along the QAA trajectory can be reflected by the *in vitro* model.

Further analysis of HSC subtype-specific gene expression changes confirmed that subpopulations along the QAA trajectory were sequentially upregulated during *in vitro* culture (**Figure 2E and S5D**). We observed that *Ccl2*^high^ and *Acta2*^high^ signatures expressed early (3 hours), followed by *S100a6*^high^ (24 hours), and later by *Mrc2*^high^ and *Htra3*^high^ signatures (96 hours). This temporal alignment suggests that the widely used *in vitro* model can serve as a comparable system for dissecting molecular mechanisms underlying the QAA trajectory.

Using PROGENy [33], we further revealed tightly coordinated signaling pathways along the QAA trajectory and *in vitro* activation (**Figure 2F**). As previously reported, JAK-STAT signaling was activated during the early phase following injury [34]. In the subsequent phase, TNFα/NFκB signaling peaked in *Ccl2*^high^ aHSCs. Fibrogenic pathways, including VEGF/EGFR and WNT/TGFβ signaling, were subsequently activated in contractile-migratory *Acta2*^high^ and collagen-producing *S100a6*^high^ aHSCs, respectively. Sequential activation of JAK-STAT, TNFα/NFκB, PI3K, and WNT pathways was confirmed by Western blot in time-series *in vitro* activation (**Figure 2G**). Notably, analysis of human primary HSCs exposed to TNFα or TGFβ[35, 36] suggests that TNFα preferentially promoted inflammatory features, whereas TGFβ enhanced contractile-migratory and collagen-producing programs (**Figure S6A-S6B**). Spatial transcriptomics of MASH mouse liver further confirmed that *Ccl2*^high^ and *S100a6*^high^ aHSC signatures co-localized with JAK-STAT/NFκB/TNFα and TGFβ signaling pathways, respectively (**Figure 2H-2I**). The attenuation phase was associated with p53 and Estrogen signaling pathways (**Figure 2F**), which were involved in promoting HSC apoptosis and suppressing HSC activation, respectively[37–39]. This stage-specific signaling cascade provides mechanistic insight into HSC transitions along the QAA trajectory. In summary, these analyses reconstruct a QAA trajectory that delineates the principal roadmap of *in vivo* HSC state transitions during liver injury. This trajectory is driven by the tight coordination of multiple signaling pathways and is notably preserved between *in vivo* and *in vitro* models.

The QAA Trajectory is Spatiotemporally Orchestrated and Supports Fibrosis-Free Repair in Acute Liver Injury After establishing the QAA trajectory computationally and revealing its preservation *in vitro* as inferred *in vivo*, we next investigated its functional features *in vivo*. We leveraged a time-series scRNA-seq dataset from an APAP-induced ALI model [19] (**Figure 3A**), which resolves spontaneously without fibrosis[8–12]. By analyzing changes in HSC subtype proportions (**Figure 3B**), marker gene expression (**Figure 3C**), and cell density distribution (**Figure 3D-3E**) in this temporal dataset, we revealed a coordinated and sequential activation along the QAA trajectory. Following injury, we observed a rapid expansion of activated HSCs. The inflammatory *Ccl2*^high^ and contractile-migratory *Acta2*^high^ aHSCs reached the highest fraction first, followed by a surge in collagen-producing *S100a6*^high^ aHSCs. This activation reached its peak around 48 hours post-injury, coinciding with a transient maximal expression of collagen genes at the same time point (**Figure S7A**). At the late stages of activation, an attenuation phase emerged, marked by the expansion of apoptosis-prone *Mrc2*^high^ and inactivated *Htra3*^high^ atHSCs (**Figure 3B-3E**), along with upregulation of canonical inactivated HSC markers (**Figure S7B**). At 168 hours post-injury, the HSC population had largely reverted to a quiescent state, indicating an effective and self-resolving response in ALI (**Figure 3B-3E**). We validated this cascade using independent temporal snRNA-seq datasets from TAA- or APAP-induced ALI in mice [24, 40, 41]. By mapping HSCs from these datasets to the mouse HSC atlas, we confirmed a consistent program underlying complete resolution in ALI (**Figure 3F-3G, S7C-S7G, and S8**). The sequential order of HSC subtypes in ALI, from early activation to late attenuation via apoptosis or inactivation, closely aligned with the QAA trajectory (**Figure 3H**). Interestingly, pHSCs transiently expanded at 36-48 hours, potentially contributing to the subsequent increase in qHSCs via the Path-3 trajectory (**Figure 3B-3D and S7F-S7G**).

**Figure 3.**
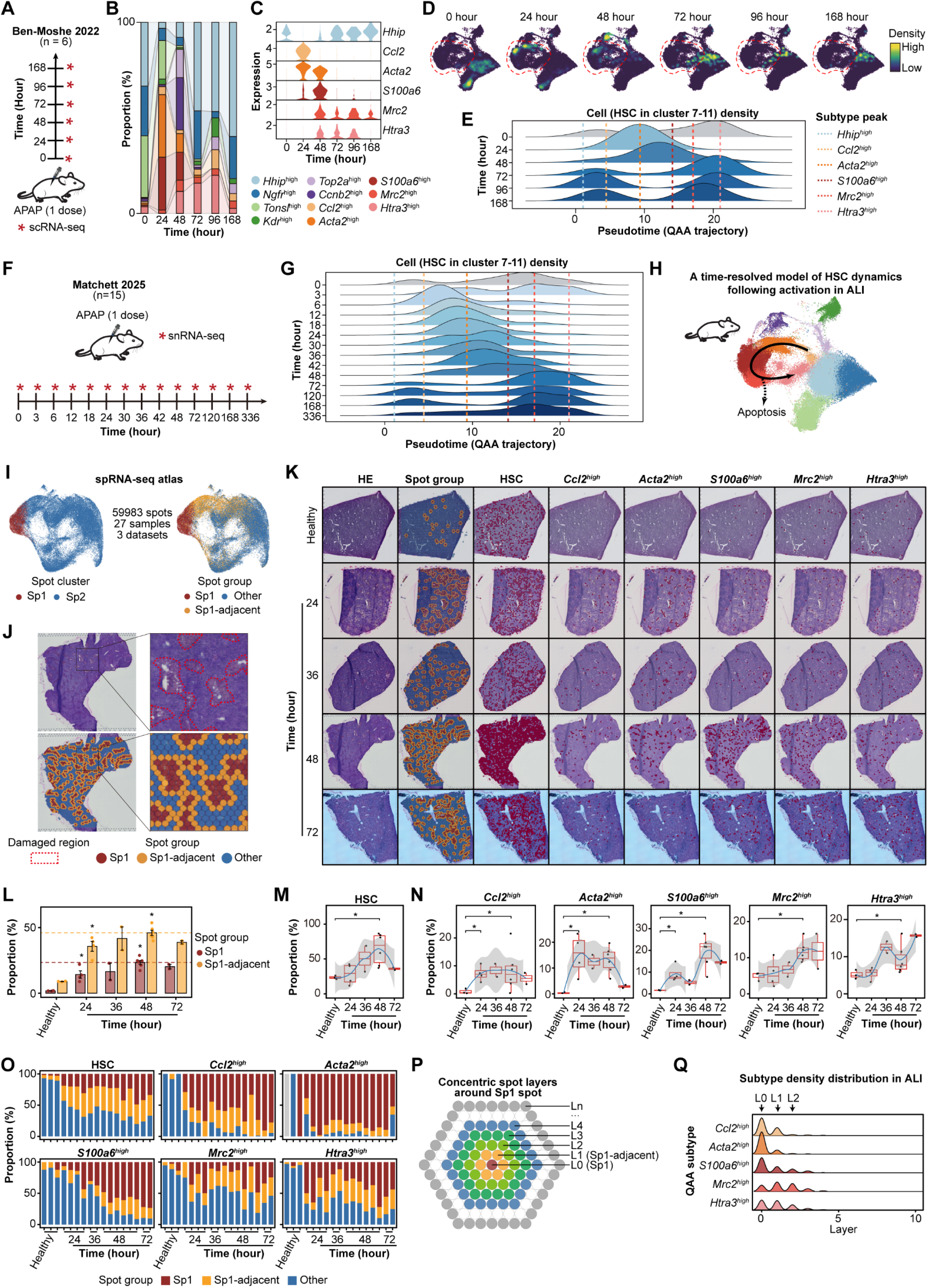
Spatiotemporal dynamics of HSC subtypes in response to acute liver injury. **(A)** Experimental design showing scRNA-seq data of APAP-induced acute liver injury (ALI) from Ben-Moshe 2022. **(B)** Temporal changes in HSC subtype composition across time points (0-168 hours). **(C)** Dynamic expression of marker genes (*Hhip*, *Ccl2*, *Acta2*, *S100a6*, *Mrc2*, and *Htra3*) in HSC subtypes following ALI. **(D)** UMAP density plots showing temporal changes in HSC distribution. **(E)** Cell density of HSCs in clusters 7-11 along the QAA trajectory. Dashed lines indicate peak density for each cluster. **(F)** Experimental design of snRNA-seq data measured during 0-168 hours post-APAP injection from Matchett 2025. **(G)** Cell density along the QAA trajectory confirming the temporal order of HCS activation. **(H)** Time-resolved model of HSC dynamics following ALI-induced activation. **(I-Q)** Independent validation using three spRNA-seq datasets from Guilliams 2022, Matchett 2025, and Ben-Moshe 2022. **(I)** Spatial transcriptomic atlas from integrated spRNA-seq analysis. Left: Identification of spot clusters (Sp1 and Sp2) in 27 samples from 3 datasets. Right: Classification of spots into three groups (Sp1: damaged spot; Sp1-adjacent: adjacent spot; Other: other spot) based on spatial proximity to Sp1 spots. **(J)** Representative HE staining and corresponding spatial distributions at 48-hours (GSM5764423 from Matchett 2025) after APAP-induced acute injury. Red dashed lines demarcate damaged regions. **(K-Q)** Spatiotemporal dynamics of HSC subtypes in ALI using spRNA-seq data from Matchett 2025 and Ben-Moshe 2022. **(K)** Representative spatiotemporal HE staining showing the spatial group, HSC-positive regions, and QAA subtype-positive regions in ALI. Red dots indicate positive spots. **(L)** Temporal changes in proportions of the two spatial groups. Statistical significance was evaluated using an unpaired one-tailed Wilcoxon test, comparing each time point with the 0-hour baseline within each spatial group (Sp1 and Sp1-adjacent). Error bars represent standard deviation. **(M)** Temporal changes in the proportion of HSC-positive spots among all spatial spots. **(N)** Dynamics of subtype-specific activation. Proportion of each subtype-positive spots among HSC-positive spots across time points. **(O)** Composition analysis of spatial groups within HSC-positive and subtype-positive spots across time points, revealing spatial localization of activated subtypes. **(P)** Schematics of multilayer concentric analysis. Layers (L0-Ln) were defined by increasing distance from Sp1-containing spots (L0). **(Q)** Density distribution of QAA subtype-positive spots across concentric layers, revealing spatial gradients of activation. Statistical significance denoted as: *, p < 0.05 (a two-sided Wilcoxon test was used except where specified).

To determine whether the spatial dynamics of HSC subtypes recapitulates the progression along the QAA trajectory, we analyzed three reported spRNA-seq datasets [19, 40, 42] (**Figure S9A**) and constructed a spRNA-seq atlas of 59,983 spots from ALI and CLI livers (**Figure 3I**). Through identifying damaged regions in HE staining and investigating marker expression, we first revealed damaged spots (Sp1) and adjacent spots (Sp1-adjacent) as well as HSC-positive spots (**Figure 3I-3K**). These three regions expanded following APAP treatment and peaked at 48 hours in ALI (**Figure 3K-3M and S9B**), indicating a transient activation of HSCs tightly coupled to injury area. Spatial mapping of HSC subtype-positive regions revealed a distinct spatiotemporally coupled distribution, which is consistent with the sequential activation of HSCs along the QAA trajectory (**Figure 3K and 3N-3O**). Specifically, HSC subtypes in the early QAA trajectory (*Ccl2*^high^, *Acta2*^high^, and *S100a6*^high^) were enriched in damaged and adjacent regions at 24 hours, with their positive regions sharply declining after 48 hours. In contrast, *Mrc2*^high^ and *Htra3*^high^ atHSCs emerged later (36-48 hours) and persisted or even expanded through 72 hours. These two atHSC subtypes exhibited broader distributions than other subtypes and extended beyond the adjacent injury region. To further delineate the spatiotemporal distribution of HSC subtypes, we developed a spatial multilayer concentric analysis that assigned damaged spots as centers and then constructed concentric spatial layers extending outward from each center (**Figure 3P**). Concentric analysis showed that all QAA subtypes were spatially constrained around the damaged region but displayed distinct radial distributions (**Figure 3Q and S9C**). Early subtypes were sharply concentrated in the innermost layers (L0) at 24 hours, whereas later subtypes (*Mrc2*^high^ and *Htra3*^high^) exhibited a broader distribution, spreading into outer layers (L1-L2) over time. These spatial transcriptomic analyses validate the QAA trajectory inferred from time-series scRNA-seq analysis, revealing a spatiotemporally coordinated cascade of HSC subtypes from activation to attenuation around damaged region.

Collectively, these findings underscore that the QAA trajectory represents a spatiotemporally coupled process *in vivo*. In ALI, HSCs undergo a QAA cascade from activation to attenuation around damaged region, which likely contributes to the efficient, fibrosis-free repair observed in ALI.

Chronic Liver Injury Disrupts the QAA Trajectory to Promote the Pathological Accumulation of Collagen-producing *S100a6*^high^ HSCs Given that the QAA trajectory effectively captured HSC spatiotemporal dynamics in ALI, we next examined whether this trajectory could also represent HSC dynamics in CLI, characterized by persistent fibrosis and imbalanced HSC subpopulations [20]. We analyzed HSC dynamics in CLI induced by repeated CCl₄ administration [20, 24] (**Figure 4A**). The inflammatory *Ccl2*^high^ and contractile-migratory *Acta2*^high^ aHSC subtypes prominently emerged during the early injury induced by a single dose of CCl₄ (**Figure 4B-4E**). With repeated injuries (4-19 dose CCl₄), the landscape pathologically shifted toward pronounced accumulation of collagen-producing *S100a6*^high^ aHSCs and increased collagen expression, together with an expansion of *Mrc2*^high^ atHSCs (**Figure 4B-4E and S10A**), leading to imbalanced HSC subpopulations [20]. Our immunofluorescence assays further confirmed the sequential emergence of *Acta2*^high^ and *Mrc2*^high^ HSC subtypes in CCl_4_-induced CLI model, consistent with their order along the QAA trajectory (**Figure 4F**). Conversely, the proportion of inactivated *Htra3*^high^ atHSCs and the expression of inactivated HSC markers remained modest increase (**Figure 4B and S10B**). These HSC dynamics, marked by the pathological accumulation of *S100a6*^high^ aHSCs and impaired progression toward *Htra3*^high^ atHSCs in CLI, were corroborated by independent datasets across diverse injury etiologies including chemical toxicity, metabolic dysfunction, and bile duct ligation (**Figure S10C-S10J and S11-S13**). These observations indicate that the QAA trajectory captures HSC dynamics in both ALI and CLI, thereby providing a unified model that makes the dynamics comparable across acute and chronic injury. The key difference lies in the HSC behavior along the QAA trajectory, where ALI shows transient activation with complete resolution, whereas CLI exhibits sustained activation and pathological accumulation of *S100a6*^high^ aHSCs.

**Figure 4.**
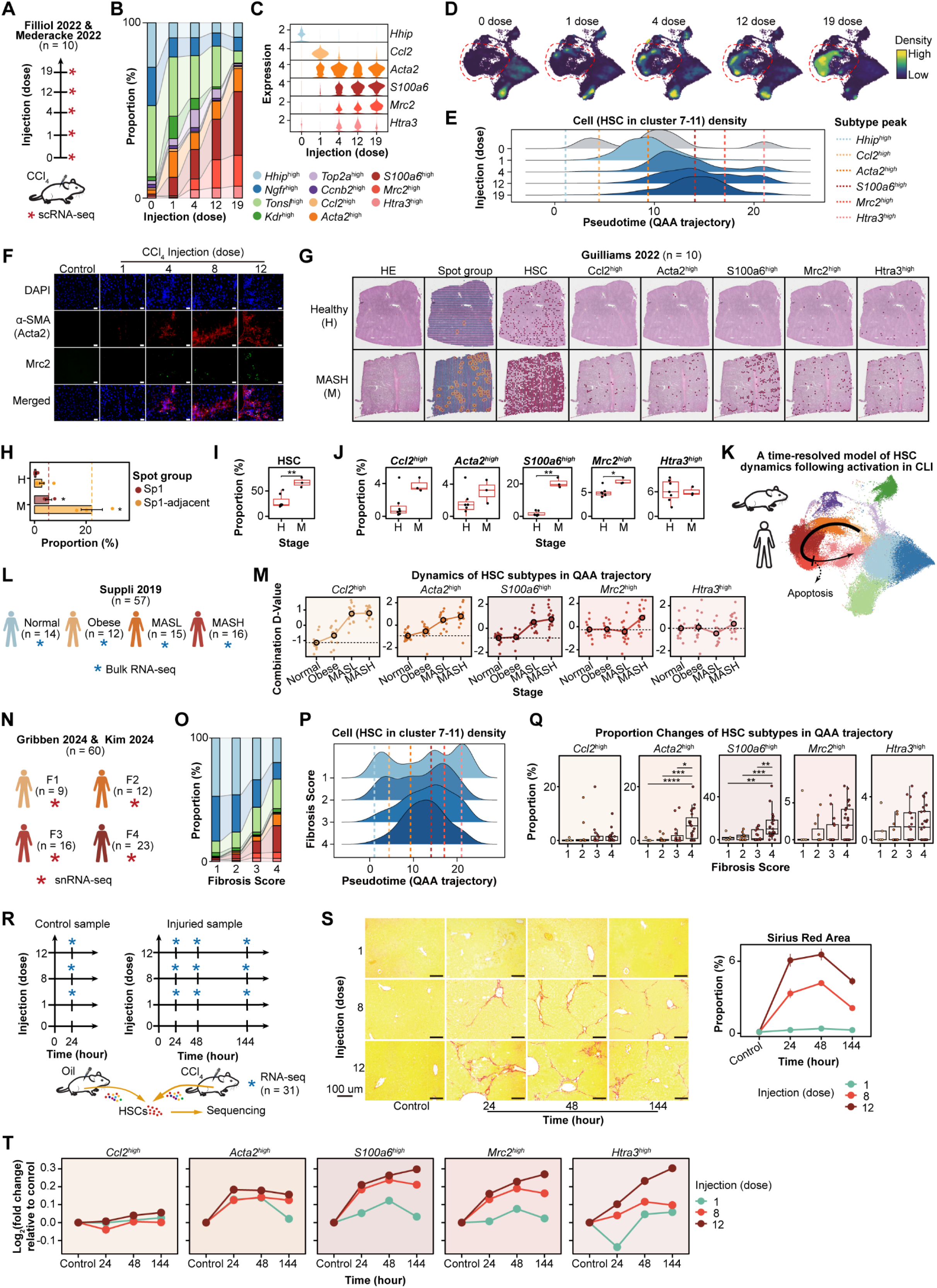
Chronic liver injury sustains profibrotic activation and prevents HSC inactivation. **(A-E)** Dynamic analysis of HSC subtypes during CCl₄-induced chronic liver injury (CLI) using scRNA-seq data from Filliol 2022 and Mederacke 2022. **(A)** Experimental design showing repeated CCl₄ injections (1-19 doses). 0 dose represents normal liver without any dose of CCl₄ injections. **(B)** Temporal changes in HSC subtype composition across injection numbers. **(C)** Dynamic expression of marker genes (*Hhip*, *Ccl2*, *Acta2*, *S100a6*, *Mrc2*, and *Htra3*) in HSC subtypes following CLI. **(D)** UMAP density plots showing HSC distribution at selected time points. **(E)** Cell density of HSCs in clusters 7-11 along the QAA trajectory. **(F)** Immunofluorescence of α-SMA (Acta2) and Mrc2 co-localization during the progression of CCl₄-induced injury. Representative images from control and different injections. Scale bar: 20 μm. **(G-J)** Spatiotemporal dynamics of HSC subtypes in CLI using spRNA-seq data from Guilliams 2022. **(G)** Spatial transcriptomic analysis of human MASH samples. HE staining and spatial distributions of the spot group, HSC-positive spots, and HSC subtype-positive spots are shown for healthy (GSM5764415) and MASH (GSM5764421) livers. **(H)** Proportions of the two spatial groups. Statistical significance was evaluated using an unpaired one-tailed Wilcoxon test, comparing each stage with the healthy baseline within each spatial group (Sp1 and Sp1-adjacent). **(I)** Proportion of HSC-positive spots among all spots. **(J)** Subtype-specific proportions among HSC-positive spots. **(K)** Schematic model illustrating HSC subpopulation dynamics during chronic liver injury. **(L-M)** HSC dynamics during MASH progression in human using bulk RNA-seq data from Suppli 2019. **(L)** Sample information showing normal, obese, MASL, and MASH groups. **(M)** D-value analysis tracking HSC subtype dynamics across disease stages. **(N-Q)** HSC dynamics in MASH patients stratified by fibrosis score using snRNA-seq from Gribben 2024 and Kim 2024. **(N)** Patient cohort information with fibrosis stages F1-F4. **(O)** Proportion of cell types across fibrosis stages. **(P)** Cell density along the QAA trajectory for different fibrosis scores. **(Q)** Quantification of HSC subtype proportions across fibrosis stages. **(R-T)** Temporal dynamics following withdrawal of fibrotic stimuli with analyzing our RNA-seq data. **(R)** Experimental design of time points for conducting RNA-seq in injury and recovery model. **(S)** Representative Sirius Red staining showing collagen deposition at different time points (left) and quantitative analysis of fibrotic area (right, n = 4). Error bars represent standard deviation. **(T)** Dynamic changes in HSC subtype marker expression following injury removal. Log₂ fold changes of subtype-specific markers (top 400, ranked by fold change) relative to corresponding controls at each time point and injection number. Scale bar: 100 μm. Statistical significance is denoted as: *, p < 0.05; **, p < 0.01; ***, p < 0.001 (a two-sided Wilcoxon test was used except where specified).

Spatial transcriptomic analysis of a diet-induced MASH model [42] provided independent support and further insight into this disrupted state. In MASH livers, damaged and adjacent regions were expanded, accompanied by accumulation of HSC-positive areas and *S100a6*^high^ and *Mrc2*^high^ subtypes, whereas *Htra3*^high^ atHSCs remained largely unchanged (**Figure 4G-4J**). Among these subtypes, *S100a6*^high^ aHSC signature was highly co-localized with regions of high collagen expression (**Figure S14**), suggesting their involvement in pathological matrix deposition. In contrast, *Mrc2*^high^ atHSCs, although expanded (**Figure 4J**), were spatially separated from collagen-rich regions (**Figure S14**), consistent with their potential protective role against liver fibrosis. Collectively, these data support a model in which recurrent injuries disrupt the QAA trajectory, pathologically shifting the balance toward persistent accumulation of collagen-producing *S100a6*^high^ aHSCs while simultaneously reducing the engagement of inactivated *Htra3*^high^ states, thereby exacerbating fibrosis and preventing resolution (**Figure 4K**).

To investigate the relevance of these findings to human disease, we applied a pseudo-bulk approach to bulk RNA-seq data from patients across the spectrum of fatty liver disease [43] (**Figure 4L**). The analysis displayed cell-type specificity (**Figure S15A**) and recapitulated changes in the liver cellular composition of cholangiocytes, macrophages, and Kupffer cells in line with a previous report [44] (**Figure S15B**). Remarkably, we observed that the activation sequence of HSC subtypes during disease progression followed the same order as the QAA trajectory (**Figure 4M**). Specifically, inflammatory *Ccl2*^high^ and contractile-migratory *Acta2*^high^ aHSCs expanded at the obesity stage, collagen-producing *S100a6*^high^ aHSCs were activated during MASL, followed by the emergence of *Mrc2*^high^ and a small population of *Htra3*^high^ atHSCs in MASH. This sequential order was independently validated by mapping human snRNA-seq HSCs across MAFLD progression onto the mouse HSC atlas (**Figure S15C-S15E**). The results showed that only a few inactivated *Htra3*^high^ atHSCs were detected in MASH, suggesting HSC attenuation in humans was incomplete. More importantly, alignment with fibrosis stages revealed that *Acta2*^high^ and *S100a6*^high^ aHSCs, unlike *Mrc2*^high^ and *Htra3*^high^ atHSCs, progressively expanded with fibrosis severity, obviously distinguishing F4 patients from earlier F1-F3 stages (**Figure 4N-4Q**). This cross-species conservation reinforces the clinical relevance of the disrupted QAA trajectory, highlighting *Acta2*^high^ and *S100a6*^high^ aHSCs as potential biomarkers of fibrosis severity.

Finally, we explored whether HSCs in severe CLI are capable of attenuation following injury withdrawal, as seen in ALI. Mouse models were established by administering different numbers of CCl₄ injections followed by recovery (**Figure 4R**), after which RNA-seq was performed on isolated primary HSCs. We found that the increased number of CCl₄ injections resulted in more severe liver fibrosis, with fibrotic areas persisting even after an extended recovery (144 hours post-injection; **Figure 4S**). Transcriptomic profiling revealed that repeated injuries substantially altered HSC dynamics. Although principal component analysis showed a transcriptomic shift toward a normal state during recovery (**Figure S16A**), matching previous reports [45], this response was insufficient to reverse fibrosis after 8 or 12 doses of CCl₄. Instead, aHSCs expanded at 24 hours post-injection and persisted throughout the 144-hour recovery period (**Figure 4T and S16B**). Independent scRNA-seq data from a mouse MASH regression mode [14] revealed that a substantial fraction (11.6%) of aHSCs persisted even 8 weeks after injury removal (**Figure S16C-S16F**). This contrasts with the transient surge of *Acta2*^+^ and *S100a6*^+^ aHSCs and the subsequent transition to *Mrc2*^+^ and *Htra3*^+^ atHSCs observed in one single dose of CCl_4_-induced ALI model (**Figure 4T**) and other ALI models (**Figure 3**). These findings indicate that, unlike ALI, severe CLI induces sustained dysfunction of the QAA trajectory after injury removal.

Together, these observations highlight that CLI disrupts the spatiotemporal dynamics of the QAA trajectory, driving the pathological accumulation of collagen-producing *S100a6*^high^ aHSCs and hindering the progression toward HSC inactivation. This persistent activation likely contributes to impaired fibrosis resolution and may underlie the limited reversibility of liver fibrosis, in stark contrast to the transient, self-limiting activation observed in ALI.

Stage-specific Transcriptional Regulators of HSC State Transitions

Next, we sought to elucidate the molecular regulatory mechanisms guiding the sequential state transitions along the QAA trajectory and to explore how these processes become disrupted in CLI. We constructed the global regulatory landscape across HSC subtypes using a connection specificity index (CSI) analysis, which revealed three distinct transcriptional modules with subtype-specific activities (**Figure 5A; Table S4**). Module 1 (M1) displayed strong internal expression correlations (**Figure 5A**) and showed high regulon activity and TF expression specifically in qHSCs (**Figure S17A-S17B**), being enriched for quiescence-related factors (e.g., *Lhx2*, *Nr1h4*, *Gata4*, and *Ets1*; **Figure 5A**) [46–49] and pathways including retinoic acid signaling (**Figure S17C**). In contrast, Module 3 (M3) was specific to activated HSCs (**Figure S17A-S17B**) and included pro-fibrotic regulators (e.g., *Runx2*, *Fos*, and *Prrx1*; **Figure 5A**) [50–54], associated with collagen production and inflammatory pathways. Module 2 (M2) defined a unique regulatory signature for *Kdr*^high^ qHSCs (**Figure S17A-S17B**), highlighting previously underappreciated heterogeneity within the quiescent HSC subpopulations. The mutually exclusive relationship between M1 and M3 was further supported by motif analysis of snATAC-seq data from normal and MASH livers [55] (**Figure 5B and S17D-S17G**). This analysis revealed a disease-stage-specific switch in activity, where M1 motifs were enriched in normal livers and M3 motifs predominated in MASH livers (**Figure 5C**), consistent with obvious regulatory differences between quiescent and activated HSCs reported previously [14].

**Figure 5.**
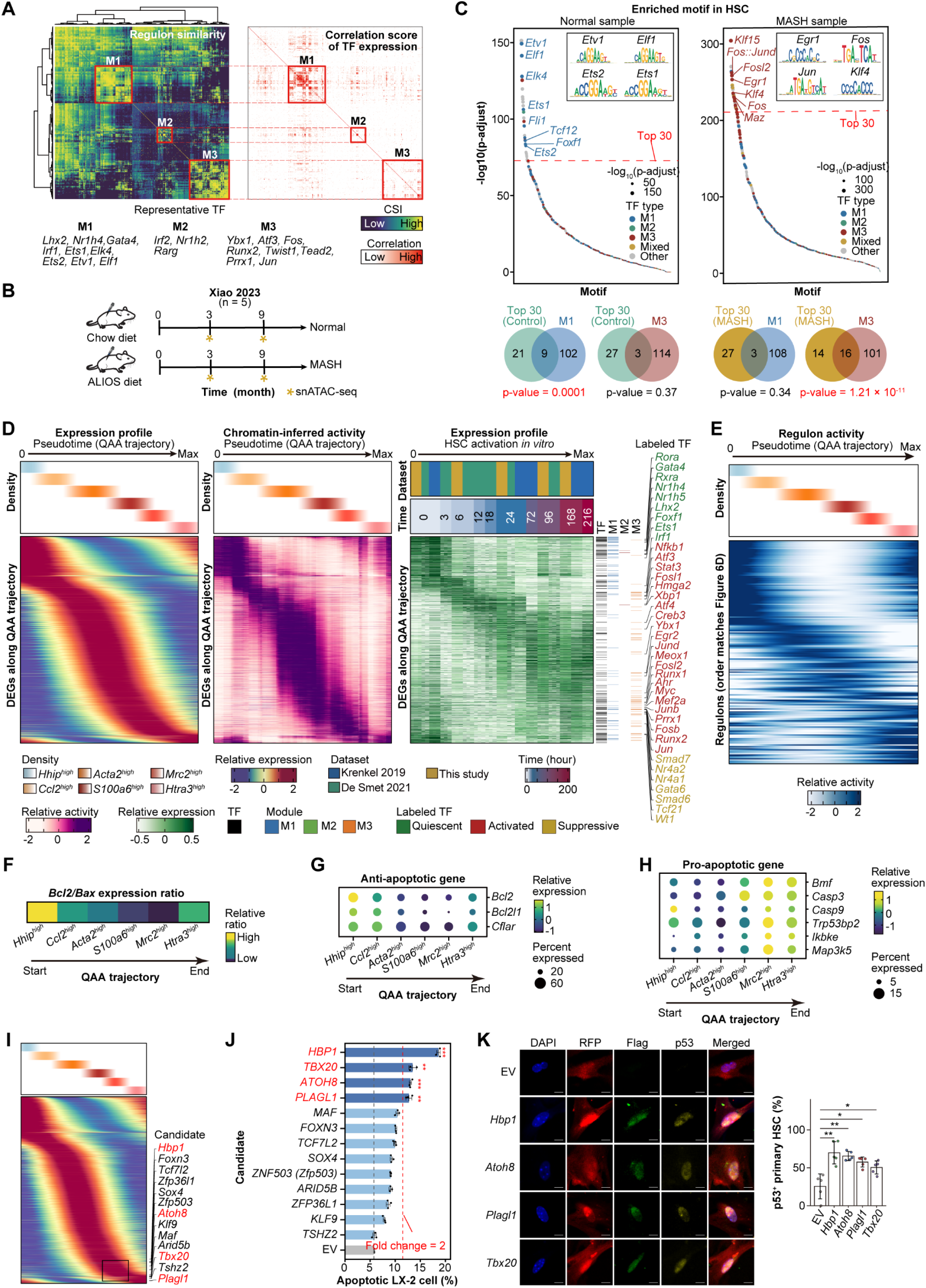
Regulatory analysis uncovers dynamic transcriptional regulations along the QAA trajectory. **(A)** Identification of transcriptional regulatory modules. Left: Heatmap showing similarity between regulons based on Connection Specificity Index (CSI) matrix, revealing three distinct modules (M1-M3). Right: Expression correlation matrix of transcription factors within the regulons using pseudo-bulk HSC transcriptomes. Representative TFs listed for each module. **(B)** Experimental design for mouse MASH model with snATAC-seq data from Xiao 2023. Timeline showing chow and ALIOS diet regimens at 0, 3, and 9 months. **(C)** Motif enrichment analysis in chromatin peaks of HSCs. Top: Volcano plots comparing motif enrichment between normal and MASH samples. TFs from top 30 enriched motifs (ranked by adjusted p-value) were intersected with M1 and M3 module TFs. Bottom: Venn diagrams showing overlap analysis. Statistical significance assessed by Fisher’s exact test. Representative motifs shown for M1 and M3 modules. **(D)** Integrated analysis of gene expression and chromatin accessibility dynamics. Left panels: Cell density along QAA trajectory and corresponding smoothed gene expression profiles. Middle panel: Dynamics of chromatin accessibility for matched genes along pseudotime. Right panel: Expression dynamics of TFs during *in vitro* HSC activation. **(E)** Regulon activity dynamics of TFs along the QAA trajectory. Heatmap showing activity scores for regulons corresponding to TFs with the same order in panel D. Missing values interpolated from adjacent TFs. **(F)** Bcl2/Bax expression ratio across HSC subtypes along the QAA trajectory, indicating apoptosis susceptibility. **(G)** Dot plot showing relative expression of anti-apoptotic genes across HSC subtypes. **(H)** Expression of pro-apoptotic genes along the trajectory. **(I)** Expression of candidate apoptosis-related TFs along the QAA trajectory. **(J)** Functional validation of TF-induced apoptosis. Quantification of apoptotic LX-2 cells following TF overexpression by flow cytometry. Red line indicates the threshold of 2-fold change. Data represent mean ± SD (n = 3). **(K)** Representative immunofluorescence images (left) and quantification (right, n = 5) showing that TF overexpression (Flag-tagged) in primary mouse HSCs (RFP-labeled) markedly increased p53 protein level. Data represent mean ± SD. Scale bars, 10 μm. Statistical significance is denoted as: *, p < 0.05; **, p < 0.01; ***, p < 0.001 (two-sided Wilcoxon test).

To investigate the transition from a binary switch to a continuous process, we analyzed the pseudotime dynamics of differentially expressed genes along the QAA trajectory identified by Monocle3 and highlighted key transcription factors in terms of expression, chromatin accessibility, and regulon activity (**Figure 5D-5E**). The early phase was dominated by quiescence-maintaining TFs from M1 (e.g., *Lhx2*, *Nr1h4*, *Gata4*) [46–49], followed by progressive engagement of pro-activation TFs from M3 (e.g., *Runx2*, *Fos*, and *Prrx1*) [50–54]. Concomitantly, pathway-specific TFs including *Stat2* in JAK-STAT, *Nfkb1* in NFκB, *Creb3* in PI3K, *Myc* in Wnt, and *Runx1* in TGFβ were sequentially activated and participated in inflammatory, contractile, and fibrogenic responses (**Figure S18A**). Notably, the attenuation phase was marked by the upregulation of HSC activation suppressors, including TGFβ pathway inhibitors (*Smad6*, *Smad7*, and *Nr4a1*) [29, 56, 57], and TFs associated with inactivation (*Gata6* and *Tcf21*) [46, 58, 59], and others (*Nr4a2* and *Wt1*)[27, 28, 30] (**Figure S18B**). This stage further reactivated some quiescence-related TFs (e.g., *Lhx2* and *Foxf1*) while retaining residual activation signals (e.g., *Runx1* and *Creb3*), representing a molecular program of state transitions (**Figure S18A and S18C**). This regulatory program might underpin the attenuation of activation features and the partial reacquisition of quiescence-related signatures during HSC inactivation. This regulatory cascade was remarkably reflected in our *in vitro* model alongside reactivation of partial quiescent HSC TFs during later stage (e.g., *Lhx2* and *Foxf1*) (**Figure 5D and S18D**).

Building on this regulatory landscape, we then examined the functions of HSC subpopulations, particularly focusing on the apoptosis-prone *Mrc2*^high^ subtype. Along the trajectory toward *Mrc2*^high^ atHSCs, the *Bcl2*/*Bax* (anti-apoptotic/pro-apoptotic gene) expression ratio [60] progressively declined, accompanied by an expression shift from survival genes (e.g., *Bcl2*) to pro-apoptotic effectors (e.g., *Bmf* and *Casp3*) [39, 61, 62] (**Figure 5F-5H**). Moreover, *Mrc2*^high^ atHSCs showed increased expression of *Nr4a2* and *Tcf21* (**Figure S18B**), both of which are known to promote HSC apoptosis [30, 59]. Interestingly, these apoptotic signals were partially attenuated during the transition toward inactivated *Htra3*^high^ HSCs, suggesting an apoptosis evasion as previously reported [10]. To functionally screen TFs implicated in this transition, we identified 13 TFs highly activated in *Mrc2*^high^ HSCs along the trajectory **(Figure 5I)**. Using lentiviral transduction, we overexpressed each of the 13 TFs in LX-2 cells **(Figure S19A)**. Flow cytometry analysis revealed that each of *HBP1*, *TBX20*, *ATOH8*, and *PLAGL1* significantly promoted the apoptosis of LX-2 cells, showing more than a two-fold change increase relative to the empty vector **(Figure 5J and S19B)**. Notably, none of these four TFs increased *COL1A1* or *COL3A1* expression, and *PLAGL1* even suppressed collagen levels (**Figure S19C**). All four TFs also showed strong correlation with p53 pathway activity **(Figure S19D)**. Furthermore, immunofluorescence assays in primary mouse HSCs confirmed that overexpression of each TFs significantly increased the proportion of p53-positive HSCs **(Figure 5K)**. Collectively, these results demonstrate that these four TFs promote apoptosis rather than fibrogenesis in HSCs. Notably, *Plgal1* and *Hbp1* have been reported to induce apoptosis separately in pancreatic β-cell and preadipocytes [63, 64]. Beyond the classical binary paradigm of quiescence and activation, this functional validation illustrates that the QAA trajectory facilitates the identification of stage-specific regulators. In summary, we delineated a sequential transcriptional cascade associated with HSC plasticity along the QAA trajectory, highlighting four TFs that promote HSC apoptosis.

Multicellular Microenvironment in Liver Injury Coordinated HSC Progression along the QAA Trajectory The transcriptional programs that contribute to HSC plasticity are thought to operate within a dynamic multicellular microenvironment. We hypothesized that the dysregulation of QAA trajectory in CLI arises from a combination of intrinsic HSC changes and impaired intercellular communication. To systematically assess how microenvironmental signals modulate HSC state, we constructed multicellular coordination networks in the liver using established approaches [65].

We integrated 11 mouse liver datasets measured by snRNA-seq, which more reliably preserves *in vivo* cell-type proportions [18]. A comprehensive atlas of 442,370 cells was constructed from 149 samples spanning 9 major cell types and 43 subtypes across ALI and CLI (**Figure 6A and S20-S21**). Analyzing co-occurrence of cell types using CoVarNet analysis [65], we identified four distinct cellular modules (CM01-CM04; **Figure 6B and S22A**), each representing coordinated multicellular programs with specialized functional profiles (**Figure 6C**). CM01 emerged as an inflammatory-fibrotic module enriched for inflammatory responses and collagen synthesis, comprising specific monocyte and macrophage populations alongside *Acta2*^high^ and *S100a6*^high^ aHSC subtypes from the QAA trajectory. CM02 comprised lymphocytes and Kupffer cells and was enriched for the activation of immune response. CM03, consisting of fibroblasts, hepatocytes, and HSCs, was linked to cell cycle and homeostasis. Notably, CM04 represented an HSC-centric module enriched for metabolic processes and uniquely contained attenuated HSCs (*Mrc2*^high^ and *Htra3*^high^).

**Figure 6.**
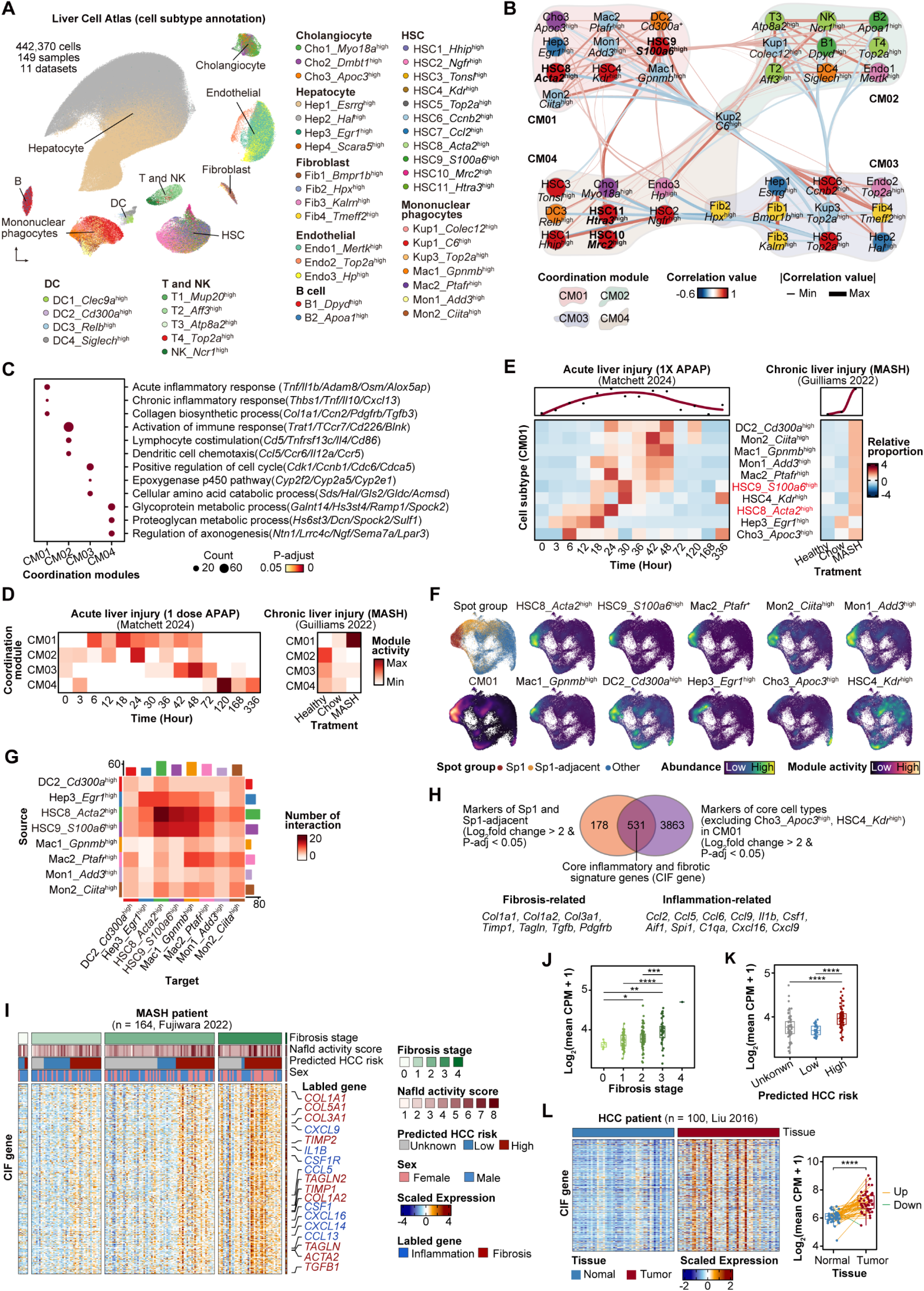
Liver multicellular coordination networks orchestrate HSC progression along the QAA trajectory. **(A)** Integrated liver cell atlas from 11 mouse snRNA-seq datasets. 43 cell subtypes were identified. Abbreviations: Cho (Cholangiocytes), Hep (Hepatocytes), DC (Dendritic cells), B (B cells), T (T cells), NK (Natural killer cells), Kup (Kupffer cells), Mon (Monocytes), Mac (Macrophages), Fib (Fibroblast), and Endo (Endothelial cells). **(B)** Cellular modules identified by CoVarNet analysis. Network visualization showing four distinct modules (CM01-CM04) based on cell type co-occurrence patterns. Edge thickness indicates correlation strength between cell subtypes. **(C)** Functional characterization of cellular modules. Dot plot showing enriched biological processes for each module. **(D)** Temporal dynamics of cellular modules during acute and chronic liver injury using snRNA-seq data from Matchett 2024 and Guilliams 2022. Heatmaps showing module activity across time points. **(E)** Cell subtype dynamics within CM01 during injury progression. Left: ALI response showing early Hep3_*Egr1*^+^ emergence followed by coordinated population recruitment. Right: CLI showing sustained cell populations. **(F)** Spatial distribution of CM01 activity and CM01-associated cell populations. Spot groups (Sp1, Sp1-adjacent, Other) showing enrichment for constituent cell types. **(G)** Intercellular communication network within CM01 module (excluding Cho3_*Apoc3*⁺ and HSC4_*Kdr*^+^). Heatmap depicting signaling interactions. Color represents the number of ligand-receptor pairs. **(H)** Identification of gene signature with core inflammation-fibrosis (CIF). Top: Spatial and cell type markers defining the 531-gene signature (log_2_ fold change > 2, adjusted p-value < 0.05). Bottom: Representative fibrosis and inflammation-related genes within the signature. **(I)** CIF signature expression in human MASH patients (Fujiwara 2022). Heatmap showing progressive increased expression of the signature with disease severity. **(J)** Quantification of CIF signature across fibrosis stages. Box plots showing significantly increased expression in advanced fibrosis (F3) versus early stages (F0-F2). **(K)** Quantification of CIF signature across hepatocellular carcinoma risks. Expression levels of CIF signature comparing patients with unknown, low, and high predicted HCC risk. **(L)** CIF signature expression in HCC tissues (Liu 2016). Heatmap showing gene expression in normal versus tumor tissues (left). Quantification demonstrating significant increase in tumor samples (right). Statistical significance is denoted as: *, p < 0.05; **, p < 0.01; ***, p < 0.001; ****, p < 0.0001 (two-sided Wilcoxon test).

Although the above analyses suggest the presence of multicellular coordination in both ALI and CLI, temporal analysis revealed markedly divergent module dynamics between these two types of injury models. In ALI, we observed a tightly orchestrated cascade, beginning with early activation of the inflammatory-fibrotic CM01 and immune-coordinating CM02 (3-48 hours), followed by CM03 engagement (30-72 hours), and culminating in activation of CM04 (72-336 hours) (**Figure 6D**). This temporal sequence closely mirrored the intrinsic HSC progression along the QAA trajectory. Detailed analysis of CM01 constituents in ALI showed the rapid emergence of *Egr1*⁺ hepatocytes within 3 hours post-injury, followed by sequential recruitment of other populations that remained active through 48 hours and declined by 72 hours (**Figure 6E**). In contrast, CLI exhibited CM01 activation with concomitant suppression of other modules (**Figure 6D**), which corresponded with the pathological accumulation of *S100a6*^high^ aHSCs (**Figure 6E**). Despite proportional increases in the *Mrc2*^high^ atHSC subpopulation in CLI (**Figure S22B**), CM04 activity remained insufficient for effective fibrosis resolution (**Figure 6D**). These observations suggest that CLI maintains a microenvironment persistently biased toward pro-fibrotic and pro-inflammatory states while failing to support fibrosis resolution. Spatial transcriptomic analysis provided architectural context for these functional modules. CM01 and most constituent populations (excluding *Apoc3*⁺ cholangiocytes and *Kdr*^high^ HSCs) were enriched in damaged and adjacent regions (**Figure 6F and S22C**), suggesting that a spatially organized pro-fibrotic and pro-inflammatory niche co-localize inflammatory and matrix-producing cells. Systematic prediction of intercellular communication identified potential bidirectional interactions between HSCs and monocytes/macrophages (**Figure 6G and S23A**). HSCs expressed monocyte-recruiting and educating factors (e.g., Postn, Il34, and Csf1), while monocytes and macrophages reciprocally produced pro-fibrotic signals (e.g., Tgfb1, Pdgfc, and Igf), suggesting the potential existence of self-reinforcing activation loops that could contribute to persistent fibrosis. Notably, *Egr1*⁺ hepatocytes emerged at the early time seem to produce signals to interact with both HSCs and monocytes/macrophages.

To assess clinical relevance, we developed an inflammation-fibrosis signature with 531 genes based on spatial and cellular markers derived from the CM01 module (**Figure 6H**). This signature, encompassing established fibrosis-related (e.g., *Col1a1*, *Col1a2*, and *Tgfb*) and inflammation-related (e.g., *Ccl2*, *Csf1*, and *Cxcl9*) genes, was significantly elevated in patients with MASH and biliary atresia liver disease compared to controls (**Figure S23B-S23C**). Expression levels progressively increased with fibrosis severity in a cohort of 164 MASH patients and were significantly higher in F3 compared with F0-F2 stages, highlighting its potential as a discriminative marker of fibrosis (**Figure 6I-6J**). Unexpectedly, the signature was associated with increased risk of hepatocellular carcinoma (**Figure 6K**) and showed elevated expression in tumor versus adjacent non-tumor tissue (**Figure 6L**), suggesting its potential utility as a prognostic biomarker bridging fibrosis and carcinogenesis.

These observations suggest that the QAA trajectory of HSC transitions is regulated within extensive multicellular networks. Coordinated modules appears to promote effective tissue restoration in ALI, while persistent CM01 activation and impaired engagement of other modules in CLI may promote self-sustaining pathological microenvironments that contribute to ongoing fibrosis and potentially elevate tumorigenic risk.

An Online Reference Map for Exploring HSC Heterogeneity

To support community-driven investigation of mouse HSC heterogeneity, we established an interactive resource featuring analytical flexibility via a Shiny-based tool (https://hscatlas.shinyapps.io/hsc_atlas_v1/). The full HSC atlas is openly accessible through Zenodo (https://zenodo.org/records/14855155).

## Discussion

Liver fibrosis is primarily driven by functionally heterogeneous HSCs, representing a major therapeutic challenge in chronic liver disease [1, 5–7]. Previous studies have largely examined ALI and CLI in isolation [14–20, 24, 25, 42, 66, 67]. Here, we compared fibrosis-free repair observed in ALI with persistent fibrogenesis in CLI to dissect the cellular and molecular mechanisms in HSCs that underlie fibrosis progression. Through integrated analysis of 86,072 single-cell HSC transcriptomes from 84 samples across seven studies, validated by independent multi-omics data from 991 human and mouse samples spanning acute and chronic liver injury models, we delineated distinct HSC responses in ALI and CLI. We established a cross-etiological and spatiotemporally resolved atlas to elucidate HSC dynamics, and suggests that pathological fibrosis in CLI is associated with the disrupted trajectory rather than solely with HSC overactivation.

A key finding of our analysis was the delineation of a continuous quiescence-activation-attenuation (QAA) trajectory, characterized by spatiotemporally cooperating processes *in vivo* and associated with sequential transcriptional and signaling cascades. Importantly, the QAA trajectory represents a shared trajectory for HSC dynamics after injury, enabling direct comparison between ALI and CLI, with injury persistence as the key differentiating factor. This shared trajectory allows us to leverage the spatiotemporally complete HSC resolution observed in ALI to elucidate the dysregulated accumulation of aHSCs in CLI. Applying this trajectory, we revealed that ALI proceeds through the full QAA trajectory to resolution, whereas CLI disrupts the trajectory, characterized by pronounced expansion of collagen-producing *S100a6*^high^ aHSCs and subtype imbalance[20]. Notably, this dysfunction persisted even after injury removal, with activated HSCs failing to resolve and fibrosis remaining, suggesting a potential mechanism underlying the limited reversibility in CLI [1, 68, 69]. These results suggest that fibrosis stems not only from sustained activation but also from impaired resolution programs, highlighting therapeutic opportunities to restore endogenous resolution pathways. Computationally, this supports the view that fibrosis-free repair in ALI provides a valuable and comparable reference for elucidating mechanisms of dysregulated fibrotic process in CLI. Evidence from a previous study[70] supports this notion, in which AAV-mediated overexpression of *Lhx2*, a key HSC quiescence-associated transcription factor re-expressed during fibrosis-free repair in ALI but suppressed in CLI, mitigated fibrosis in a chronic model.

We constructed multicellular coordination networks, revealing that diverse cell types organized into functional modules (CM01-CM04) to coordinate liver injury responses. These modules appear to execute coordinated biological programs, from promoting fibrosis to facilitating resolution. Their temporal alignment with the QAA trajectory suggests that HSC state is shaped by multicellular context. In ALI, sequential module activation parallels HSC transitions, likely providing microenvironmental cues that support complete resolution. In contrast, CLI exhibited disrupted dynamics, characterized by persistent CM01 activity coinciding with pathological *S100a6*^high^ HSC accumulation around damaged regions, while limited engagement of other modules. This pattern implies a programmatic dysfunction, where sustained CM01-driven inflammatory-fibrotic signaling may constrain HSCs in activated states.

The clinical relevance of our findings is strengthened by cross-species conservation. Analysis of human MASH progression revealed HSC transitional stages consistent with observations in mouse models. The abundance of *S100a6*^high^ and *Acta2*^high^ aHSCs correlated with fibrosis severity and effectively discriminated F3 and F4 from F0-F2 patients. Moreover, the inflammation-fibrosis signature genes derived from the CM01 module and damaged regions exhibited progressive enrichment with advancing fibrosis and were highly expressed in hepatocellular carcinoma, highlighting their potential as diagnostic or prognostic biomarkers. Notably, our finding of the coordinated activities of *S100a6*^high^ and *Acta2*^high^ aHSCs within CM01 aligns with the previous report that demonstrated opposing functions of HSC subpopulations in hepatocarcinogenesis [20]. Specifically, myofibroblastic HSCs, a collagen-producing state, were shown to promote tumor progression via secretion of pro-tumorigenic factors, whereas other subpopulations exerted protective effects[20]. This is consistent with our observation that the CM01 signature, covering *S100a6*^high^ and *Acta2*^high^ aHSCs, is not only correlated with fibrosis stage but also highly enriched in hepatocellular carcinoma. Together, these findings suggest that persistent CM01 activity may not only sustain fibrogenesis but also foster a tumor-permissive microenvironment, thereby highlighting the pathogenic and therapeutic relevance of specific HSCs and microenvironment imbalances.

Another key finding is the identification of a previously unrecognized, apoptosis-prone *Mrc2*^high^ atHSCs conserved in both mice and humans. Although, *Mrc2* was implicated in collagen clearance and fibrosis resolution [71], its association with an apoptosis-prone HSC population has not been reported. Here, we found that *Mrc2*^high^ atHSCs were characterized by reduced collagen production and increased sensitivity to apoptosis, displaying apoptosis-prone features alongside expression of anti-fibrotic genes (eg., *Gata6*, *Wt1*, *Tcf21*, *Nr4a1*, and *Nr4a2*)[27–30, 46, 57–59]. Immunofluorescence assays in mouse and human livers confirmed that *Mrc2*-positive HSCs significantly co-expressed with p53 protein. We also uncovered that *Mrc2*^high^ atHSCs were critical for characterizing the QAA trajectory, which may explain why previous studies failed to capture a continuous quiescence-activation-attenuation trajectory [14, 17, 22, 72]. Trajectory analysis revealed a pivotal position of *Mrc2*^high^ atHSCs between activated and inactivated states, with concomitant collagen downregulation that may signify an hyperactivated state reminiscent of T cell exhaustion observed during chronic infections and cancer [73]. Although situated upstream of inactivation, these cells likely constitute a state bifurcation point rather than directly transitioning toward inactivation. They may undergo either apoptosis or inactivation during fibrosis regression, consistent with previous reports on aHSC state transition following stimulus withdrawal [10, 11]. Notably, the elevated expression of *Tcf21*, implicated in both apoptosis and inactivation of HSCs[58, 59], together with *Gata6*, which is essential for HSC inactivation[46], underscores the molecular ambiguity and finely balanced nature of their state transition. While lineage tracing will be needed to definitively demonstrate the native HSC state transition *in vivo*[22, 74], our findings provide new insights into HSC biology, showing that activated HSCs during CLI are not uniformly pro-fibrotic.

Several limitations should be recognized. First, the contributions of qHSC subtypes in healthy livers (clusters 1-4) to fibrogenesis remain incompletely understood. Investigating their dynamics under homeostasis and during tissue repair could reveal diverse roles in maintaining hepatic equilibrium, orchestrating pro-fibrotic responses, and facilitating parenchymal regeneration. Second, this HSC atlas is currently based primarily on drug-induced and metabolic injury models. Incorporation of datasets from additional etiologies, such as viral hepatitis, will be required to establish a more comprehensive pan-etiology resource.

Overall, through analysis of spatiotemporally resolved HSC heterogeneity and dynamics in ALI and CLI, this study proposes that liver fibrogenesis involves dysfunction of the QAA trajectory, suggesting that pathological fibrosis arises partly from trajectory dysfunction rather than solely from HSC overactivation. Analogous to regeneration studies using developmental models as reference [75–78], comparative analysis of fibrosis-free repair in ALI offers a distinct perspective for understanding fibrosis in CLI. The analytical strategy established in this study is broadly applicable to other cell types and fibrotic processes in diverse organs, such as pancreatic fibrosis in chronic pancreatitis and intestinal fibrosis in inflammatory bowel disease, providing a methodological framework for investigating universal mechanisms of cross-organ fibrogenesis.

## Abbreviations

aHSCs: activated HSCs
ALI: acute liver injury
APAP: acetaminophen
atHSCs: attenuated HSCs
CCl4: carbon tetrachloride
CLI: chronic liver injury
CM: cellular module
CSI: connection specificity index
HSCs: hepatic stellate cells
iHSCs: inactivated HSCs
MAFLD: metabolic dysfunction-associated fatty liver disease
MASH: metabolic dysfunction-associated steatohepatitis
MASL: metabolic dysfunction-associated steatotic liver
pHSCs: proliferative HSCs
qHSCs: quiescent HSCs
scRNA-seq: single-cell RNA sequencing
snATAC-seq: single-nucleus ATAC sequencing
spRNA-seq: spatial RNA sequencing
TFs: transcription factors

## Acknowledgement

We thank the Biomedical Data Analytics and Supercomputing Core and the Analytical Instrumentation Core at the Guangzhou Institutes of Biomedicine and Health, Chinese Academy of Sciences, for their technical support. We gratefully acknowledge the researchers who generated and shared the publicly available datasets used in this study.

## Data availability statement

Bulk RNA-seq data measured in this study have been deposited in CRA023202 and CRA018107 of Genome Sequence Archive (GSA, https://ngdc.cncb.ac.cn/gsa/).

## Materials and methods

### Animal experimental protocol

A mouse model of liver injury was established through intraperitoneal administration of CCl_4_ (carbon tetrachloride). Experimental mice received either 1,4, 8, or 12 doses of CCl_4_ (1μl/g body weight, twice weekly), while control mice were injected with an equivalent volume of olive oil. The CCl_4_ solution was prepared by diluting in olive oil to achieve a final concentration of 20%. All animals were maintained under controlled conditions with a 12-hour light/dark cycle and provided with standard laboratory diet and water ad libitum. The experimental protocol was conducted in accordance with ethical guidelines and was approved by the Animal Ethics Committee of the Guangzhou Institutes of Biomedicine and Health, Chinese Academy of Sciences.

### Liver samples from clinical patients

Fresh liver tissue samples were obtained from patients with benign liver tumors (including hemangiomas) and from individuals with cirrhotic livers associated with liver cancer at the Eastern Hepatobiliary Surgery Hospital (Shanghai, China). Resected specimens were fixed and paraffin-embedded within 6 hours after surgical removal. Histological grading and staging of liver fibrosis were assessed according to the Ishak scoring system[79]. The study was approved by the Institutional Review Board of the Eastern Hepatobiliary Surgery Hospital, and written informed consent was obtained from all participants. All data were anonymized, and no patient-identifiable information was included.

### Preparation and staining of pathological sections

Liver tissue specimens were collected from the mid-portion of the largest lobe, fixed overnight in 4% paraformaldehyde (PFA), and processed into paraffin-embedded sections. Hematoxylin and eosin (H&E) staining and Sirius red staining were performed following established protocols from published literature [70]. For immunohistochemical and immunofluorescent analysis, routine procedures were conducted, including deparaffinization, endogenous peroxidase blocking, antigen retrieval, and sequential incubation with primary and secondary antibodies. The following primary antibodies were used and provided in Table S5: αSMA (ab124964, Abcam, 1:500 dilution), p53 (AF0879, Affinity, 1:100 dilution; human tissue), p53 (2524T, CST, 1:200 dilution; mouse tissue), Wt1 (12609-1-AP, Proteintech, 1:200 dilution), and Mrc2 (AF0564, Affinity, 1:200 dilution). For immunofluorescence co-staining of antibodies from the same species, a Tyramide Signal Amplification (TSA) kit was employed. The TSA method, as previously described [80, 81], utilizes horseradish peroxidase (HRP)-conjugated secondary antibodies to catalyze the deposition of fluorescently-labeled tyramide molecules onto target antigens in the presence of hydrogen peroxide (H_2_O_2_). This enzymatic reaction generates strong fluorescent signals directly bound to the antigen. Following microwave heat treatment, primary and secondary antibodies are removed, while the fluorescent label remains intact, allowing subsequent incubation with another antibody of the same species for dual staining.

### Isolation and culture of primary HSCs from mouse livers

Primary HSCs were isolated from both normal and injured mouse livers. The procedure began with intraperitoneal administration of anesthetics, followed by sequential perfusion of the liver via the portal vein using Hanks’ buffer containing EDTA and collagenase. All buffers and collagenase solutions were maintained at a constant temperature of 37°C throughout the process. After enzymatic digestion, liver cells were dissociated and subjected to two rounds of low-speed centrifugation (50g) to remove hepatocytes. The remaining supernatant, containing non-parenchymal cells, was collected and mixed with OptiPrep Density Gradient Medium (#1893, Serumwerk Bernburg AG) according to the manufacturer’s instructions. Density gradient centrifugation was then performed to isolate HSCs. For molecular analysis, HSCs were immediately preserved in Trizol to facilitate subsequent cDNA library construction and RNA sequencing. For cell culture experiments, isolated HSCs were resuspended in complete medium and maintained in a standard cell culture incubator. The culture medium was refreshed daily to ensure optimal growth conditions. Primary mouse HSCs were cultured in high-glucose DMEM (Gibco) supplemented with 10% fetal bovine serum (FBS, Gibco) and 1% penicillin-streptomycin.

### Western blot assay

Total protein was extracted from primary HSCs cultured *in vitro* at different time points through adding radio-immunoprecipitation assay (RIPA) lysis buffer with protease inhibitors. Protein bands were separated by 10% SDS-PEGA and transferred by the PVDF membrane. Using 5% BSA blocked nonspecific binding, and the primary antibody was incubated at 4^◦^C overnight. Details of the antibodies used in the study were provided in Table S5.

### Library preparation and RNA sequencing

Total RNA was extracted from cells using the Trizol reagent, followed by chloroform extraction and isopropyl alcohol purification. RNA sequencing (RNA-seq) libraries were constructed using the VAHTS Universal V8 RNA-seq Library Prep Kit for Illumina (NR605, Vazyme) according to the manufacturer’s instructions. The procedure involved the removal of genomic DNA (gDNA) using magnetic beads, followed by PCR amplification of the cDNA library. The amplified products were subsequently purified to ensure high-quality library preparation. The final RNA-seq library was sequenced on an Illumina HiSeq2500 platform, generating 150 bp paired-end reads.

### Cell lines and cell culture

Human hepatic stellate cells (LX-2) were maintained in Dulbecco’s Modified Eagle Medium (DMEM) supplemented with 2% fetal bovine serum (FBS) at 37°C in a humidified incubator with 5% CO₂. Cells were routinely tested and confirmed mycoplasma-negative using the MycoBlue Mycoplasma Detector (Vazyme).

### Lentiviral vector construction and stable cell line generation

The cDNA-encoding human transcription factors (FOXN3 [O00409-1], KLF9 [Q13886], TSHZ2 [Q9NRE2-1], ZFP36L1 [Q07352], MAF [O75444-2], ARID5B [Q14865-1], TCF7L2 [Q9NQB0-1], TBX20 [Q9UMR3], PLAGL1 [Q9UM63-1], HBP1 [O60381-1], ATOH8 [Q96SQ7-1], ZNF503 [Q96F45-1], and SOX4 [Q06945]) and mouse transcription factors (Hbp1 [Q8R316-1], Tbx20 [Q9ES03-1], Plagl1 [Q9JLQ4], and Atoh8 [Q99NA2]) were obtained from MiaoLingBio, China. Each cDNA was PCR-amplified and cloned into lentiviral expression vector using a restriction enzyme-based ligation. Human transcription factors cDNAs were insert into the pLenti-III-EF1α vector (Applied Biological Materials), whereas mouse transcription factor cDNAs were cloned into the pLenti-III-EF1α-Flag vector. Both vectors contain a puromycin resistance cassette, enabling antibiotic selection after transduction.

For lentivirus production, recombinant plasmids were co-transfected with the packaging plasmids psPAX2 and pMD2.G into HEK293T cells using polyethyleneimine (PEI, Yeasen). The medium was replaced with a fresh complete medium 8 hours post-transfection. Viral supernatants were collected at 36 and 60 hours post-medium replacement, centrifuged at 1000 × rpm for 5 minutes, and filtered through 0.45μm filters to remove cell debris. The viral stocks were stored at 4°C and used within one week.

For stable cell line generation, LX-2 cells were seeded in 6-well plates at 3×10^5^ cells per well. After 16-18 hours, cells were infected with viral supernatant in the presence of 8μg/mL polybrene (Merck). After 72 hours, 1μg/mL puromycin (InvivoGen) was added for selection. Stable transduced cells were used for subsequent experiments on day 8 post-transduction.

### Cell apoptosis assay

3×10^5^ cells were collected and processed using the Annexin V-FITC/PI Apoptosis Detection Kit (Vazyme), following the manufacturer’s instructions. Briefly, cell pellets were washed twice using ice-cold PBS and re-suspended in a 100μL binding buffer. Cells were incubated with 5μL Annexin V-FITC and 5μL PI in the dark at room temperature for 10 minutes, then a 400μL binding buffer was added to stop the reaction. Flow cytometry was performed immediately on a BD LSRFortessa (BD Biosciences). Control samples included unstained cells, cells stained with Annexin V-FITC alone, and cells stained with PI alone, for proper gating and compensation. Data analysis was conducted using the FlowJo software.

### Immunofluorescence experiments in mouse primary HSCs

RFP-tagged HSCs were isolated from *Lrat-cre-Rosa26-LSL-RFP* mice (purchased from GemPharmatech). Primary HSCs were cultured *in vitro* for 9 days to reach an activated state, transduced with lentivirus for 4 days. Cells were processed using standard immunofluorescence procedures, including fixation, permeabilization, blocking, and antibody incubation, for detection of rabbit anti-Flag (F7425, Sigma-Aldrich, 1:1000) and p53 (2524T, CST, 1:200).

### Quantitative real-time PCR (qPCR)

Total RNA was extracted with RNAiso Plus (Takara) according to the manufacturer’s protocol. RNA purity and concentration were determined spectrophotometrically using NanoPhotometer (Implen). For cDNA synthesis, 1μg RNA was reverse-transcribed using PrimeScript RT Master Mix (Takara) in 20μL reactions. qPCR was performed in triplicates using TB Green Premix Ex Taq II (Takara) on a CFX96 Touch Real-Time PCR Detection System (Bio-Rad). Expression levels of target genes (*FOXN3*, *KLF9*, *TSHZ2*, *ZFP36L1*, *MAF*, *ARID5B*, *TCF7L2*, *TBX20*, *PLAGL1*, *HBP1*, *ATOH8*, *ZNF503*, and *SOX4*) were normalized to *β-actin*. Relative gene expression was calculated using the ΔΔCt method, with the empty vector (EV) group serving as the reference. Primer sequences are provided in Table S6.

### Integration of public mouse single-cell RNA-seq datasets and downstream analysis

To minimize potential batch effects introduced by technological differences, we collected publicly available single-cell RNA sequencing (scRNA-seq) datasets generated using the 10X Genomics platform. Seven processed datasets covering four etiologies were downloaded, with the detailed accession information provided in Table S1. Data processing and analysis were performed using the Seurat R package (v5.0.1) [82]. For quality control, doublets were identified and removed using scDblFinder (v1.14.0)[83], while potential ambient RNA contamination was detected with decontX (v1.4.0) [84, 85] and subsequently refined using ddqcR (v0.1.0) [86]. HSCs, as defined in the original papers, were extracted from the integrated dataset of all liver cell types for downstream analysis. Count matrices from individual datasets were merged into a single file, followed by uniform data filtering. To address discrepancies in gene numbers across datasets, only shared genes were retained. Cell cycle scores were calculated using the CellCycleScoring function. The merged dataset was normalized (NormalizeData function, method = ‘LogNormalize’, scale.factor = 10,000) and scaled to regress out the S score, G2M score, and the percentage of mitochondrial gene expression. Principal component analysis (PCA) was performed using 3,500 highly variable genes. The Harmony R package (v1.2.0) was employed to correct potential batch effects arising from differences in studies, samples, and replicates [26]. The top 40 Harmony dimensions were used as input for Uniform Manifold Approximation and Projection (UMAP), with clustering performed at a resolution of 0.44. Ambiguous clusters were identified and excluded by validating the expression of established HSC markers, including *Dcn*, *Des*, *Ecm1*, and *Reln*. Differentially expressed genes (DEGs) were identified using the Seurat function FindAllMarkers with a Wilcoxon rank sum test, and multiple testing correction was applied using the Bonferroni method. DEGs with an absolute log2 fold change > 0.25 and adjusted p-value < 0.05 were retained for further analysis. To further select marker genes specific to each cluster, pairwise differential expression analysis was performed between each cluster and all other clusters using the FindMarkers function. Only genes with log2 fold change > 0 were retained, and the intersection between this result and the previous marker gene list was used to ensure that the final marker genes are those most highly expressed in the target cluster compared to all others. The ComplexHeatmap R package (v2.15.4) [87] was employed to visualize gene expression, signaling pathway activity, and regulon activity along pseudotime.

### Function enrichment analysis

Enrichment analyses for upregulated DEGs or maker genes in each HSC subtypes were conducted using the enrichGO function from clusterProfiler R package v4.4.4 [88], with the parameters set to adjusted p-value < 0.05 and count > 2. For enrichment analysis in downstream genes of regulon, count > 3 was used. Gene sets were retrieved from the msigdbr R package v7.5.1 [89].

### Prediction of signaling pathway activities using PROGENy

Signaling pathway activities in HSCs were inferred using the PROGENy R package (v1.10.0) [33]. The analysis was performed with the ‘top’ parameter set to 500 for mouse and human data, while all other parameters remained at their default settings. The PROGENy scores for cells within each HSC subtype were averaged for visualization.

### Gene regulatory network analysis

To identify gene regulatory networks (regulons) activated in each HSC subtype, we employed the SCENIC software (v0.12.1) [90], using the raw count matrix as the input. The analysis involved three main steps: (1) Construction of a regulatory network using GRNBoost2; (2) Identification of regulons utilizing RcisTarget; (3) Calculation of regulon activity for each cell type via AUCell. To assess the similarity between regulons, the connection specificity index (CSI) was computed for distinct regulons using the calculate_csi function from the scFunctions R package (v0.0.0.9000).

### Distribution of HSCs across injury stages or time points

To analyze the distribution of HSC subtypes across different injury stages or time points, we calculated the proportion of each subtype relative to the total HSC population in different pathological samples. The spatial distribution of HSC subtypes was visualized using density plots on UMAP embeddings, generated with the Plot_Density_Custom function from the scCustomize package (v1.1.1). To quantify the enrichment of HSC subtypes in specific injury stages (normal, acute injury, chronic injury), we computed the ratio of observed to expected cell numbers (R_o/e_) for each subtype using the calTissueDist function from the Startrac R package (v0.1.0) [91]. Expected cell numbers were derived from chi-square tests. A subset was considered enriched at a particular injury stage if its R_o/e_ value was greater than 1.

### Trajectory inference and pseudotime reconstruction

Trajectory analysis of HSCs was performed using Dynamo [31] and Monocle3 [32]. For Dynamo analysis, the Python (v3.8) dynamo library (v1.2.0) was employed, following the guidelines outlined at https://dynamo-release.readthedocs.io/. Due to the unavailability of fastq files for five datasets, this analysis was restricted to two datasets with accessible fastq data: GSE137720 [13] and E-MTAB-8263 [24]. The fastq files were processed using Cell Ranger (v7.0.0) [92] with the mm10 reference genome (downloaded from https://www.10xgenomics.com) and used to generate BAM files. Unspliced and spliced raw count matrices were derived using the velocyto.R package (v0.6) and subsequently processed with the recipe_monocle function in Dynamo. For dimensionality reduction, the top 3,000 highly variable genes were identified and projected onto a UMAP embedding using default parameters. RNA velocity vectors were estimated from the normalized matrix and visualized in the spatial context.

Trajectory analysis of HSCs was further conducted using the Monocle3 package (v1.3.4) [32]. The processed Seurat object was converted into a Monocle3 object, and clustering was performed based on the UMAP dimensionality reduction obtained from Seurat. The trajectory structure was constructed using the learn_graph and order_cells functions. To infer the pseudotime of the four inferred trajectories, the choose_graph_segments function was employed to isolate the branch extending from the start point to the end point in each trajectory. The pseudotime was then calculated using the order_cells function, with the start point set as the root in each trajectory. Then, differentially expressed genes were inferred by the graph_test function for each trajectory.

### Cross-validation by using public scRNA-seq and snRNA-seq data in mice

For cross-validating HSC dynamics in ALI, the publicly available matrices from snRNA-seq datasets GSE280652 and GSE223558 [40, 41] were downloaded. For cross-validating HSC dynamics upon CLI, publicly available count matrices from scRNA-seq data were downloaded from independent datasets (GSE176042, GSE189600, GSE145086, GSE200366, GSE171904, and GSE148339) [16, 17, 24, 45, 55, 66, 93, 94]. Then, the data were processed by using the Seurat package (v5.0.1). For quality control, singlets, identified by scDblFinder (v1.18.0) [83], were retained and further processed by ddqcR (v0.1.0) [86] in each sample, while potential ambient RNA contamination was detected with decontX (v1.4.0). Then, all samples were merged. The same functions used in the above analysis were applied to calculate cell cycle scores, normalize and scale the merged dataset, and perform PCA and batch effect correction. For identifying HSCs, gene expression of known markers for various cell types were assessed, including fibroblasts (*Clec3b*, *Cd34*), cholangiocytes (*Anxa4*, *Cftr*), hepatocytes (*Alb*, *Cps1*, *Asgr1*), endothelial cells (*Stab2*, *Kdr*), lymphocytes (*Cd2*, *Cd247*), macrophages (*Marco*, *Cd163*), B cells (*Mzb1*), T/NK cells (*Nkg7*, *Themis*), and HSCs (*Dcn*, *Ecm1*, *Reln*, *Des*). Then, the identified HSCs were mapped to mouse HSC atlas to explore cell dynamics.

### Cross-validation by using publicly available spatial transcriptomes in mice

Publicly available spatial transcriptomes were collected from GSE192742 [42] and processed using the Seurat package (v5.0.1). Raw spatial transcriptomics objects were filtered by retaining genes expressed in at least one spot. Each sample was individually normalized using the SCTransform method, preserving the complete transcriptome for integration. For integration, principal component analysis (PCA) was conducted per sample. 3,000 shared features were selected using SelectIntegrationFeatures, and the objects were prepared for SCTransform-based integration via PrepSCTIntegration. Integration anchors were identified using reciprocal PCA (RPCA) reduction with the FindIntegrationAnchors function with ‘dims = 1:30’. The final integrated object was generated using IntegrateData with the same dimensional settings. Following integration, PCA was recalculated on the integrated assay, followed by the UMAP dimensionality reduction (RunUMAP, dims = 1:20). Nearest neighbors and unsupervised clustering were performed using FindNeighbors and FindClusters. UMAP plots and spatial visualizations were generated to assess cluster distributions and sample contributions. To identify the injury-associated cluster, Euclidean distances were computed based on tissue coordinates using GetTissueCoordinates. For each sample, the nearest five spots to each injury-associated cluster spot were identified, and an adaptive threshold was used to annotate neighboring spots as adjacent regions.

To investigate the spatial dynamics of HSC subtypes, the data was normalized using the NormalizeData function. For each sample, HSC-positive spots were identified based on the co-expression of the canonical mesenchymal marker *Vim* and the HSC marker *Dcn*, given that HSCs are mesenchymal in origin. The analysis was restricted to these HSC-positive spots.

Top 20 marker genes in each HSC subtypes were curated. For each cluster, a module score was calculated using the AddModuleScore function across all HSC-positive spots. In parallel, a collagen signature (using *Col1a1*, *Col1a2*, *Col3a1*, *Col5a1*, and *Col5a2*) was also profiled. Additionally, pathway activity was inferred using PROGENy (top 500 responsive genes, mouse model) for each spatial transcriptomics dataset.

To assess HSC subtype-positive spots, we implemented an adaptive thresholding strategy in each sample. Specifically, the Otsu method from the R package autothresholdr was applied to linearly rescaled module scores (0-255), and final thresholds were set as:

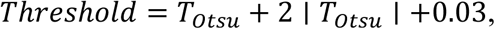

where 𝑇*_otsu_* is the rescaled Otsu threshold. Spots exceeding the adjusted threshold were defined as positive for the corresponding HSC subtype. The spatial distribution of each subtype was visualized using SpatialPlot. To assess temporal and pathological dynamics, we calculated the proportion of subtype-positive spots relative to the total number of HSC-positive spots per sample.

Spatial multilayer concentric analysis for damaged regions

To investigate the spatial organization of HSC subpopulations around damaged regions, we implemented a multilayer concentric analysis for spatial transcriptomics data. Specifically, we recognized the central injury-associated core spots (L0) based on prior cluster annotation (St1) from the integrated spatial dataset. From each L0 spot, concentric spatial layers (R1, R2, …) were iteratively constructed by identifying neighboring spots based on Euclidean distance in image coordinates.

For each L0 spot, adjacent spots were defined by computing pairwise Euclidean distances and selecting those within a dynamic distance threshold (mean of the 2nd to 6th nearest neighbor distances + 10 units). Candidate spots for each subsequent layer were determined from the unvisited neighbors of the previous layer. This expansion was iterated up to 50 layers or until no further spots were reachable.

After generating all concentric layers for each L0 spot, we assigned each Visium spot its minimum concentric layer index, corresponding to the shortest path from any L0 spot. Spots not captured by any expansion path were assigned the maximum observed layer within the sample.

Subsequently, for each cluster, we quantified the proportion of positive spots across spatial layers and visualized them using stacked bar plots and ridge density plots to assess changes of spatial regions.

Cross-validation by using publicly available snATAC-seq data in mice

Publicly available single-nucleus ATAC-seq (snATAC-seq) data, represented as fragment files, were obtained from the Gene Expression Omnibus (GEO) under accession number GSE189600 [55]. Data processing was performed using the R packages signac (version 1.14.0) and Seurat [95]. Peaks for each sample were identified using the CallPeaks function and subsequently merged to generate a unified peak set. Quality control metrics, including the transcription start site (TSS) enrichment score (TSS.enrichment), nucleosome signal (nucleosome_signal), and blacklist region ratio (blacklist_ratio), were computed using the TSSEnrichment, NucleosomeSignal, and FractionCountsInRegion functions, respectively. The following quality thresholds were applied to filter cells: 5,000 < nCount_peaks (counts of peaks) < 100,000, blacklist_ratio < 0.05, nucleosome_signal < 4, and TSS.enrichment > 4. To remove potential doublets, the scDblFinder function from the scDblFinder R package (version 1.18.0) [83] was employed. Dimension reduction was performed using singular value decomposition (SVD) on a subset of 3,000 highly variable features. Batch effects arising from sample variability were corrected using the *Harmony* R package (version 1.2.0) [26]. The corrected Harmony dimensions (2-30) were used as the input for UMAP visualization, and clustering was performed at a resolution of 0.1. Gene activity scores were calculated using the GeneActivity function. To identify HSCs, we assessed gene activities of known markers for various cell types. The known markers used here are the same as those applied in the preceding analysis of gene expression.

To identify enriched peaks specific to HSCs in control and MASH samples, snATAC-seq data from HSCs were isolated. Differentially enriched peaks between control and MASH conditions were identified using the FindMarkers function. Additionally, enriched motifs within these condition-specific peaks were determined using the FindMotifs function. To infer chromatin accessibility dynamics of QAA DEGs along the QAA trajectory, HSCs from the snATAC-seq dataset were mapped onto the mouse HSC atlas based on gene activity scores and further analyzed using the computational pipeline detailed in the Methods section to characterize dynamic features of clusters 7-11 along this trajectory.

Cross-validation by using time-series transcriptome profiles of HSCs from *in vitro* activation models

To investigate HSC dynamics during *in vitro* activation, three datasets were analyzed: a bulk RNA-seq dataset generated in this study, a bulk RNA-seq dataset from GSE173920 [45], and a scRNA-seq dataset from GSE132662 [96]. For the bulk RNA-seq data in this study, preprocessing was performed using Fastp (v0.21.0) [97] to remove adapter sequences and low-quality reads. Clean reads were aligned to the GRCm38 genome (retrieved from https://www.gencodegenes.org/mouse/release_M25.html) using HISAT2 v2.0.4 [64]. Gene counts and normalized expression values (TPM, Transcripts Per Kilobase Million) were generated using StringTie (v2.1.4) [98]. For the GSE173920 dataset, processed gene counts were downloaded and normalized to TPM. For the GSE132662 scRNA-seq dataset, processed counts from HSC cells were retrieved. To enable joint analysis with the bulk RNA-seq datasets, HSC cells from the same time points were randomly selected and assigned into three pseudobulk datasets, which were subsequently normalized to TPM.

The three TPM matrices were merged, and batch effects were corrected using the limma package (v3.60.4) [99]. To analyze HSC dynamics over time, the average log2 fold change of the top 400 upregulated DEGs in each HSC subtype relative to the 0-hour time point was calculated for each sample. Smoothed log2 fold change trajectories over time were used to quantify subtype-specific dynamics. For each HSC subtype, time segments with smoothed values greater than 0.5 or less than -0.5 were identified as periods of stimulation or repression, respectively.

Comparative pseudo-bulk correlation analysis between *in vivo* trajectories and *in vitro* HSC activation

To investigate the correspondence between HSC state transitions upon distinct *in vivo* trajectories and *in vitro* activation dynamics, we performed a pseudo-bulk correlation analysis comparing single-cell trajectory data with time-resolved bulk RNA-seq profiles. For each inferred trajectory (Path1-Path4), we utilized pseudotime calculated by Monocle3 and divided all single cells into 50 equally sized intervals. For each interval, the pseudo-bulk expression of each gene was calculated as the mean log-normalized expression across all cells within the interval. This yielded a smoothed gene expression matrix (50 intervals × genes) per trajectory, representing the temporal transcriptional dynamics along the *in vivo* transition. TPM matrices of *in vitro* cultured HSCs were preprocessed by log-transformation. Samples were ordered by time. To focus on informative genes, the top 3,000 genes with the highest standard deviation across pseudotime intervals in each trajectory were selected. The intersection of these genes with those in the bulk dataset was retained for downstream analysis. Both the trajectory-based and *in vitro* matrices were independently z-score-normalized by gene before calculating Pearson correlation coefficients between pseudotime intervals (columns) and bulk samples (columns). Correlation heatmaps were generated using the ComplexHeatmap package in R. Each heatmap (Path1-Path4) was scaled to a common color range (Pearson r from -0.5 to 0.5) and combined into a multi-panel figure for comparative visualization.

### Cross-validation by using publicly available snRNA-Seq datasets in humans

We compiled publicly available snRNA-seq datasets generated using the 10X platform, including from GSE174748 [20], GSE185477 [18], GSE202379 [100], GSE212046 [20], GSE212837 [21], GSE243981 [101], GSE223581 [40], and GSE244832 [22]. Because processed data was not available for GSE244832, GSE223581, and GSE243981, we applied a similar pipeline as the one used for processing mouse HSC scRNA-seq data to identify HSCs. HSCs in all datasets were then identified by their expression of established markers (*DCN, DES, RELN,* and *ECM1*).

Next, we integrated HSCs initially annotated across all 8 datasets. To improve HSC identification, we reprocessed the merged dataset using a modified pipeline: PCA was performed with 4000 highly variable genes, UMAP utilized the top 30 Harmony dimensions, and clustering was conducted at a resolution of 0.05. Contaminating cell clusters (e.g., macrophages, and other non-HSC populations) were removed based on their distinct marker expression (Fibroblast: *CLEC3B, CD34*; Cholangiocyte: *ANXA4, CFTR*; Hepatocyte: *ALB, CPS1, ASGR1*; Endothelial: *STAB2, KDR*; Lymphocyte: *CD247, CD2*; Macrophage: *CD163, MARCO*; B: *MZB1, IGKC*; T/NK: *NKG7, THEMIS*). HSCs identified through these two steps were retained for downstream analyses and mouse atlas mapping. For data integration in refined HSCs, we employed a similar pipeline to mouse HSC integration with adjustments: PCA used 2500 highly variable genes, UMAP was performed based on the top 35 Harmony dimensions.

### Mouse RNA-seq analysis for exploring HSC dynamics after stimulus withdrawal *in vivo*

Similar steps were used for processing bulk RNA-seq data generated in this study. After obtaining the normalized TPM expression values, batch effects were corrected using the limma package (v3.60.4) [99], with Log_10_(TPM + 0.1) as the input. To infer the dynamics of HSC subtypes in response to injury, public and local datasets were analyzed separately. For each HSC subtype, the top 400 upregulated DEGs (sorted by average log_2_ fold change) were selected. Average expression changes (log_2_ fold changes) of these genes for CCl_4_-injected samples at different time points relative to corresponding control samples were calculated. This approach allowed us to assess subtype-specific transcriptional changes over time.

Exploring HSC dynamics in RNA-seq of liver tissues using a pseudo-bulk analytical strategy

To investigate HSC dynamics in bulk RNA-seq data, we employed a pseudo-bulk analytical strategy modified from a published method [102], where the coordinated expression of a marker gene set derived from single-cell RNA-seq data was utilized to analysis the dynamics of specific cell populations in bulk RNA-seq samples. Pseudobulk data was generated from the scRNA-seq dataset at Zenodo.6035873. [19], which was also used in the integrative analysis. This dataset encompasses major liver cell types (**Figure S5A**), including HSCs with refined cell subtype annotations. To construct pseudobulk samples, 1000 random samplings of 3000 cells were performed for each sample. Reads from each sampling were pooled to create a bulk sample, which was then normalized to log_10_(CPM + 1). This process yielded 3000 pseudobulk samples, along with proportion information for each cell type.

Next, we aimed to identify combinations of genes whose expression values highly correlated with the proportion of specific cell types. The following steps were implemented: (1) Gene sorting: Genes were ranked based on their correlation with the proportion of each cell type; (2) D-value calculation: For each cell type, the difference (D-value) between the mean expression of the top N genes and the bottom M genes was computed for every pseudobulk sample; (3) Scaling: D-values for each cell type were scaled across all pseudobulk samples to obtain scaled D-values; (4) Correlation assessment: The correlation between scaled D-values and cell type proportions was calculated; (5) Iterative optimization: Steps 2–4 were repeated, iterating over N (2 < N < 40) and M (40 < M < 50), to identify gene combinations with the highest correlation between scaled D-values and proportions. The gene combinations with good performance were used to infer the relative proportions of specific cell types in new bulk RNA-seq datasets.

To evaluate the dynamics of cell types during MAFLD development in humans, raw expression counts were downloaded from GSE126848 [44]. Counts were normalized to log_10_ (CPM + 1). The gene combinations identified above were converted to their human orthologs using the convert_mouse_to_human_symbols function in the nichenetr package (v1.1.1) [103]. D-values for each cell type were calculated across these bulk RNA-seq samples to infer cell type dynamics during MAFLD progression.

### Mapping independent scRNA-seq and snRNA-seq cells to mouse HSC atlas

The HSCs identified for cross-validation were mapped to the developed mouse HSC atlas using the following approach adjusted from a previous study [104]. Pearson’s correlation coefficients were calculated between the new cells and atlas cells based on a gene set covering the top 400 marker genes in each HSC subtype. This approach ensured equal representation of all HSC subtypes by incorporating the most discriminative genes specific to each subtype, preventing potential bias toward subtypes with larger marker gene repertoires. To mitigate technical differences between datasets, only genes present in both the new dataset and the atlas were included in the correlation analysis. This gene selection strategy helped minimize the impact of platform-specific gene detection differences and enhanced the robustness of cell type assignment across datasets with potentially varying gene compositions. For integrating human HSCs into the mouse HSC atlas, human gene symbols were converted to their mouse orthologs using the convert_human_to_mouse_symbols function from the nichenetr package v1.1.1 [103]. For each new cell, we assigned its identity based on a two-tiered analysis of k-nearest neighbor classification (KNN) from mouse atlas cells: firstly, prioritize the most frequent cell type among the top 5 correlated mouse atlas cells, and then determine the final cell type using the sum of correlation values.

### Dynamic analysis of cells in clusters 7-11 along the QAA trajectory

To systematically characterize HSC dynamics along the QAA trajectory across biological contexts from integrated and independent datasets, cells from aHSC and atHSC clusters (clusters 7-11) were computationally isolated. We performed trajectory alignment by calculating Pearson correlation coefficients between query cell transcriptomes and pseudotime-ordered target cells along the QAA trajectory from Monocle3. This enabled the transfer of pseudotemporal coordinates from maximally correlated reference cells to individual query cells. The obtained cell-state distributions across biological conditions were subsequently visualized through kernel density estimation ridge plots, quantitatively delineating context-dependent shifts of HSC.

### Analysis of human bulk RNA-seq data to assess TNFA and TGFB effects in HSCs

To evaluate the impact of TNFA and TGFB on HSCs, two bulk RNA-seq datasets were analyzed: GSE22806 [35] and GSE148849 [36]. Raw expression matrices from GSE22806 and GSE148849 were normalized using the approach reported in the original studies. Genes associated with HSC features, including inflammatory, contractile-migratory, extracellular matrix-producing, and inactivated, were identified and visualized. Differences in expression between control and treatment groups were statistically assessed using the Wilcoxon test.

### Construction of liver cell atlas

To investigate intercellular coordination through shifts in cell proportions, we employed snRNA-seq, which more reliably preserves the true liver cellular composition [18], to develop a liver cell atlas. A total of 11 publicly available raw snRNA-seq FASTQ datasets (GSE148339, GSE200366, GSE211018, GSE223558, PRJEB56116, GSE184506, GSE189600, GSE192740, GSE225381, GSE254121, and GSE268112)[40, 42, 55, 66, 94, 105–110] generated on the 10x Genomics platform were collected to minimize batch effects arising from technical variation. Raw sequencing reads were processed with CellRanger (v7.0.0) using the mm10 reference genome, and the expression matrices obtained were analyzed with Seurat (v5.0.1). For quality control, doublets were removed with scDblFinder (v1.14.0), and potential ambient RNA contamination was detected with decontX (v1.4.0) [84, 85] and further refined with ddqcR (v0.1.0). Count matrices from all datasets were merged, retaining only genes shared across studies. To reduce memory usage, the merged matrices were processed in Python using Scanpy (v1.10.3)[111], retaining only protein-coding genes for downstream analysis. Gene expression data were log-normalized (scale factor = 10,000) and scaled while regressing out cell cycle scores (S and G2/M phases), mitochondrial gene percentage, and total RNA counts. Highly variable genes were identified in a batch-aware manner, excluding genes associated with cell cycle, mitochondrial, and ribosomal functions. The top 3,000 highly variable genes were retained for downstream analysis. Principal component analysis (PCA) was performed on scaled data, followed by batch correction using Harmony (harmonypy, v0.0.10) to integrate samples across studies and replicates. Neighborhood graphs were constructed from the Harmony-corrected embeddings, and UMAP was applied for visualization. Unsupervised clustering was performed using the Leiden algorithm with a resolution of 2.0. Cell cluster annotation was first conducted using CellTypist (v1.7.0)[112, 113] with the Healthy_Mouse_Liver model, and further validated by examining canonical marker genes for major liver cell types, including mononuclear phagocytes (*Cd14, Lyz2, Cd68, Vsig4*), dendritic cells (*Flt3, Xcr1, Cd209a, Siglech*), T and NK cells (*Bcl11b, Cd3d, Klra4, Ncr1*), B cells (*Cd79a, Cd19, Cd22, Mzb1*), hepatocytes (*Cyp2e1, Asgr1, Cps1*), cholangiocytes (*Anxa4, Sox9, Epcam*), endothelial cells (*Stab2, Kdr, Mecom, Efnb2*), fibroblasts (*Clec3b, Cd34*), and hepatic stellate cells (*Dcn, Ecm1, Reln, Des*). Each major cell type was subsequently reanalyzed using the same pipeline to resolve finer subtypes. Marker genes for each cell subtype were subsequently identified using the FindAllMarkers function in Seurat. This workflow generated a comprehensive liver cell atlas with subtype-level annotations, enabling downstream analyses of cellular composition and intercellular coordination.

### Identification of coordination module

Identification of multicellular coordination modules was performed using the R package CoVarNet (v0.3.0)[65], based on the mouse liver cell atlas constructed from the aforementioned snRNA-seq datasets. The analytical workflow proceeded as follows. First, the raw frequency matrix was calculated. For each sample, the number of cells belonging to each annotated cell subtype was counted, converted into relative frequencies (i.e., the proportion of each subtype within the total cells of that sample), and assembled into a cross-sample frequency matrix. To systematically compare cell coordination patterns between ALI and CLI, we incorporated datasets from both conditions, including GSE223558, GSE189600, GSE200366, GSE192740, and GSE225381 [40, 42, 55, 66, 108]. Next, frequency normalization was performed using the freq_normalize function in CoVarNet to apply Z-score scaling within each dataset, thereby reducing batch effects and enhancing cross-study comparability. Subsequently, cellular modules were identified by applying non-negative matrix factorization (NMF) using the nsNMF algorithm on the normalized frequency matrix. The decomposition was conducted with k ranging from 2 to 30 iterations (nrun = 30). Based on consensus analysis, k = 4 was selected as the optimal rank. Representative cell subtypes of each module were extracted from the basic matrix, selecting those with weights > 0.05 and among the top 10 ranked subtypes as the defining members of each module. Finally, cell correlation networks were constructed using the pair_correlation function to calculate Pearson correlation coefficients between cell subtypes, identifying cell pairs exhibiting cooperative (positive) or antagonistic (negative) frequency changes. Pairs with |r| > 0.2 were retained to build the intercellular network, which was then visualized in Cytoscape [114] to delineate the characteristic multicellular coordination modules under different injury contexts.

### Functional enrichment analysis of multicellular coordination modules

To characterize the potential biological functions of each cellular module, the following analytical procedure was performed. First, module-specific marker gene sets were derived by collecting the marker genes of all cell subtypes included in each module. Within each module, marker genes of the constituent subtypes were ranked in descending order based on their average log₂ fold change (avg_log2FC), and the top 100 significantly upregulated genes (defined as avg_log2FC > 0.25 and FDR < 0.05) were selected and merged to form the module marker gene set. Subsequently, functional enrichment analysis was conducted, with only enrichment terms having count > 4 retained.

### Spatial localization analysis of multicellular coordination modules

To investigate the spatial distribution of multicellular coordination modules, the following analytical workflow was implemented. First, reference data were collected from the mouse liver cell atlas. Marker genes were identified for each cell subtype using the thresholds average log₂ fold change > 0.5 and adjusted p-value < 0.05. The top 40 marker genes from each subtype were selected to form the candidate gene set. RNA count matrices, cell subtype labels, and UMI counts were then extracted from the Seurat object to generate the reference object required by RCTD. Next, spRNA-seq data were prepared by reading and integrating multiple 10x Visium datasets. Tissue coordinates and spatial gene expression matrices were extracted to construct the SpatialRNA object for RCTD. Cell-type deconvolution was then performed using the spacexr package (v2.2.1)[115]. The create.RCTD and run.RCTD functions were applied with key parameters set as doublet_mode = “doublet”, gene_cutoff = 1.25e-7, and fc_cutoff = 0.25. After completion, the cell-type weight matrix for each spatial spot was obtained and normalized to relative proportions for subsequent visualization. Finally, spatial projection of cellular module activity was performed to further interpret the spatial organization of cellular coordination. The module matrix (W, K = 4) derived from NMF decomposition was projected onto the spRNA-seq data. The RCTD weight matrix was first mean-centered and normalized across spots, with negative values truncated to zero. The processed weights were then matched to the trained NMF basis matrix to compute a module activity score for each spatial location, thereby mapping the spatial activity patterns of multicellular coordination modules.

### Cell-cell communication analysis of multicellular coordination modules

To explore the intercellular signaling mechanisms underlying multicellular coordination modules, the following analytical workflow was applied. First, data preparation was performed using the integrated liver snRNA-seq atlas. To ensure computational efficiency and minimize sampling bias, no more than 5,000 cells per subtype were randomly selected based on cell subtype annotations. The target cell populations associated with the CM01 module (excluding Cho3_*Apoc3*⁺ and HSC4_*Mrc1*^high^) were extracted as the focus of the analysis. A CellChat object was then constructed from these cells, with cell subtypes designated as grouping labels. Subsequently, cell-cell communication analysis was conducted using the CellChat R package (v1.6.1)[116] to infer ligand-receptor-mediated interactions. The createCellChat function was used to generate CellChat objects from Seurat data, followed by subsetData to retain expressed ligand-receptor pairs. The CellChatDB.mouse database was applied as the reference ligand-receptor framework. Interactions were inferred using the computeCommunProb function, integrated into signaling pathway-level networks via computeCommunProbPathway, and finally aggregated using aggregateNet. The resulting communication networks were visualized to depict intercellular signaling patterns within the cellular module.

### Identification of core inflammation-fibrosis signature genes and their clinical relevance

To identify a core set of inflammation-fibrosis signature genes, we employed the following strategy. First, using the FindMarkers function from the Seurat R package, spatial spots from the damaged region (St1) and its adjacent regions were compared with spots from other regions to identify marker genes (avg_log2FC > 2 and FDR < 0.05). Next, integrating the module analysis, we selected subpopulations enriched in the CM01 module within St1 and its adjacent regions, excluding Cho3_*Apoc3*^+^ and HSC4_*Mrc1*^+^, which were not aggregated in these areas, and extracted their marker genes (avg_log2FC > 2 and FDR < 0.05) to form a gene set. The intersection of these two gene sets was then defined as the core inflammation-fibrosis signature gene set.

To assess the clinical relevance of this core gene set, we further obtained multiple liver disease-related RNA-seq datasets (GSE77314, GSE122340, GSE162694, and GSE193066)[117–120]. Raw count data were normalized to TPM and integrated with corresponding clinical information. Expression differences of the core inflammation-fibrosis signature genes across patient stratifications were visualized using heatmaps and boxplots, allowing evaluation of their association with clinical features

### Statistical analysis

In this study, statistical analyses were conducted in R platform (3.9.1/4.4.1) and included Wilcoxon test and t-test as described in the figure legends.

## Declaration of generative AI and AI-assisted technologies in the writing process

During the preparation of this work the authors used ChatGPT to assist with grammar revision. After using this tool/service, the authors reviewed and edited the content as needed and take full responsibility for the content of the publication.

## Authors’ contributions

Study conceptualization and design: Jie Wang, Xiangqian Kong, Jiongliang Wang, Jiawang Tao

Methodology: Jiongliang Wang, Jiawang Tao, Cuicui Xia

Collection of omics data: Jiongliang Wang, Miaoxiu Tang

Bioinformatic analysis: Jiongliang Wang

Experiment: Jiawang Tao, Cuicui Xia

Acquisition of samples and data: Jiongliang Wang, Jiawang Tao, Cuicui Xia, Shengxian Yuan

Drafting of the manuscript: Jie Wang, Xiangqian Kong, Jiongliang Wang

Critical revision of the manuscript: Jiawang Tao, Andrei-Florian Stoica, Shengxian Yuan, Yinxiong Li, Lin Guo, Xiangqian Kong, Jie Wang

Conceptualization, writing-review & editing, supervision, funding acquisition: Jie Wang, Xiangqian Kong, Lin Guo, Yinxiong Li

## Supplemental Figure and Figure Legend

**Figure S1.**
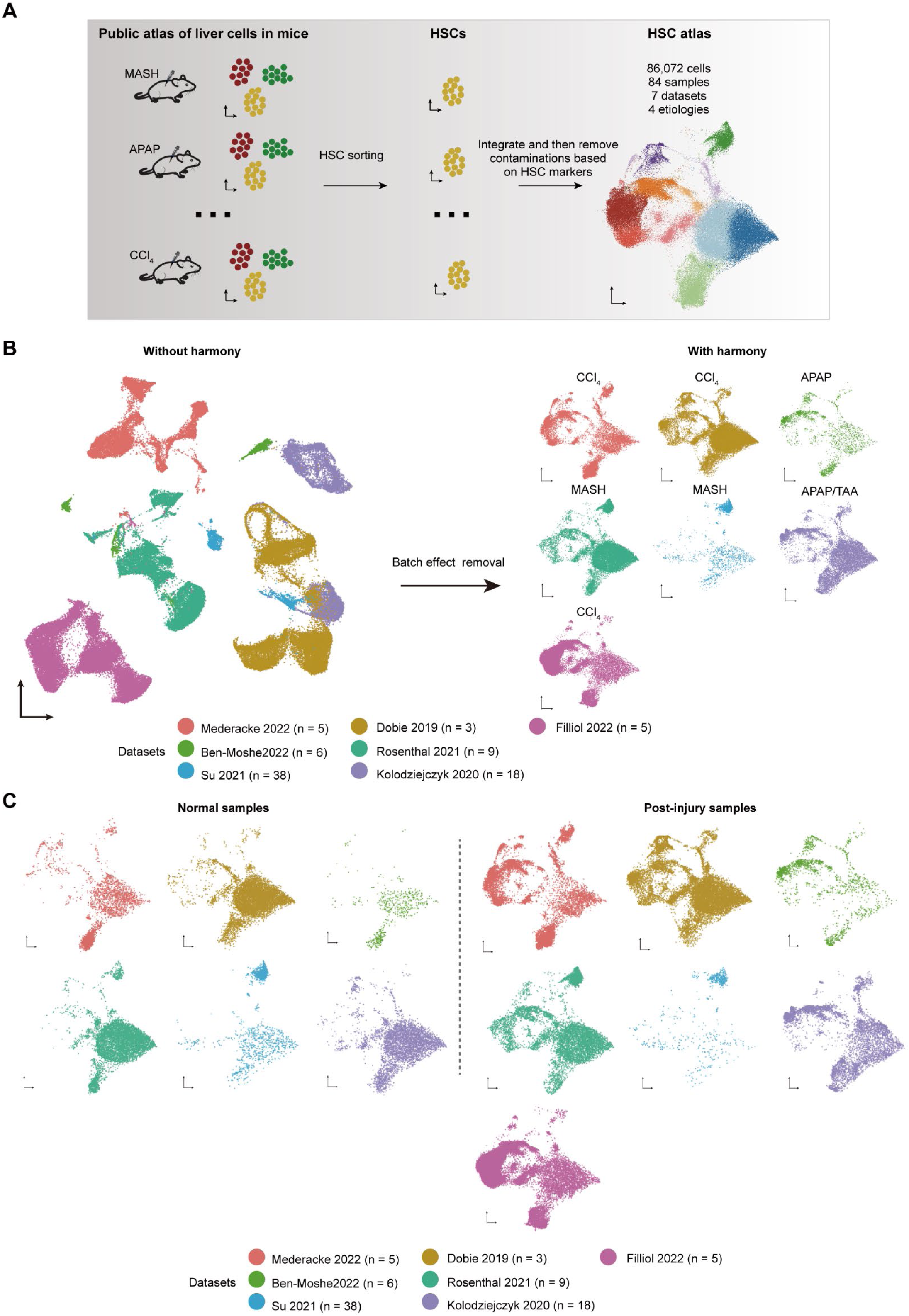
Data integration and batch effect removal for mouse HSC atlas construction. **(A)** Schematic workflow for integrating mouse liver HSC data from Mederacke 2022, Dobie 2019, Filliol 2022, Ben-Moshe2022, Rosenthal 2021, Su 2021, and Kolodziejczyk 2020. Public atlas datasets from multiple liver injury models were processed to extract HSCs, followed by integration and contamination removal based on HSC markers, resulting in a comprehensive HSC atlas. **(B)** UMAP visualization demonstrating batch correction efficacy. Left: Pre-integration data showing substantial batch effects with dataset-specific clustering. Right: Harmony integration showing successful batch correction. Colors indicate individual datasets as labeled. **(C)** Comparison of integrated data structure between normal and post-injury samples. Left: Distribution of cells from normal/healthy samples across datasets. Right: Distribution of cells from post-injury samples, demonstrating consistent integration across experimental conditions. Each dataset is represented by distinct colors with sample sizes indicated in parentheses.

**Figure S2.**
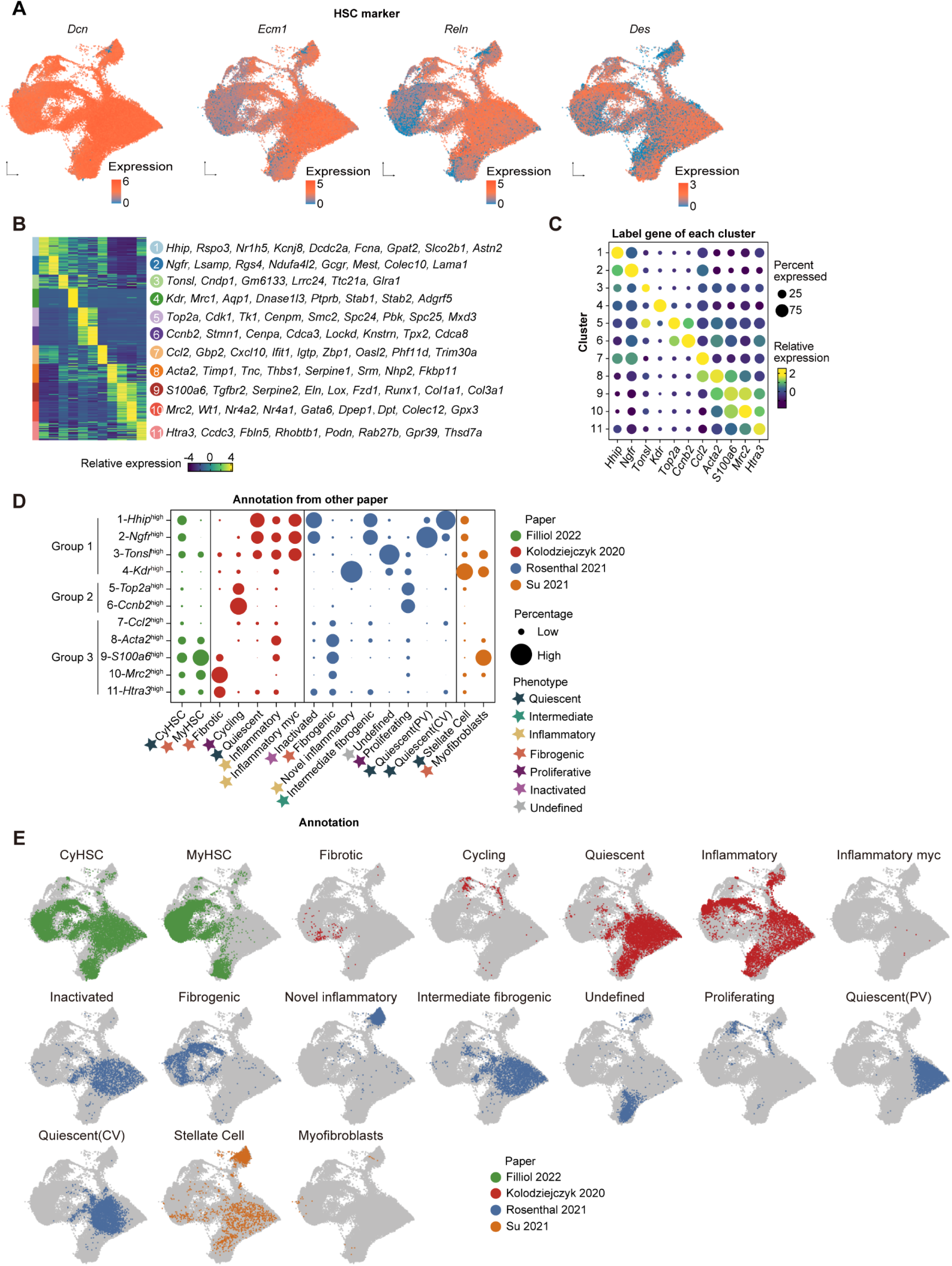
Analysis of HSC cell identity and marker gene expression across subtypes. **(A)** Validation of HSC identity through canonical marker gene expression. UMAP visualization showing expression patterns of HSC markers. **(B)** Heatmap displaying relative expression of cluster-defining marker genes. **(C)** Dot plot showing expression patterns of label genes across 11 HSC clusters. **(D)** Mapping of previously reported HSC phenotypes to current cluster annotations from Filliol 2022, Rosenthal 2021, Su 2021, and Kolodziejczyk 2020. Dot plot comparing cluster identities with published HSC subtypes from five independent studies **(E)** Spatial distribution of HSC phenotypes across the integrated atlas. UMAP projections highlighting cells from different studies colored by previously assigned phenotypes.

**Figure S3.**
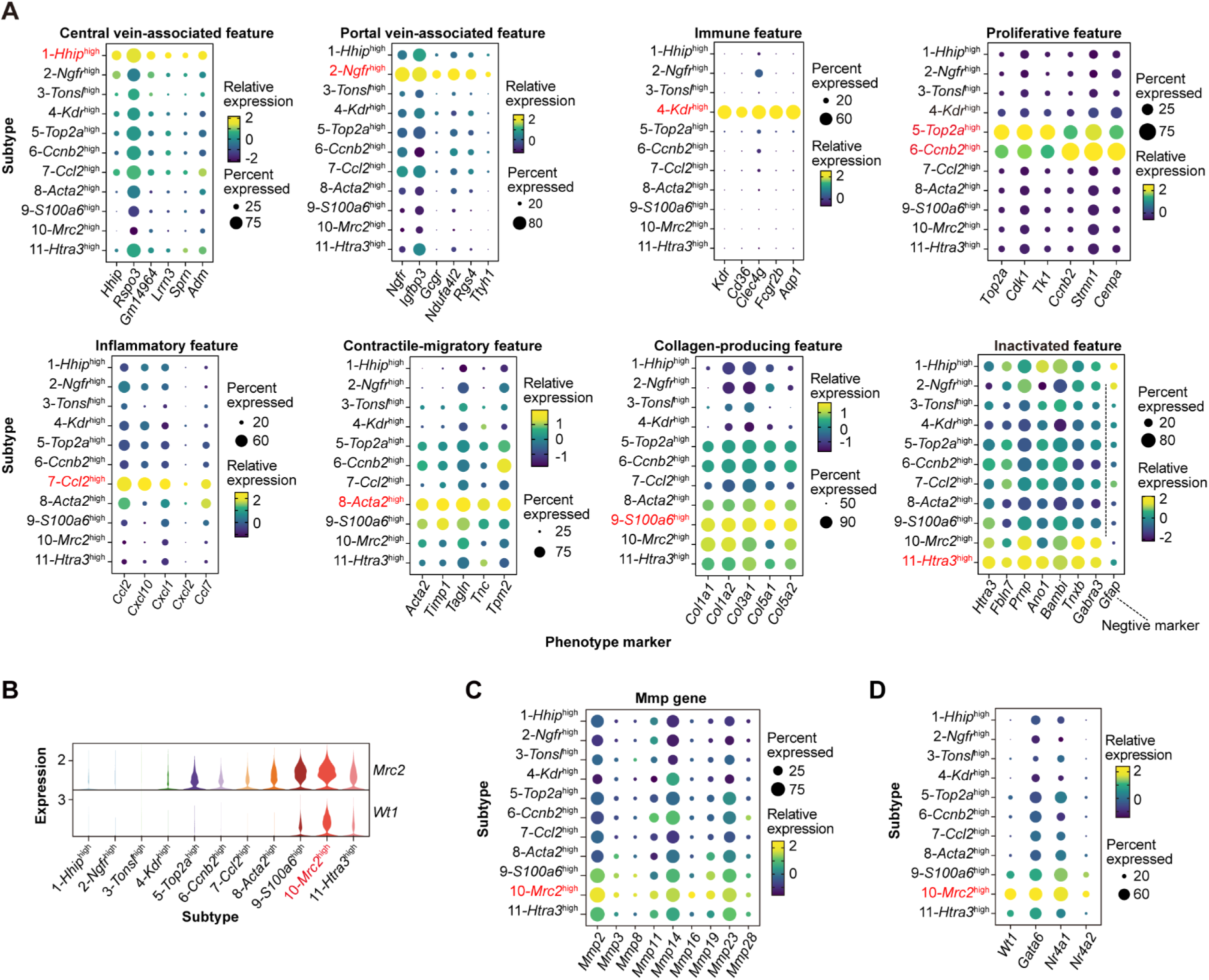
Analysis of HSC feature-related markers across subtypes. **(A)** Expression profiles of HSC phenotype markers. Dot plots showing marker gene expression patterns categorized by phenotypic features: central vein-associated, portal vein-associated, immune-inflammatory-senescence, inflammatory, contractile-migratory, collagen-producing, and inactivated states. *Gfap* serves as a negative marker for inactivated HSCs. **(B)** Violin plots displaying expression of *Mrc2* and *Wt1* across 11 HSC subtypes. **(C)** Dot plot showing expression of Mmp genes across HSC clusters. **(D)** Dot plot showing *Wt1*, *Gata6*, *Nr4a1*, and *Nr4a2* expression across HSC clusters.

**Figure S4.**
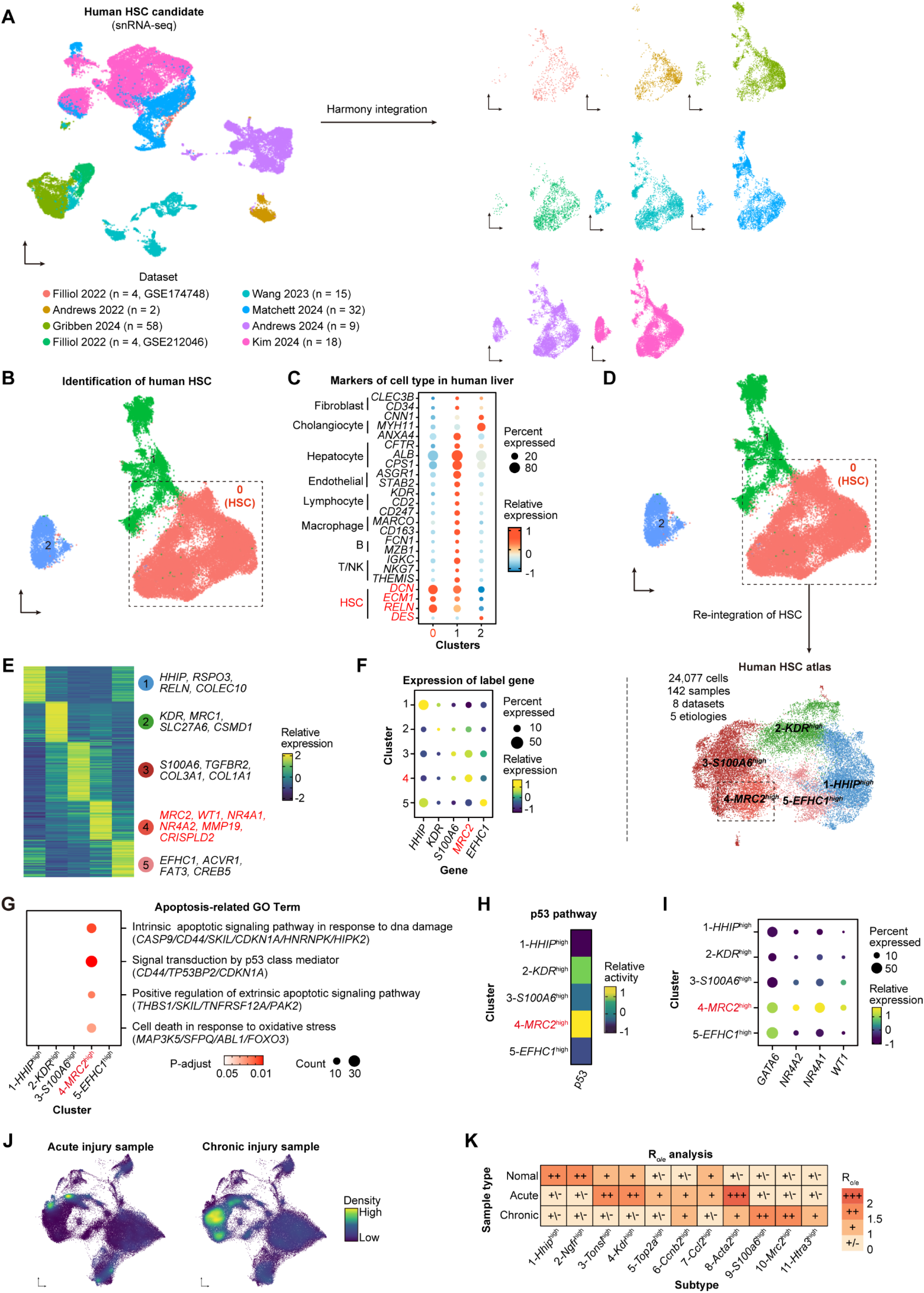
Integration of snRNA-seq datasets from human HSCs identified cross-species conserved *MRC2*^high^ HSCs. **(A)** UMAP visualization of human HSC integration from 8 snRNA-seq datasets (Filliol 2022, Wang 2023, Andrews 2022, Matchett 2024, Gribben 2024, Andrews 2024, and Kim 2024). Left: Pre-integration data showing batch effects with dataset-specific clustering. Right: Harmony integration displaying successful batch correction while preserving biological variation. Colors indicate individual datasets with sample sizes in parentheses. **(B)** UMAP visualization of re-clustered integrated cells. Cluster 0 was identified as HSCs (red dashed box) based on specific marker expression profiles. **(C)** Dot plot validating cell type identities across clusters. HSC-specific markers show enriched expression in cluster 0. **(D)** Schematic workflow for human HSC atlas construction. Re-integration of validated HSCs from cluster 0 yielding refined atlas. **(E)** Heatmap showing relative expression of cluster-defining marker genes across 5 human HSC clusters. Key subtype markers were shown. **(F)** Expression patterns of label genes across human HSC clusters. **(G)** Enrichment analysis of apoptosis-related GO terms across HSC clusters. Scatter plot showing term significance (p-value) and count, with representative genes listed for each pathway. **(H)** PROGENy p53 pathway activity across human HSC clusters. Heatmap showing differential pathway activation scores. **(I)** Dot plot displaying expression of 4 HSC activation suppressors across human HSC clusters. **(J-K)** Stage preferences of HSC subtypes shown by cell density (J) and R_o/e_ analysis (K).

**Figure S5.**
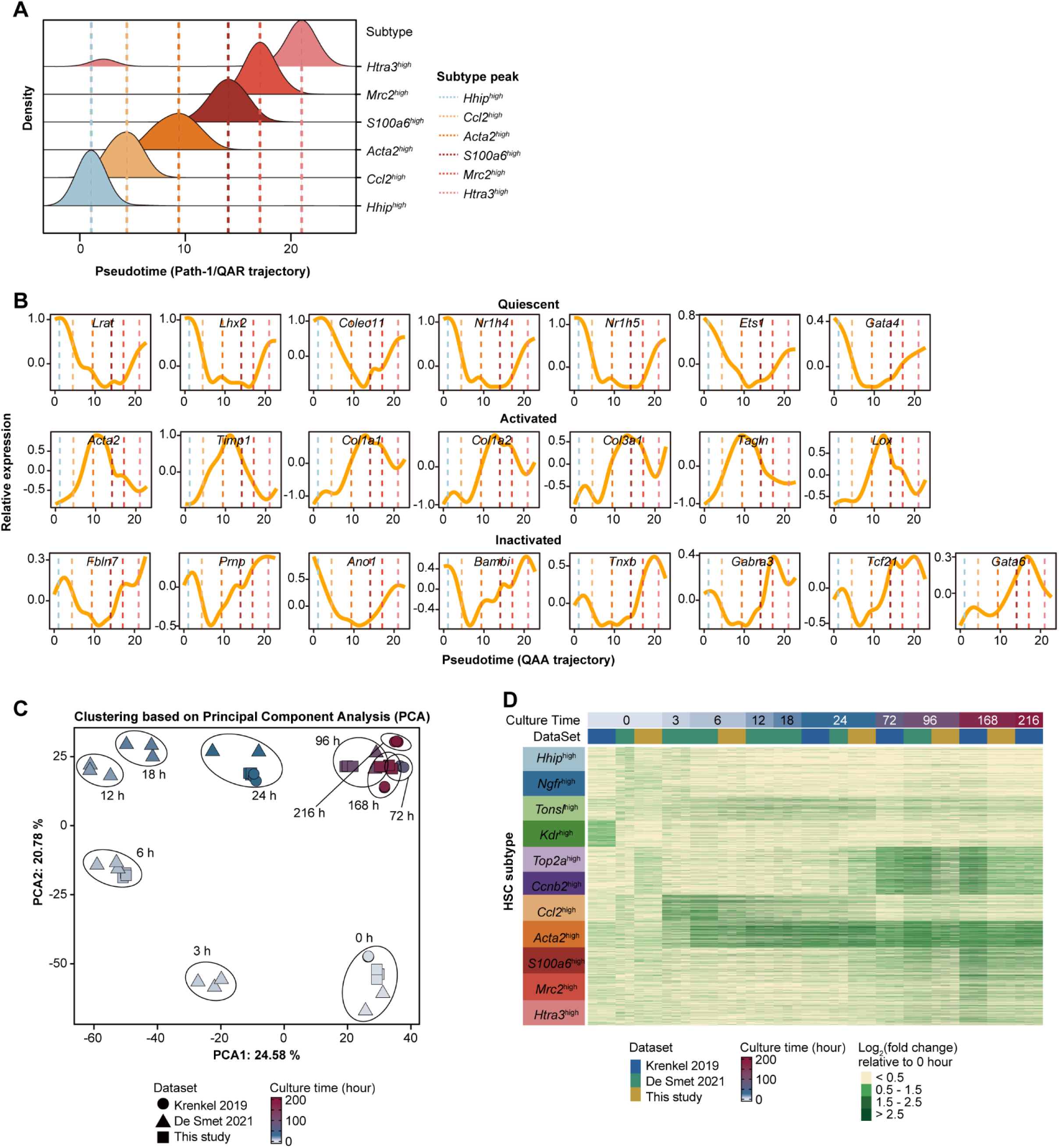
Validation of sequential HSC subtype marker expression. **(A)** Cell density distribution of HSC subtypes along the QAA trajectory. Density plots showing temporal peaks for each subtype, with dashed lines indicating peak density points for trajectory progression analysis. **(B)** Dynamic expression patterns of HSC state markers along the QAA trajectory. Smoothing spline curves showing relative expression of quiescent (top row), activated (middle row), and inactivated (bottom row) HSC markers. Dashed lines correspond to peak density points of HSC subtypes from panel A. **(C)** Principal component analysis of integrated *in-vitro* HSC activation datasets. PCA plot demonstrating minimal batch effects across three independent studies. **(D)** Temporal dynamics of subtype-specific gene expression during *in-vitro* activation. Heatmap displaying log₂ fold changes of top 400 markers (ordered by fold change) for each HSC subtype relative to 0-hour baseline. Top color bar indicates culture time points and data sources. Expression patterns validate the sequential activation of subtype markers corresponding to the QAA trajectory progression.

**Figure S6.**
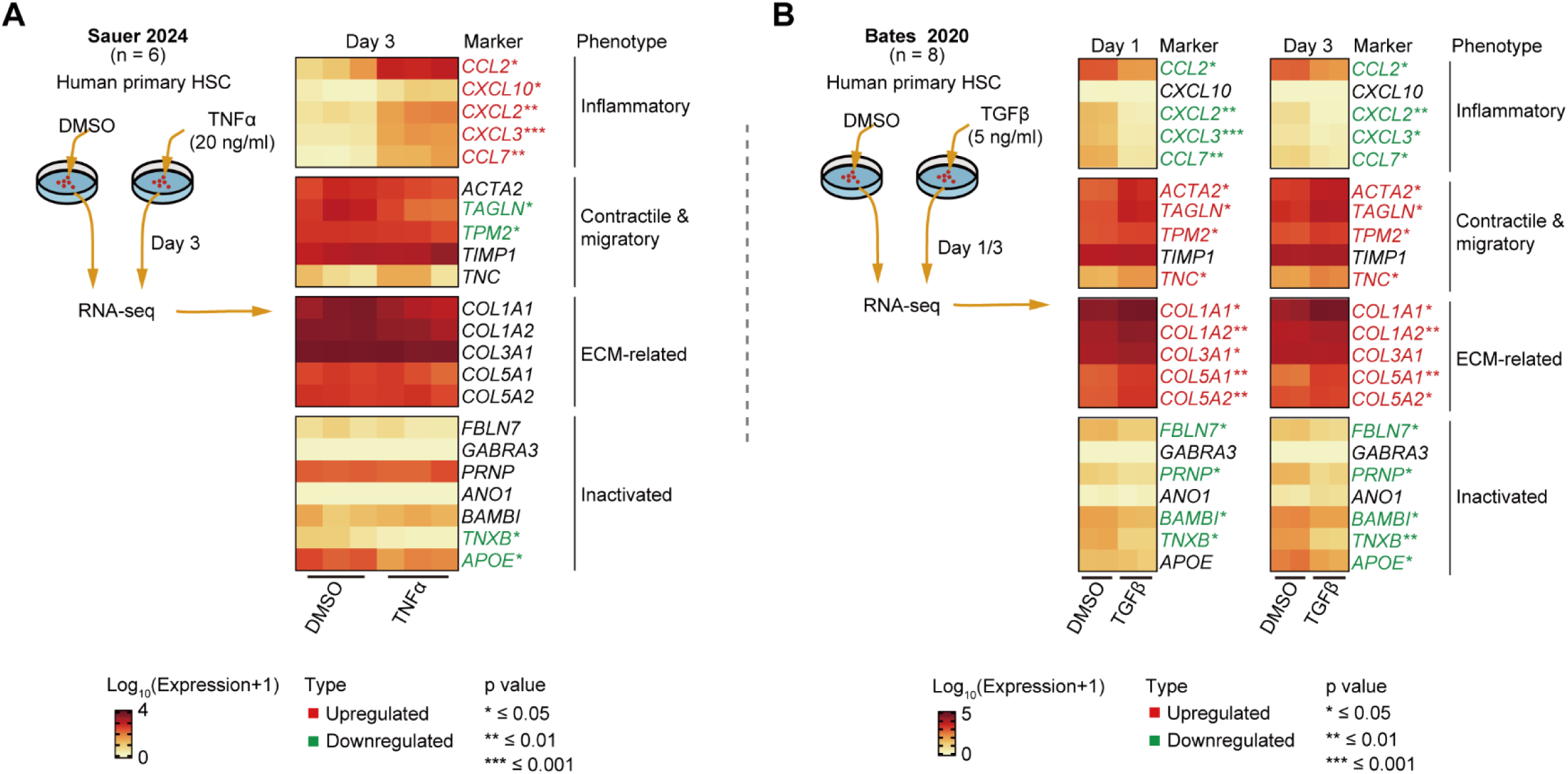
Validation of pathway-driven HSC state transitions in human primary HSCs. **(A)** TNFα-induced activation of human primary HSCs (Sauer 2024). Left: Experimental design showing treatment with TNFα (20 ng/ml) versus DMSO control for 3 days. Right: Heatmap displaying log₁₀ (expression+1) of phenotype-specific markers. Statistical significance indicated for upregulated (red) and downregulated (green) genes. **(B)** TGFβ-mediated activation of human primary HSCs (Bates 2020). Left: Experimental design showing treatment with TGFβ (5 ng/ml) versus DMSO control sampled at days 1 and 3. Right: Heatmaps showing log₁₀ (expression+1) at both time points. Significance levels: ns, p ≥ 0.05; *, p < 0.05; **, p < 0.01; ***, p < 0.001; ****, p < 0.0001 (unpaired two-side Wilcoxon test).

**Figure S7.**
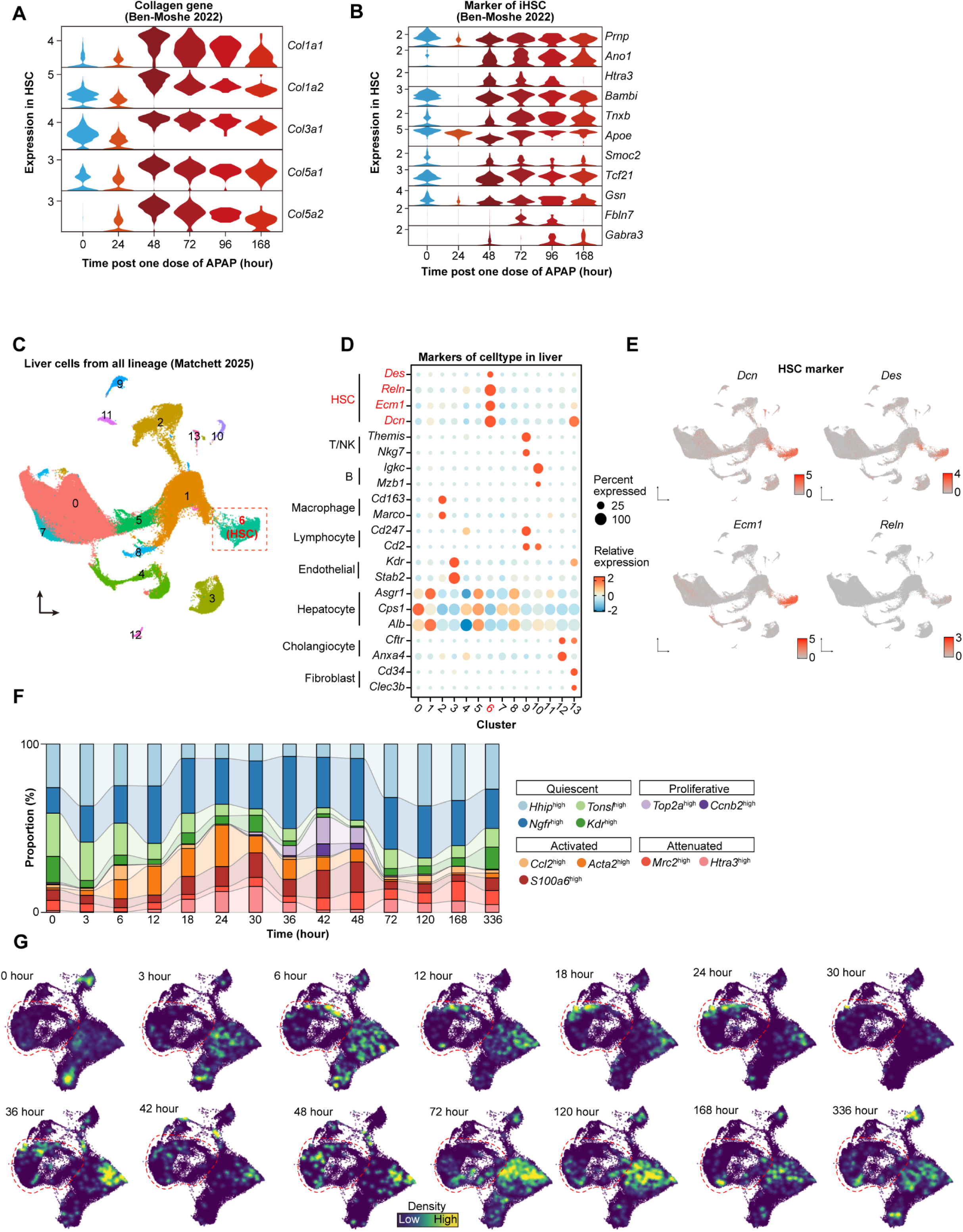
Validation of HSC dynamics during acute liver injury using independent datasets. **(A-B)** Temporal expression patterns following acute APAP injury. **(A)** Violin plots showing collagen gene expression in HSCs at multiple time points (0-168 hours) post-APAP injection. **(B)** Expression dynamics of inactivated HSC markers across the same time course. **(C-G)** Independent validation using snRNA-seq dataset from Matchett 2025. **(C)** UMAP visualization of integrated liver cell populations. Cluster 4 identified as HSCs (boxed region). **(D)** Dot plot showing cell type marker expression across clusters, confirming HSC identity. **(E)** UMAP projections displaying spatial distribution of canonical HSC markers. **(F)** Temporal dynamics of HSC subtype composition during ALI progression. Stacked bar plot showing proportional changes of quiescent, proliferative, activated, and resolutive HSC populations across 14 time points (0-336 hours). **(G)** Cell density visualization along the time course. UMAP density plots demonstrating temporal evolution of HSC populations.

**Figure S8.**
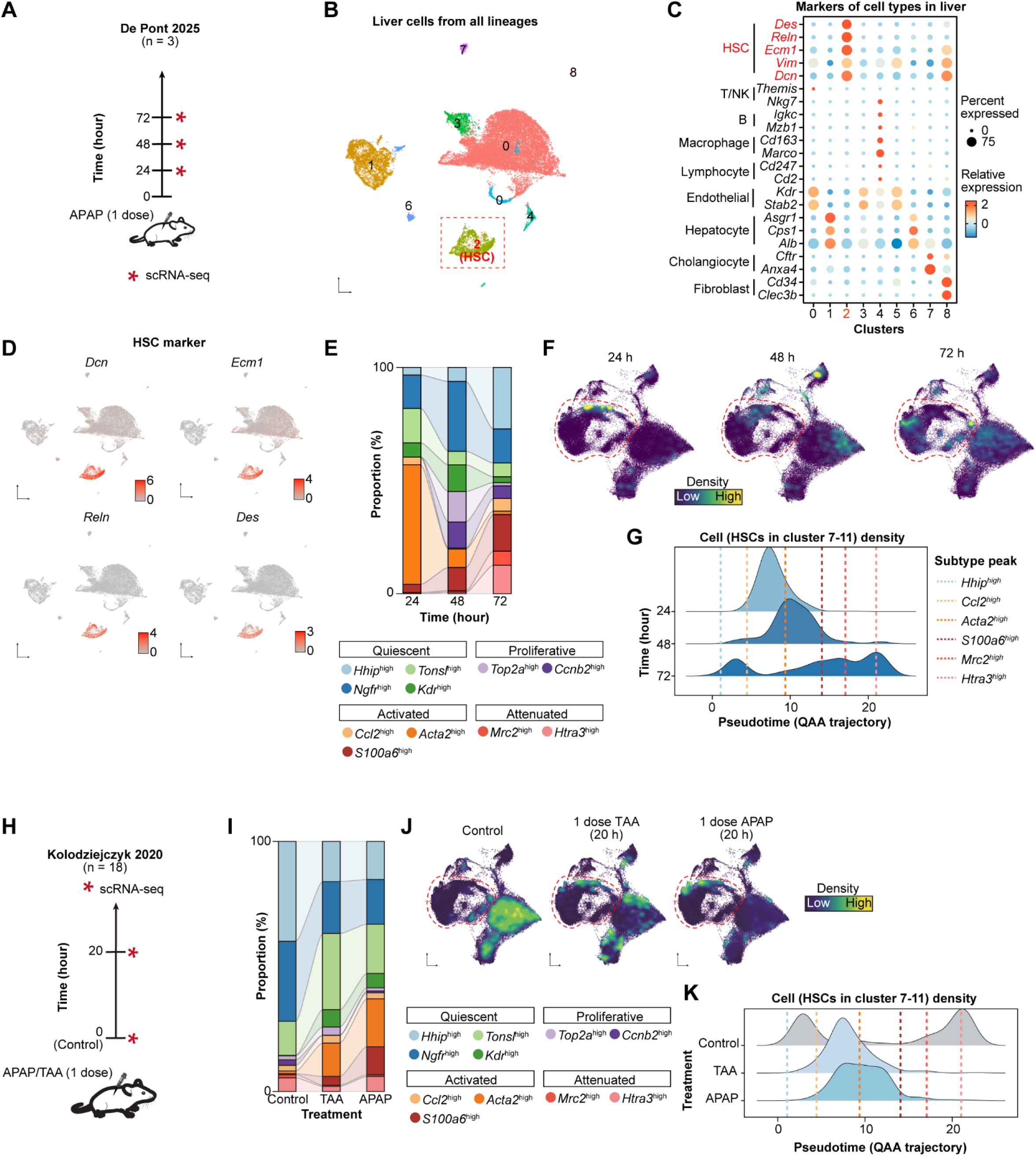
Independent validation of HSC dynamics during acute liver injury using additional datasets. **(A-G)** Validation using scRNA-seq dataset from De Ponti 2025. **(A)** Experimental design showing APAP injection timeline with sampling at 24, 48, and 72 hours. **(B)** UMAP visualization of integrated liver cell populations. Cluster 2 identified as HSCs (dashed box). **(C)** Dot plot confirming cell type identities across clusters through marker expression. **(D)** UMAP validation showing expression pattern of canonical HSC markers. **(E)** Temporal dynamics of HSC subtype composition at 24-, 48-, and 72-hours post-injury. **(F)** UMAP density plots showing HSC population changes over time. **(G)** Ridge plot showing cell density distribution of HSC clusters 7-11 along the QAA pseudotime trajectory. Dashed lines indicate peak density for each subtype. **(H-K)** Validation using scRNA-seq dataset from Kolodziejczyk 2020. **(H)** Experimental design comparing control, TAA, and APAP treatments. **(I)** HSC subtype composition across treatment conditions. **(J)** UMAP density plots showing HSC distribution in control, TAA (20 hours), and APAP (20 hours) treatments. **(K)** Ridge plots comparing cell density distribution of HSC clusters 7-11 along the QAA trajectory between treatments. Dashed lines indicate peak density positions demonstrating treatment-specific activation patterns.

**Figure S9.**
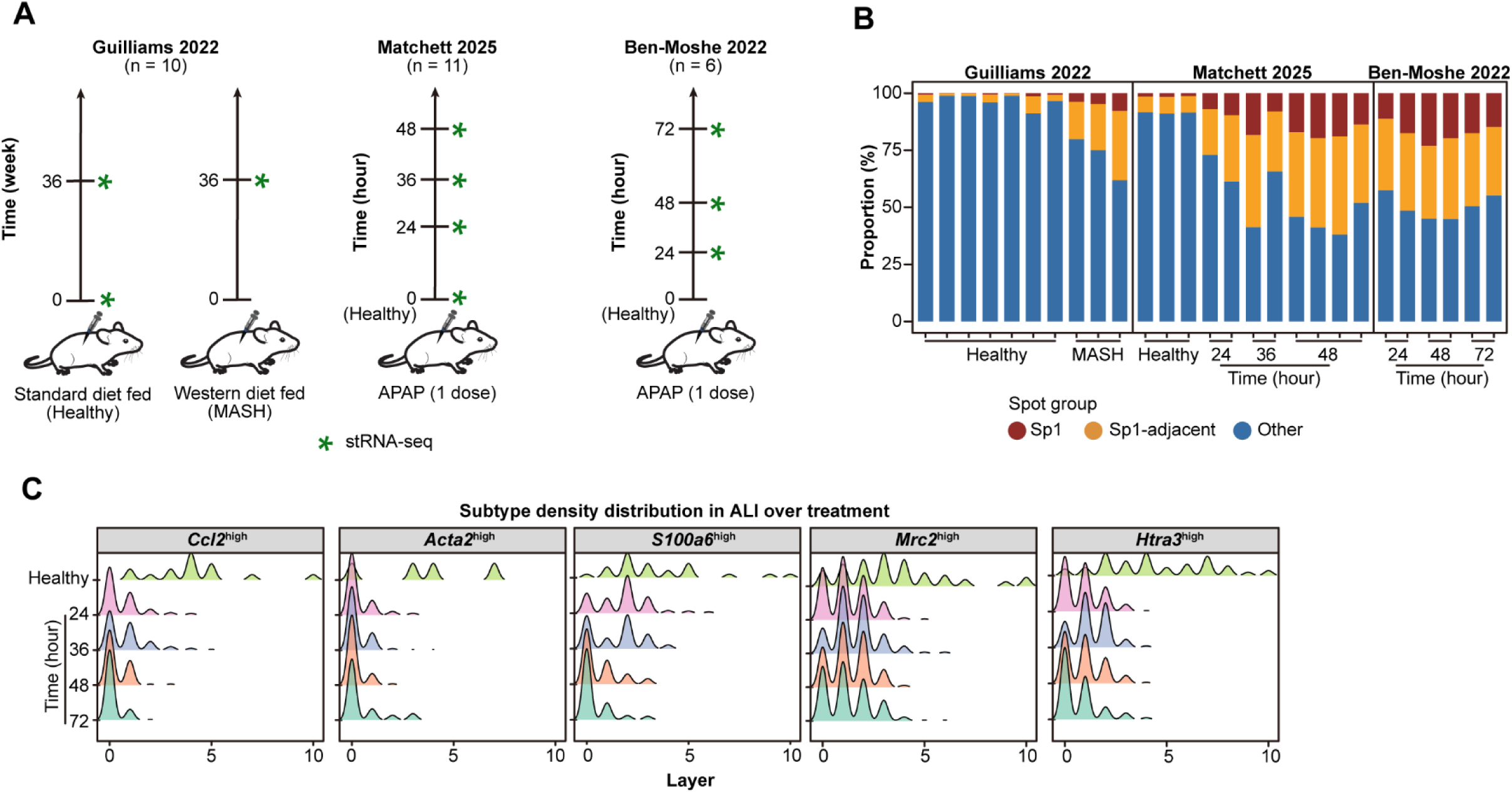
Spatial transcriptomic analysis reveals spatiotemporal organization of HSC subtypes. **(A)** Experimental designs for three spatial transcriptomic datasets used in the integrated analysis. **(B)** Temporal dynamics of spatial group proportions across datasets. Stacked bar plot showing distribution of Sp1, Sp1-adjacent, and other regions in healthy, MASH, and ALI samples at different time points (24-72 hours). Sp1 and Sp1-adjacent regions expand following injury, peaking at 48 hours. **(C)** Concentric layer analysis of HSC subtype distribution during ALI progression. Density profiles of *Ccl2*^high^, *Acta2*^high^, *S100a6*^high^, *Mrc2*^high^, and *Htra3*^high^ positive spots across radial layers from injury epicenter.

**Figure S10.**
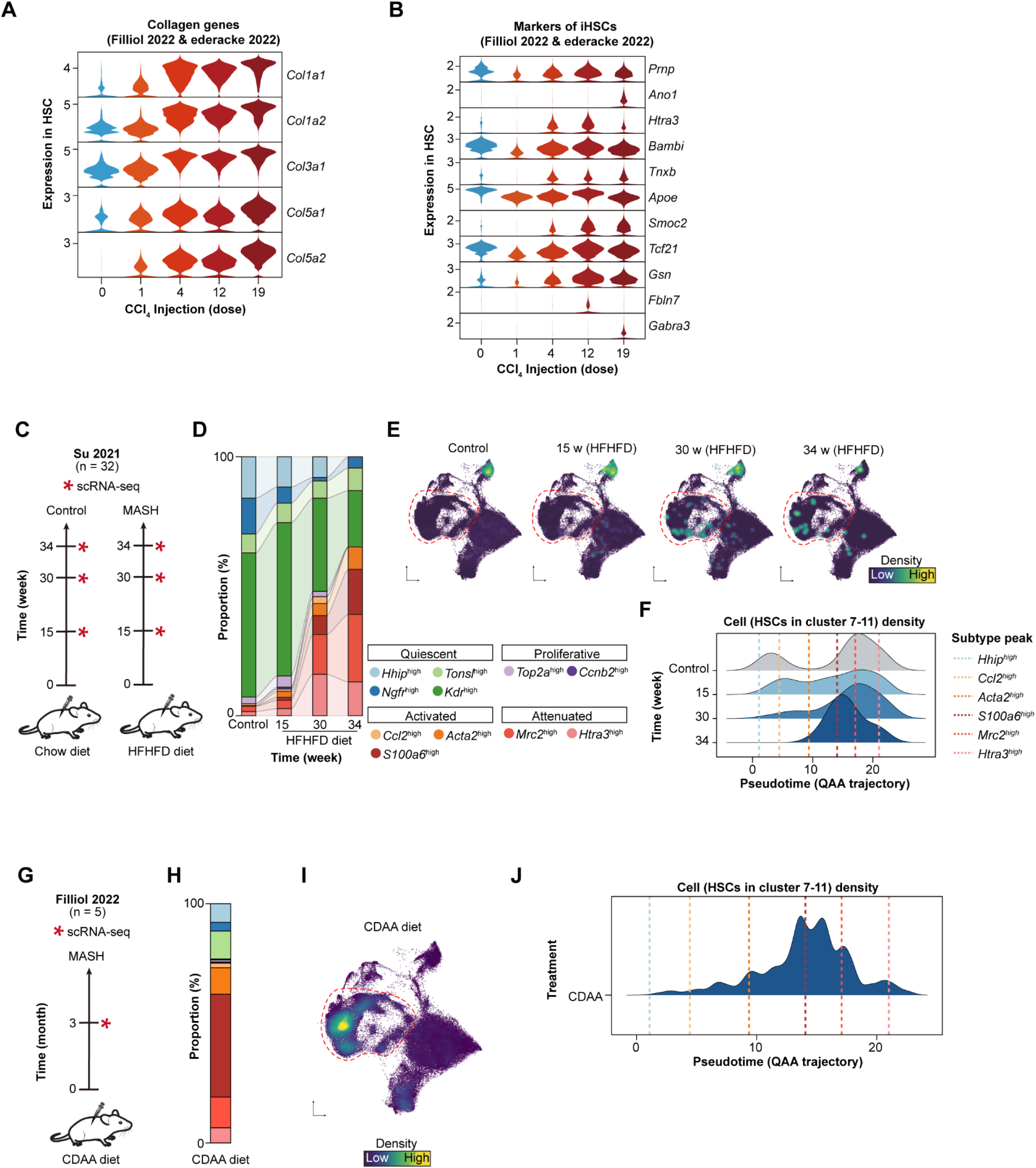
Validation of HSC dynamics during chronic liver injury using integrated datasets. **(A-B)** Temporal expression patterns during CCl₄-induced chronic injury. **(A)** Violin plots showing collagen gene expression across CCl₄ injection numbers (0-19). **(B)** Expression dynamics of inactivated HSC markers during injury progression. **(C-F)** Validation using HFHFD mouse model using scRNA-seq data from Su 2021. **(C)** Experimental design comparing control chow diet versus HFHFD at 15, 30, and 34 weeks. **(D)** HSC subtype composition showing progressive shifts toward activated and resolutive populations. **(E)** UMAP density plots demonstrating temporal HSC distribution changes across HFHFD progression. **(F)** Ridge plot showing cell density of HSC clusters 7-11 along the QAA pseudotime trajectory. Dashed lines indicate peak density for each subtype. **(G-J)** Validation using CDAA diet-induced MASH model using scRNA-seq from Filliol 2022. **(G)** Experimental timeline showing CDAA diet treatment for 3 months. **(H)** HSC subtype composition in CDAA-induced MASH. **(I)** UMAP density visualization showing HSC population distribution. **(J)** Ridge plot displaying cell density along the QAA trajectory following CDAA treatment. Abbreviations: HFHFD (High-Fat-High-Fructose Diet); CDAA (Choline-Deficient, L-Amino Acid-defined diet).

**Figure S11.**
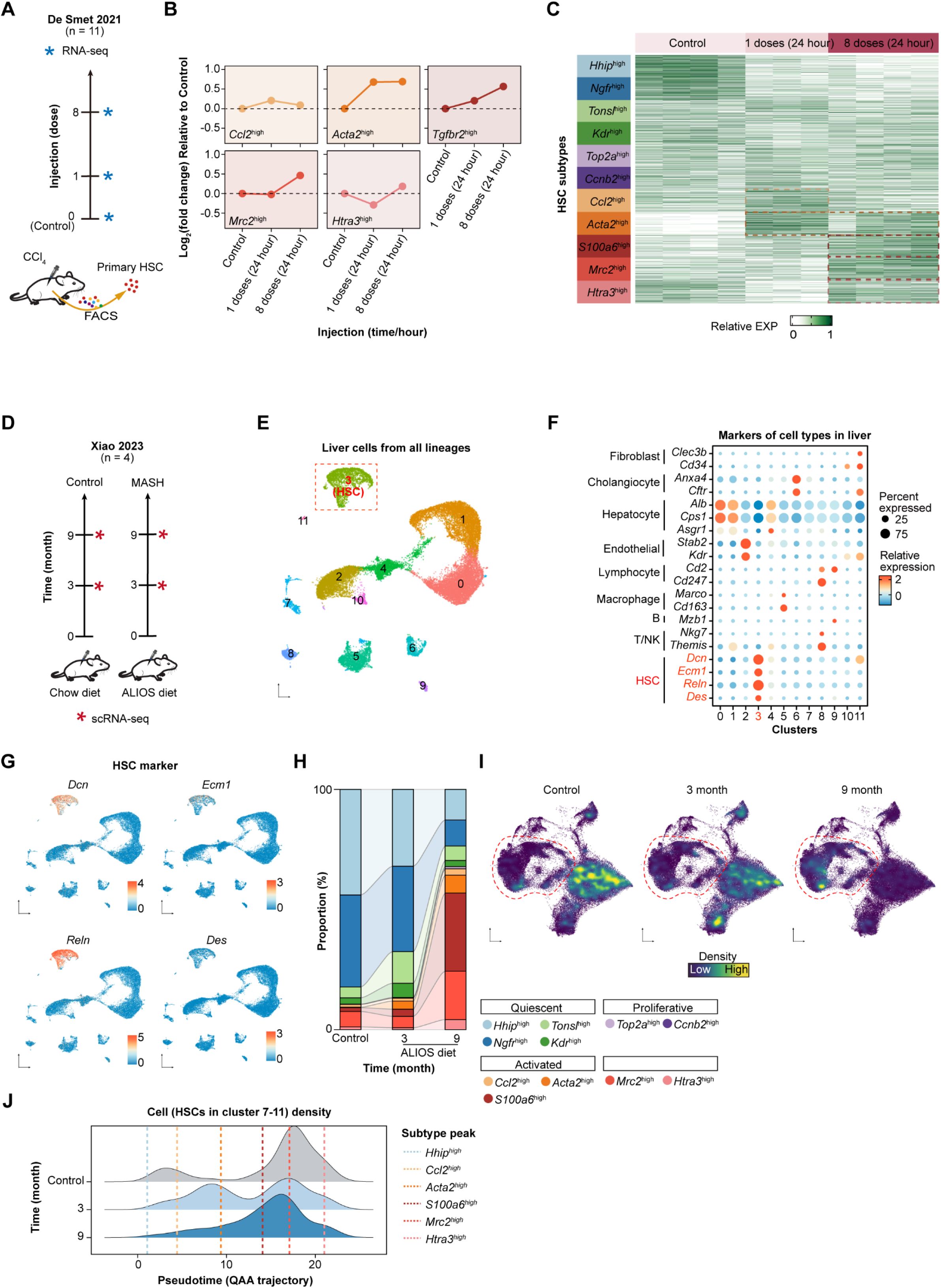
Independent validation of HSC dynamics during chronic liver injury using bulk and single-cell RNA-seq datasets. **(A-C)** Validation using bulk RNA-seq of FACS-isolated HSCs from CCl₄ induced mice model (De Smet 2021). **(A)** Experimental design showing CCl₄ injection timeline with primary HSC isolation for RNA-seq analysis. **(B)** Line plots showing log₂ fold changes relative to control for top 400 upregulated DEGs of HSC subtypes. Early activation markers (*Ccl2*^high^, *Acta2*^high^) peak after single injection, while resolution markers (*Mrc2*^high^, *Htra3*^high^) emerge after 8 injections. Dashed lines indicate baseline. **(C)** Heatmap displaying relative expression dynamics of subtype-specific DEGs across CCl₄ treatment progression. **(D-J)** Validation using scRNA-seq (Xiao 2023) from ALIOS diet model. **(D)** Experimental timeline comparing chow diet versus ALIOS diet at 3 and 9 months. **(E)** UMAP visualization of integrated liver cell populations. HSCs identified in cluster 2 (dashed box). **(F)** Dot plot confirming cell type identities through marker expression across 11 clusters. **(G)** UMAP visualization showing spatial distribution of canonical HSC markers. **(H)** HSC subtype composition at control, 3-month, and 9-month ALIOS diet time points. **(I)** UMAP density plots showing progressive HSC activation with diet duration. **(J)** Ridge plot showing cell density of HSC clusters 7-11 along the QAA trajectory. Dashed lines indicate peak density positions, revealing progressive shifts toward activated and resolutive populations with ALIOS progression. Abbreviation: ALIOS (American Lifestyle-Induced Obesity Syndrome).

**Figure S12.**
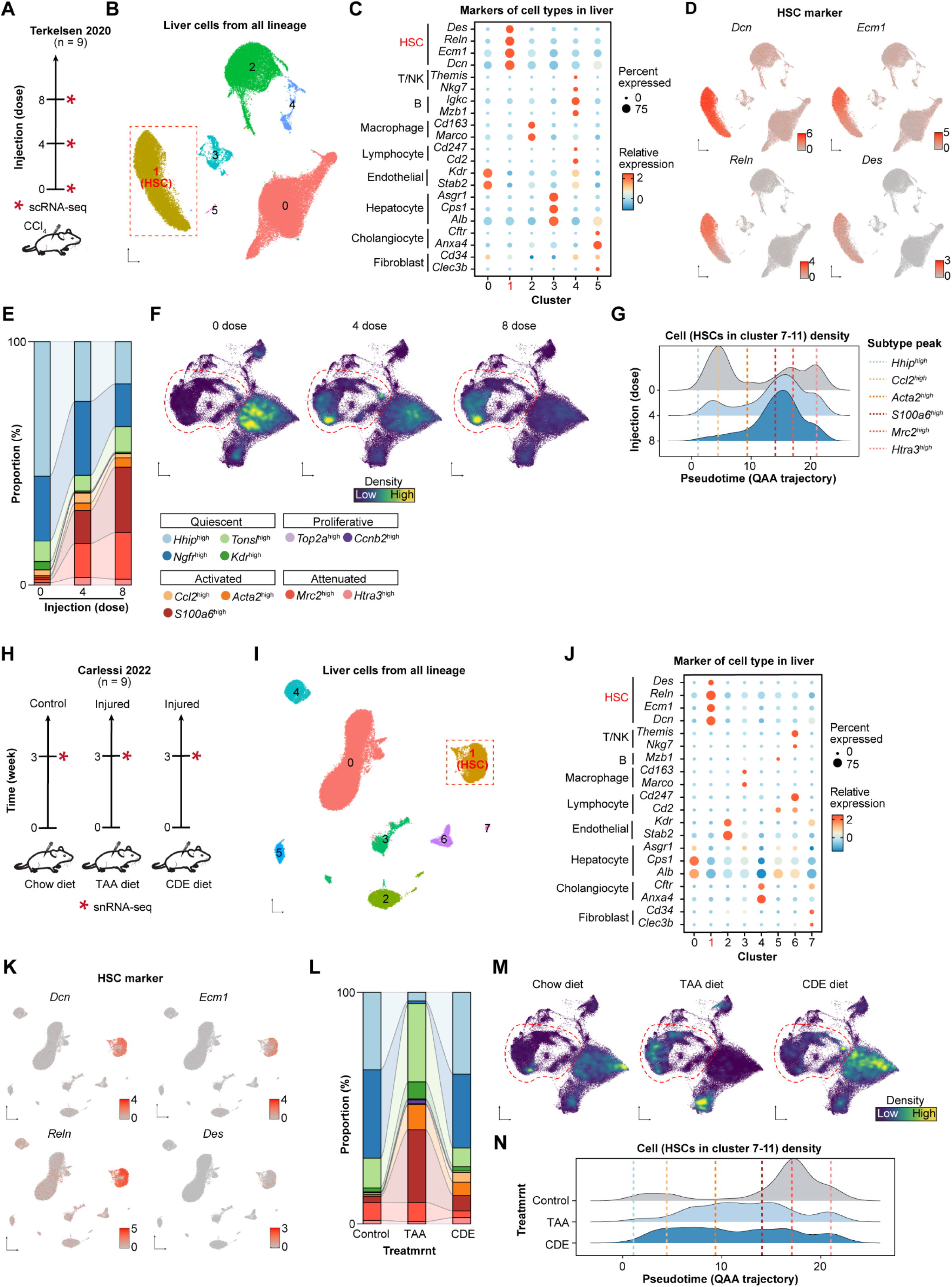
Independent validation of HSC dynamics during chronic liver injury using additional scRNA-seq datasets. **(A-G)** Validation using CCl₄-induced injury model using scRNA-seq data from Terkelsen 2020. **(A)** Experimental design showing CCl₄ injections at 0, 4, and 8 doses. **(B)** UMAP visualization of integrated liver cell populations. HSCs identified in cluster 3 (dashed box). **(C)** Dot plot confirming cell type identities through marker expression across clusters. **(D)** UMAP visualization showing spatial distribution of canonical HSC markers. **(E)** HSC subtype composition across CCl₄ injection numbers. **(F)** UMAP density plots showing progressive HSC activation at 0, 4, and 8 injections. **(G)** Ridge plot showing cell density of HSC clusters 7-11 along the QAA trajectory. Dashed lines indicate peak density positions. **(H-N)** Validation using dietary injury models using snRNA-seq data from Carlessi 2022. **(H)** Experimental timeline comparing chow diet, TAA diet, and CDE diet for 3 weeks. **(I)** UMAP visualization of integrated liver cell populations. HSCs identified in cluster 1 (dashed box). **(J)** Dot plot showing cell type marker expression across clusters. **(K)** UMAP visualization displaying HSC marker distribution. **(L)** HSC subtype composition across different dietary treatments. **(M)** UMAP density plots showing HSC population changes in control, TAA, and CDE diet groups. **(N)** Ridge plot comparing cell density along the QAA trajectory between treatments. Dashed lines indicate peak positions demonstrating diet-specific activation patterns. Abbreviations: TAA (Thioacetamide); CDE (Choline-Deficient, Ethionine-supplemented diet).

**Figure S13.**
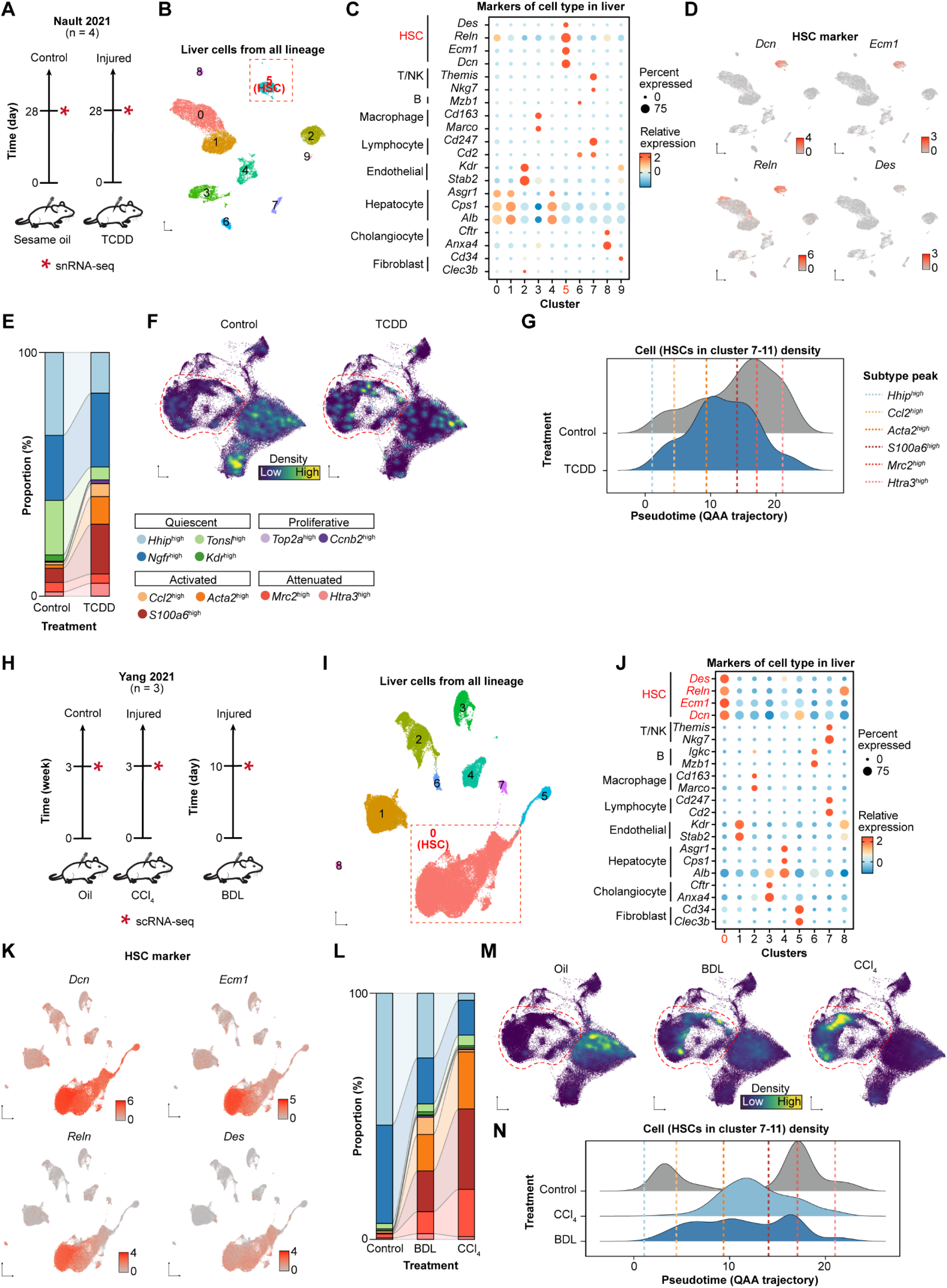
Cross-validation of HSC dynamics during liver injury using independent scRNA-seq datasets. **(A-G)** Validation using TCDD-induced injury model using snRNA-seq data from Nault 2021. **(A)** Experimental design showing sesame oil control versus TCDD treatment at 28 days. **(B)** UMAP visualization of integrated liver cell populations. HSCs identified in cluster 0 (dashed box). **(C)** Dot plot confirming cell type identities through marker expression across 9 clusters. **(D)** UMAP visualization showing spatial distribution of canonical HSC markers. **(E)** HSC subtype composition in control versus TCDD-treated samples. **(F)** UMAP density plots showing HSC population changes following TCDD exposure. **(G)** Ridge plot showing cell density of HSC clusters 7-11 along the QAA trajectory. Dashed lines indicate peak density positions. **(H-N)** Validation using bile duct ligation and CCl₄ injury models using scRNA-seq data from Yang 2021. **(H)** Experimental timeline comparing oil control, CCl₄ injection (3 weeks), and bile duct ligation (10 days). **(I)** UMAP visualization of integrated liver cell populations. HSCs identified in cluster 0 (dashed box). **(J)** Dot plot showing cell type marker expression across clusters. **(K)** UMAP projections displaying HSC marker distribution. **(L)** HSC subtype composition across treatment groups. **(M)** UMAP density plots showing HSC population dynamics in oil control, BDL, and CCl₄ treatments. **(N)** Ridge plot comparing cell density along the QAA trajectory between treatments. Dashed lines indicate peak positions demonstrating treatment-specific activation patterns with CCl₄ and BDL showing distinct activation profiles. Abbreviations: TCDD (2,3,7,8-Tetrachlorodibenzo-p-dioxin); BDL (Bile Duct Ligation).

**Figure S14.**
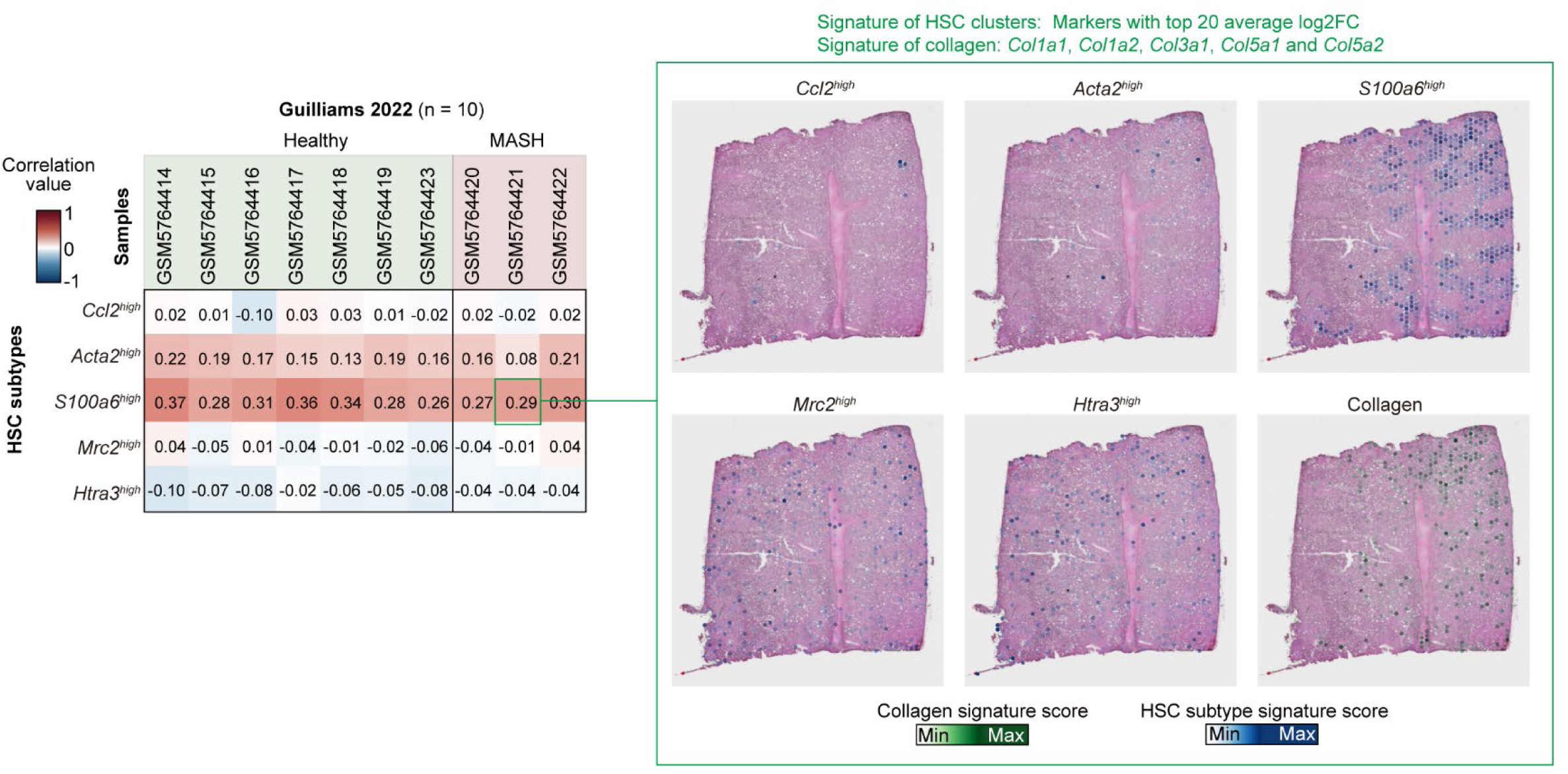
Spatial analysis reveals disrupted HSC subtype distribution and collagen co-localization in chronic liver injury. Spatial co-localization analysis between HSC subtypes and collagen deposition using spRNA-seq data from Guilliams 2022. Left: Correlation matrix showing Pearson coefficients between subtype signature scores and collagen gene expression across samples. *S100a6*^high^ shows strong positive correlation with collagen. Right: Representative HE images and corresponding spatial maps displaying HSC subtype signatures and collagen signature distribution. *S100a6*^high^ co-localizes with collagen-expression zones while *Mrc2*^high^ shows spatial dissociation from collagen-expression zones.

**Figure S15.**
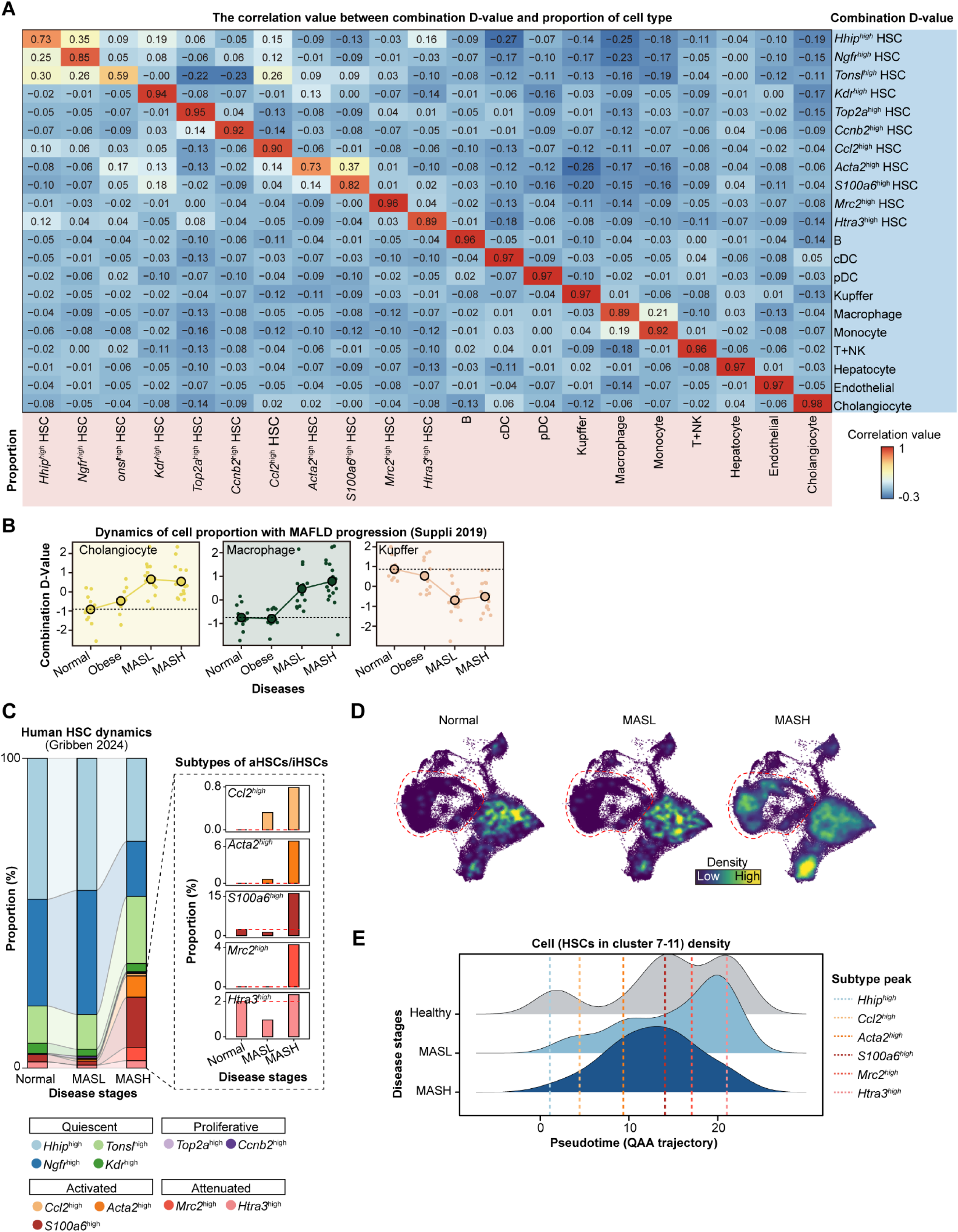
Pseudo-bulk strategy and subtype mapping reveal conserved HSC subtype dynamics across human MAFLD progression. **(A)** Performance evaluation of pseudo-bulk strategy. Heatmap showing correlation coefficients between combination D-values and actual cell type proportions from zenodo.6035873 dataset, demonstrating high specificity for HSC subtype identification. **(B)** Validation of the approach through known cellular dynamics in MAFLD. Line plots displaying temporal changes of cholangiocytes, macrophages, and Kupffer cells across disease stages (healthy, obesity, MASL, MASH) using combination D-values from bulk RNA-seq data. Cholangiocyte and macrophages expansion, along with Kupffer cell reduction confirm established disease-associated cellular shifts. **(C-E)** Validation using MASH patients using snRNA-seq data from Gribben 2024. **(C)** Human HSC subtype composition mapped to mouse HSC atlas. Stacked bar plot showing progressive shifts in HSC subtype proportions across MAFLD stages. *Ccl2*^high^, and *Acta2*^high^ HSCs expand during MASL, *S100a6*^high^, *Mrc2*^high^, and *Htra3*^high^ emerges in MASH. **(D)** Density contour plots illustrating HSC population redistribution during disease progression. UMAP visualizations showing kernel density estimates at healthy, MASL, and MASH stages, revealing progressive shift from quiescent to activated regions along the QAA trajectory. **(E)** Quantitative assessment of HSC activation states along QAA trajectory pseudotime. Ridge plots displaying cell density distributions for HSC clusters along the QAA trajectory axis. Vertical dashed lines indicate density peaks, showing incomplete resolution with persistent activation in MASH.

**Figure S16.**
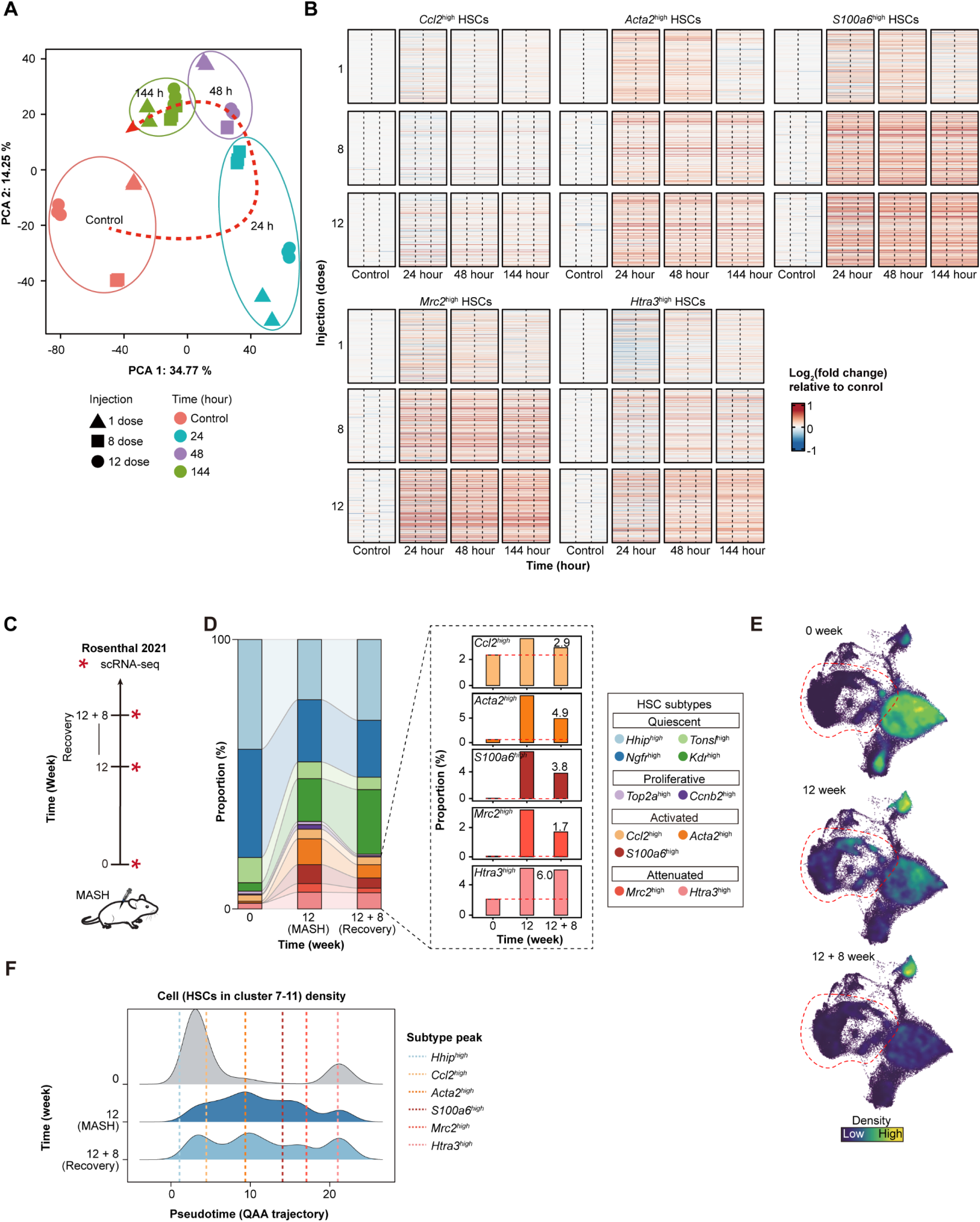
HSC dynamics following injury removal reveal impaired resolution after chronic injury. **(A)** Principal component analysis of bulk RNA-seq data from isolated primary HSCs following CCl₄ withdrawal. Samples grouped by injection number (1, 8, or 12) and recovery time point (0-144 hours). Arrows indicate transcriptomic trajectory during recovery period, showing partial return toward healthy state. **(B)** Temporal expression dynamics of subtype-specific markers following injury cessation. Heatmaps displaying log₂ fold changes relative to control for top 400 markers (ranked by fold change) of five HSC subtypes (*Ccl2*^high^, *Acta2*^high^, *S100a6*^high^, *Mrc2*^high^, and *Htra3*^high^). Expression patterns shown at control, 24, 48, and 144 hours post-final injection for 1, 8, and 12 injection regimens. **(C-F)** Independent validation using MASH regression model using scRNA-seq data from Rosenthal 2021. **(C)** Experimental timeline showing MASH induction and recovery strategy. **(D)** HSC subtype composition at baseline (0 weeks), MASH (12 weeks), and recovery (12+8 weeks). Bar plot showing persistent activated HSCs (11.6%) after 8-week recovery. Inset: Detailed proportions of individual subtypes. **(E)** UMAP density plots demonstrating HSC population distribution at 0, 12, and 12+8 weeks. **(F)** Ridge plot showing cell density of HSC clusters 7-11 along the QAA trajectory. Dashed lines indicate peak density positions, revealing incomplete resolution.

**Figure S17.**
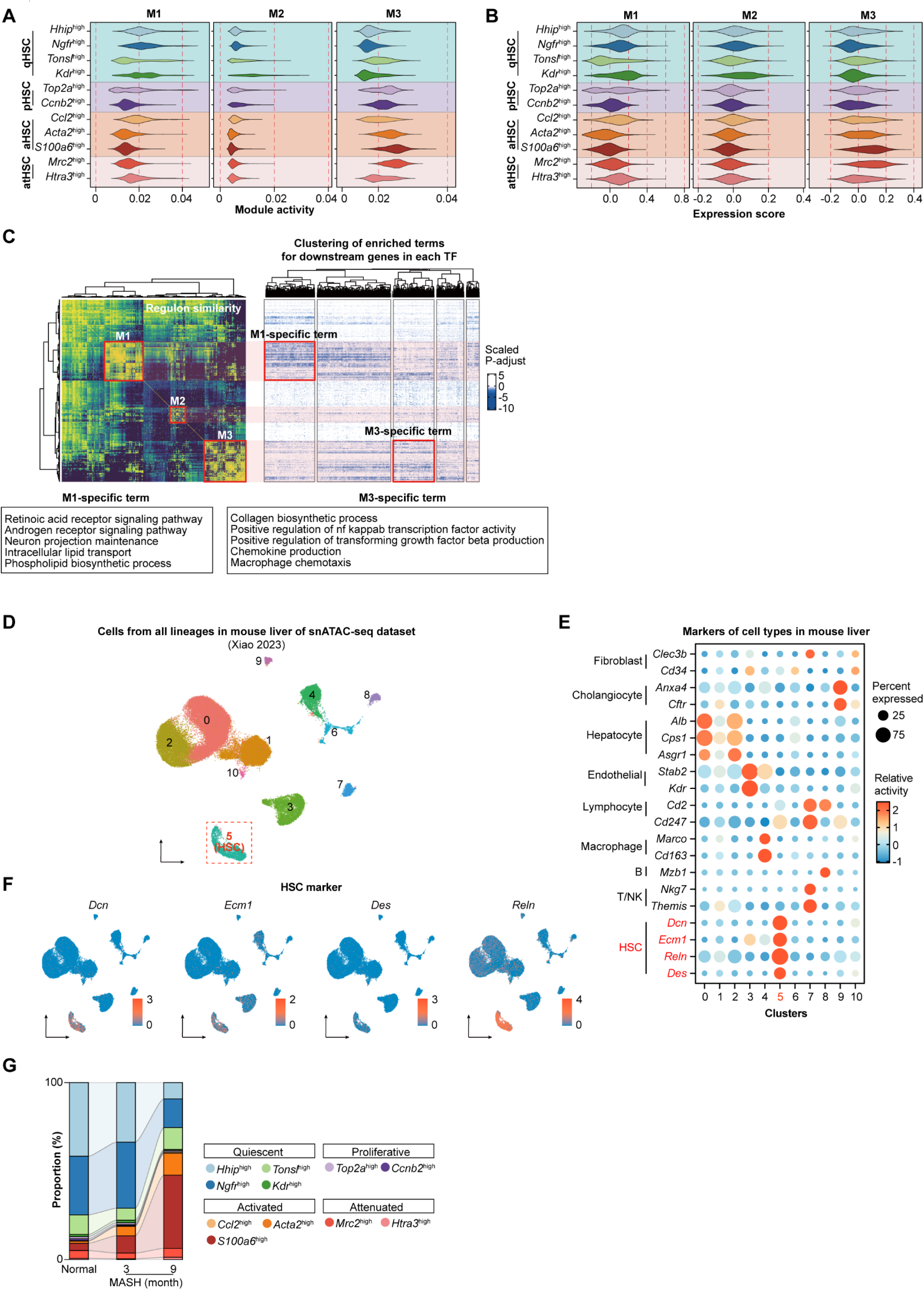
Regulatory module analysis reveals subtype-specific transcriptional programs validated by snATAC-seq. **(A)** Module activity profiles across HSC subtypes. Violin plots showing regulon activity scores for three transcriptional modules (M1-M3) across 11 HSC clusters. M1 shows highest activity in quiescent subtypes, M3 in activated subtypes, and M2 in *Kdr*^high^ populations. **(B)** Expression levels of module-associated transcription factors. Violin plots displaying average TF expression within each regulatory module across HSC subtypes, revealing module-specific expression patterns corresponding to activation states. **(C)** Functional characterization of regulatory modules through target gene analysis. Left: Heatmap showing similarity between regulons based on CSI matrix (as in Figure 5A). Right: Hierarchical clustering of enriched GO terms for downstream genes of each regulon. Module-specific functional categories highlighted.**(D-G)** Independent validation using snATAC-seq data from Xiao 2023. **(D)** UMAP visualization of integrated liver cell populations from chromatin accessibility profiles. Cluster 5 identified as HSCs (dashed box). **(E)** Dot plot confirming cell type identities through gene activity scores of canonical markers across clusters. **(F)** UMAP visualization showing gene activity patterns of HSC markers. **(G)** HSC subtype composition at baseline, 3-month, and 9-month MASH progression, revealing shift from quiescent to activated populations.

**Figure S18.**
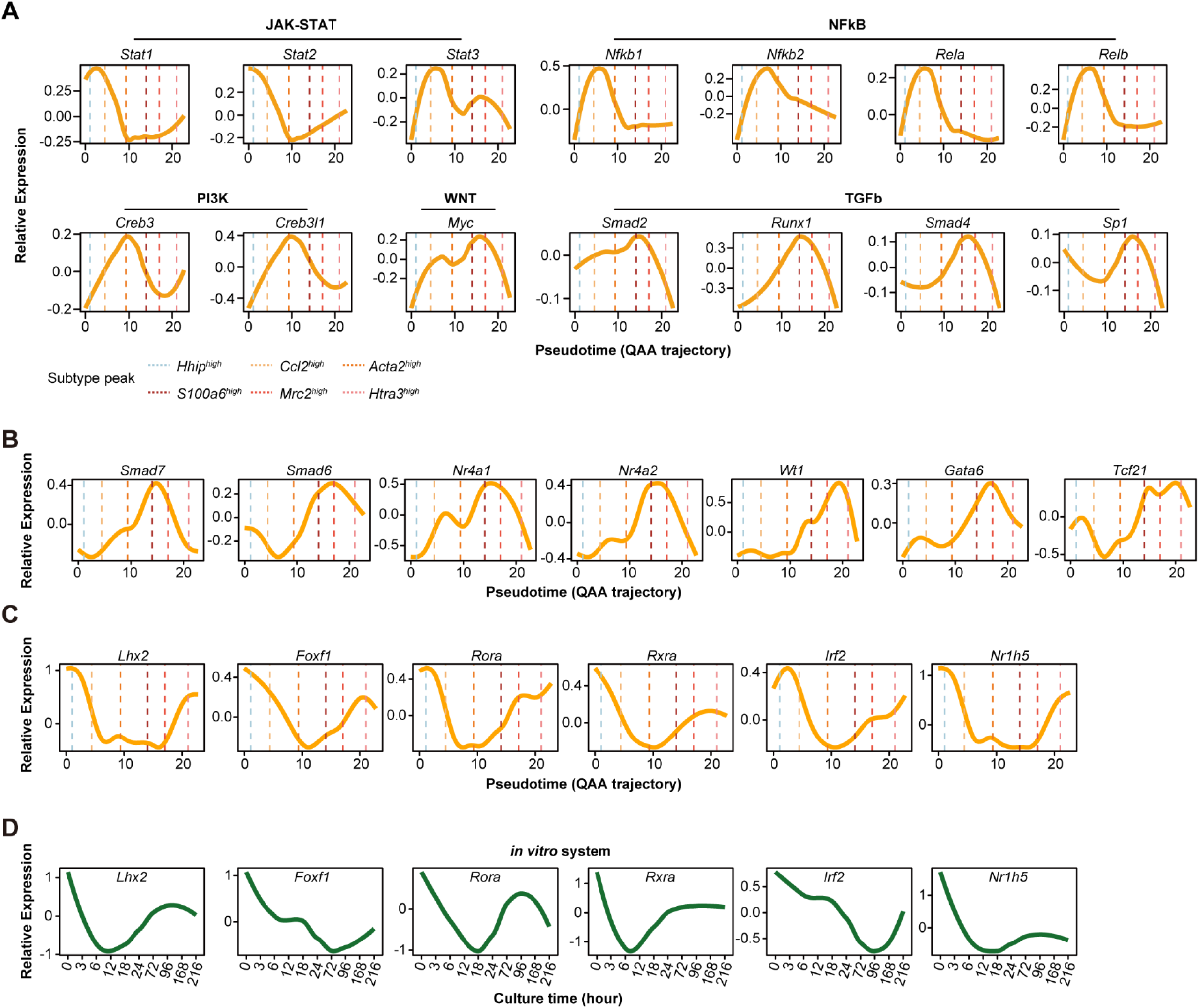
Transcriptional dynamics reveal sequential pathway activation and apoptotic program along the QAA trajectory. **(A-C)** Smoothing spline curves showing relative expression levels of key TFs representing distinct signaling pathways (A), activation suppressors (B) and quiescence-associated (C) along the QAA pseudotime. Vertical dashed lines indicate peak cell density positions for clusters along pseudotime. **(D)** *In vitro* validation of partial quiescence TF dynamics. Smoothing spline curves showing relative expression of quiescence-related TFs across culture time points (0-216 hours) in primary HSCs.

**Figure S19.**
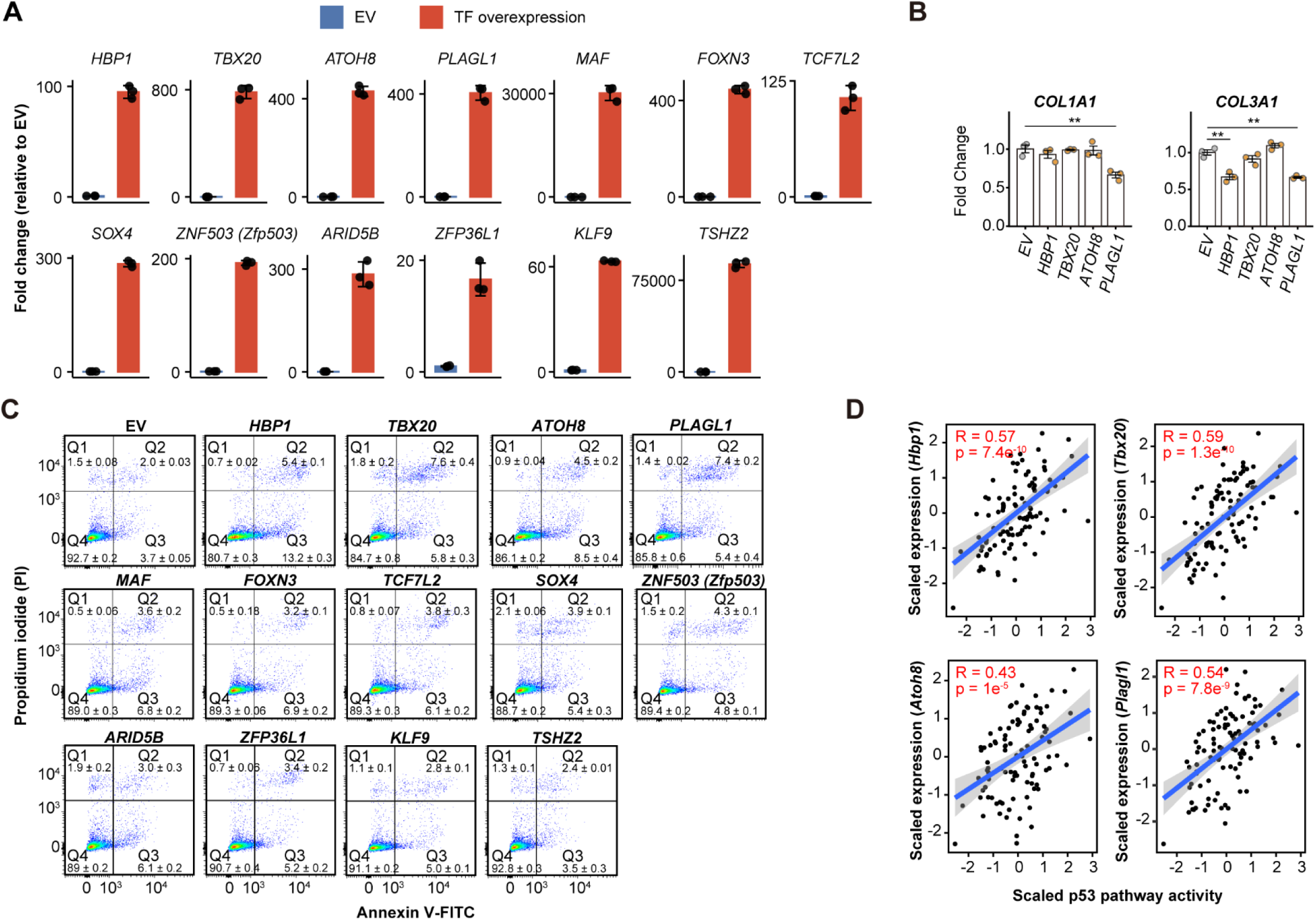
Functional validation identifies pro-apoptotic transcription factors enriched in *Mrc2*^high^ HSCs. **(A)** Validation of TFs overexpression efficiency in LX-2 cells. Bar plots quantifying relative expression levels of candidate TFs following lentiviral transduction compared to empty vector (EV) control. Expression normalized to EV. Data presented as means ± SD (n = 3). **(B)** Flow cytometric analysis showing TF-induced apoptosis in human HSCs. Representative scatter plots showing Annexin V/PI signal in LX-2 cells 8 days post-transduction with indicated TFs versus EV control. Data shown as means ± SD (n = 3), representative of three independent experiments. **(C)** Effect of TF overexpression on fibrogenic gene expression. Fold changes in *COL1A1* and *COL3A1* expression upon overexpression of indicated TFs in LX-2 cells. Data represent mean ± SD (n = 3). **(D)** Correlation analysis linking candidate TFs to p53-mediated apoptosis pathway. Dot plot displaying Pearson correlation coefficients between PROGENy p53 pathway activity scores and expression of *Hbp1*, *Tbx20*, *Atoh8*, and *Plagl1* across HSC clusters along the QAA trajectory using pseudobulk expression profiles. Statistical significance is denoted as: ns, p > 0.05; *, p < 0.05; **, p < 0.01; ***, p < 0.001; ****, p < 0.0001 (unpaired two side Wilcoxon test).

**Figure S20.**
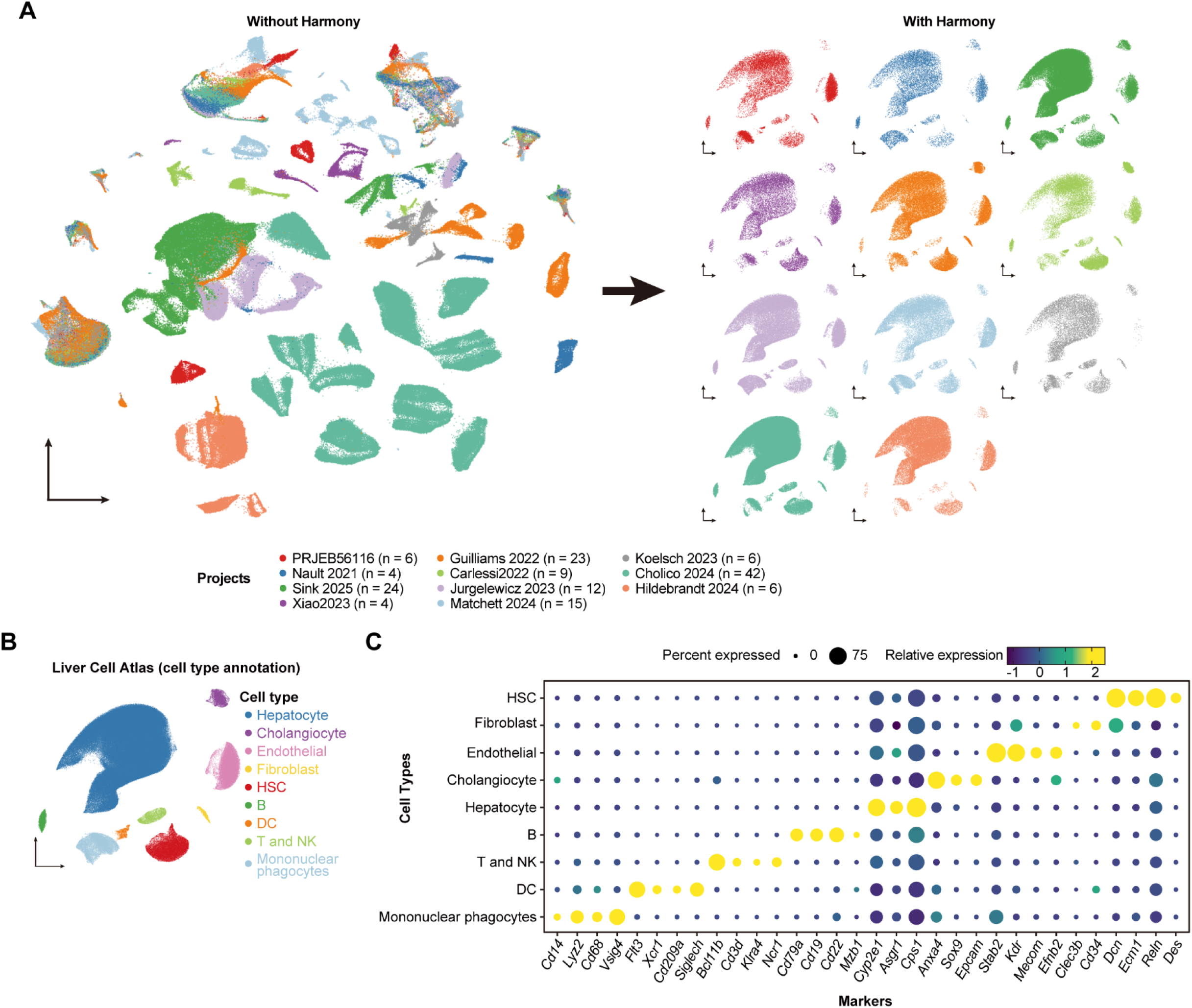
Integration of mouse liver snRNA-seq datasets for cell atlas construction. **(A)** Batch correction analysis demonstrating successful data integration across 11 snRNA-seq datasets (PRJEB56116, Guilliams 2022, Koelsch 2023, Nault 2021, Carlessi 2022, Cholico 2024, Sink 2025, Jurgelewicz 2023, Hildebrandt 2024, Xiao 2023, and Matchett 2024). Left: Pre-integration UMAP visualization showing substantial batch effects with dataset-specific clustering patterns. Right: Harmony integration displaying uniform cell distribution while preserving biological variation. Total integration encompasses 442,370 cells from 149 samples. **(B)** Integrated liver cell atlas from 11 mouse snRNA-seq datasets. Cell subtype annotation showing 9 identified cell types. **(C)** Cell type validation through canonical marker expression analysis. Dot plot displaying expression patterns of lineage-specific markers across 9 major liver cell populations. Clear segregation of markers confirms accurate cell type annotation following integration.

**Figure S21.**
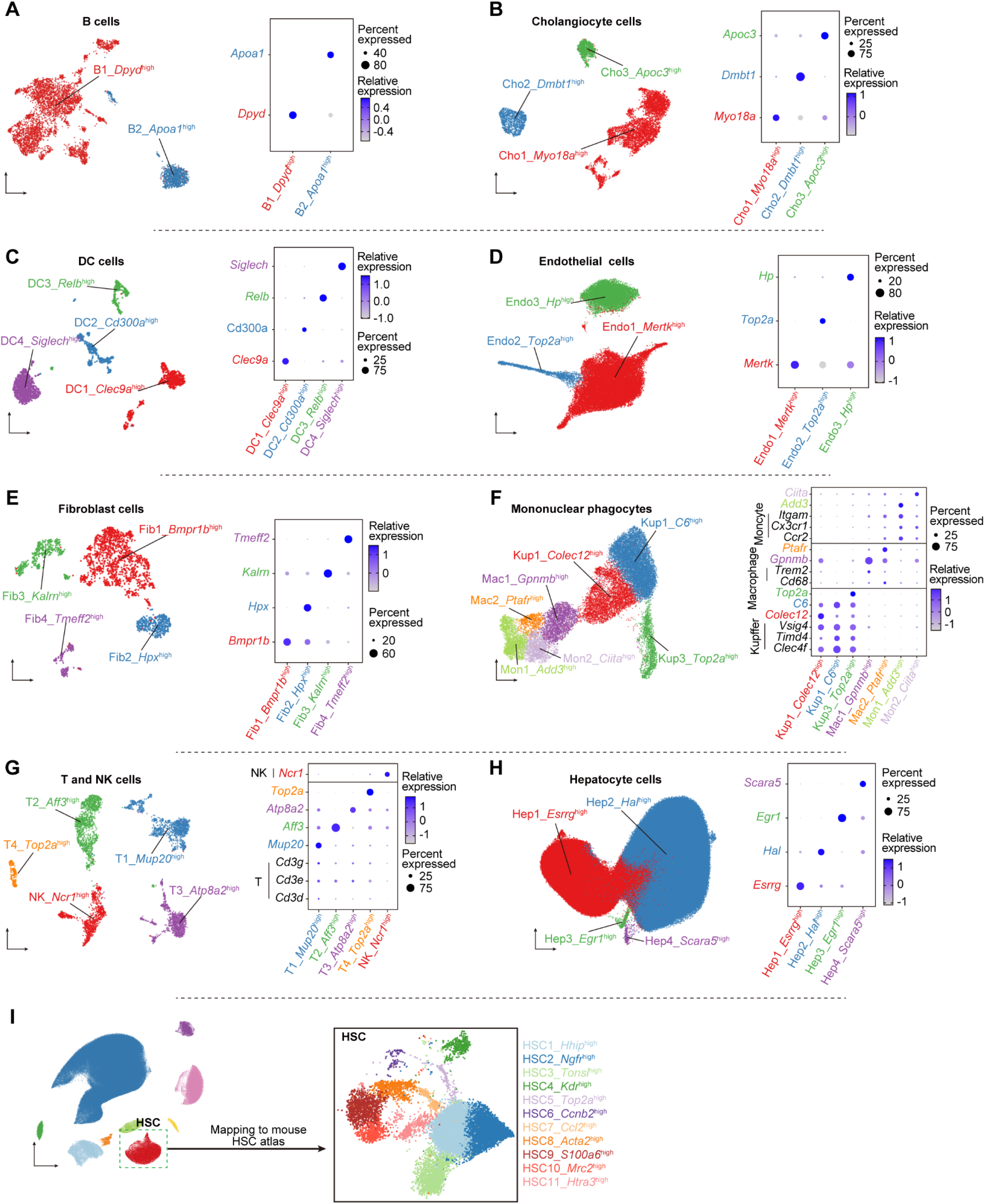
Comprehensive annotation of liver cell subtypes reveals cellular heterogeneity across major lineages. **(A-H)** B (A), Cholangiocyte (B), DC (C), Endothelial (D), Fibroblast (E), Mononuclear phagocyte (F), T/NK (G), and Hepatocyte (H) cell subset characterization. Left: Distinct populations identified. Right: Subtype-specific marker expression patterns. **(I)** HSC subtype integration with mouse atlas. Left: Overview of 11 HSC clusters within the integrated liver cell atlas. Right: Detailed UMAP of HSC populations mapped to the established mouse HSC atlas, confirming successful integration with the QAA trajectory framework. Abbreviations: Cho (Cholangiocytes), Hep (Hepatocytes), DC (Dendritic cells), B (B cells), T (T cells), NK (Natural killer cells), Kup (Kupffer cells), Mon (Monocytes), Mac (Macrophages), Fib (Fibroblast), Endo (Endothelial).

**Figure S22.**
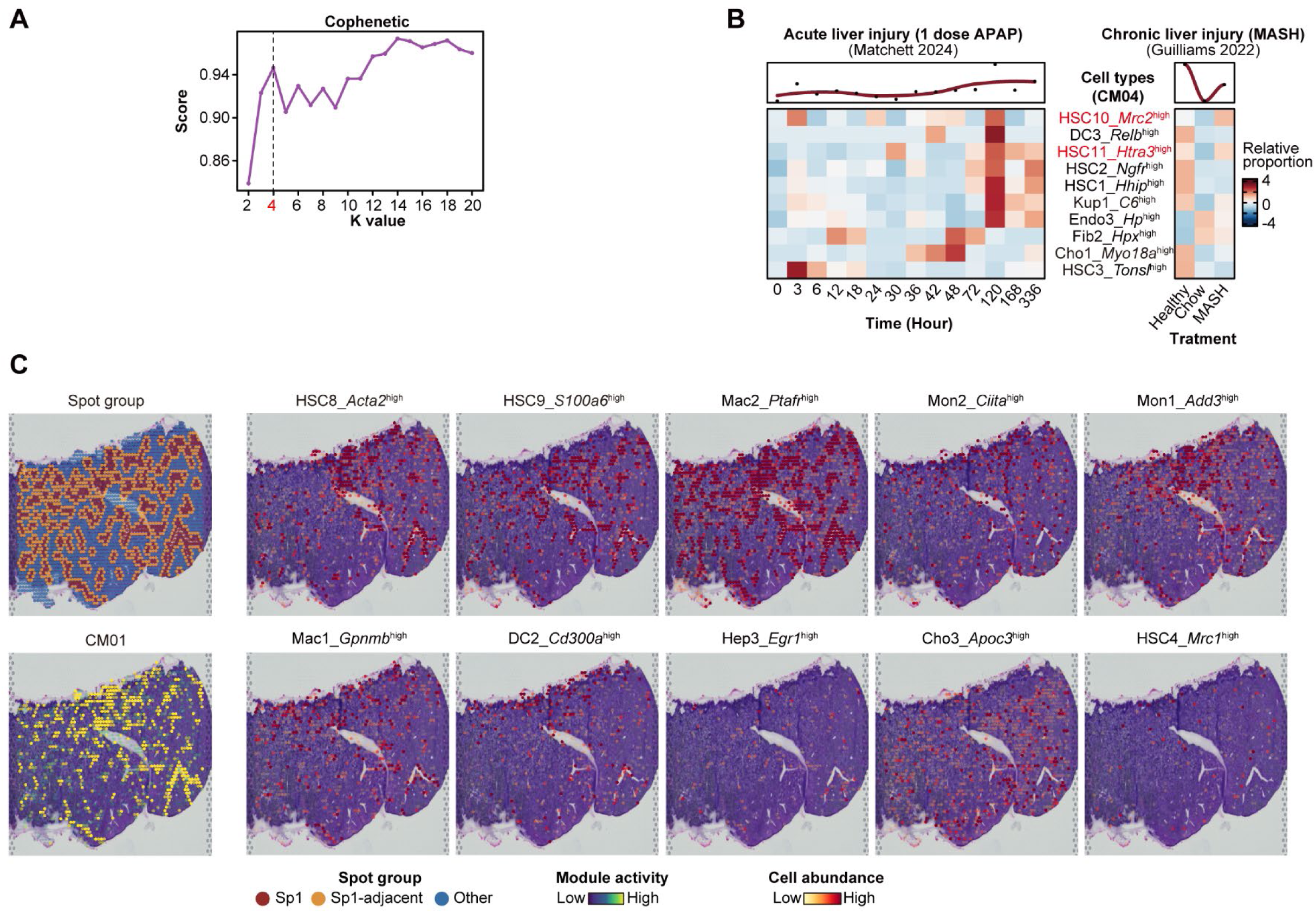
Cellular module dynamics and spatial organization in acute and chronic liver injuries. **(A)** Cophenetic analysis for optimal clustering of cellular modules. Plot showing cophenetic scores across different K values (2-20), with K = 4 selected as optimal based on score plateau, indicating robust clustering of cell type co-occurrence patterns into four distinct cellular modules. **(B)** Temporal dynamics of cell populations within cellular module CM04 during injury progression. Heatmaps comparing cell type proportions in acute liver injury versus chronic liver injury. **(C)** Spatial distribution patterns of CM01 module constituents in injured liver tissue. Representative spatial transcriptomic images from MASH liver (GSM5764421) showing spot group classification (Sp1, Sp1-adjacent, Other), module activity and spatial localization of CM01-associated cell populations. Abbreviations: Cho (Cholangiocytes), Hep (Hepatocytes), DC (Dendritic cells), B (B cells), T (T cells), NK (Natural killer cells), Kup (Kupffer cells), Mon (Monocytes), Mac (Macrophages), Fib (Fibroblast), Endo (Endothelial).

**Figure S23.**
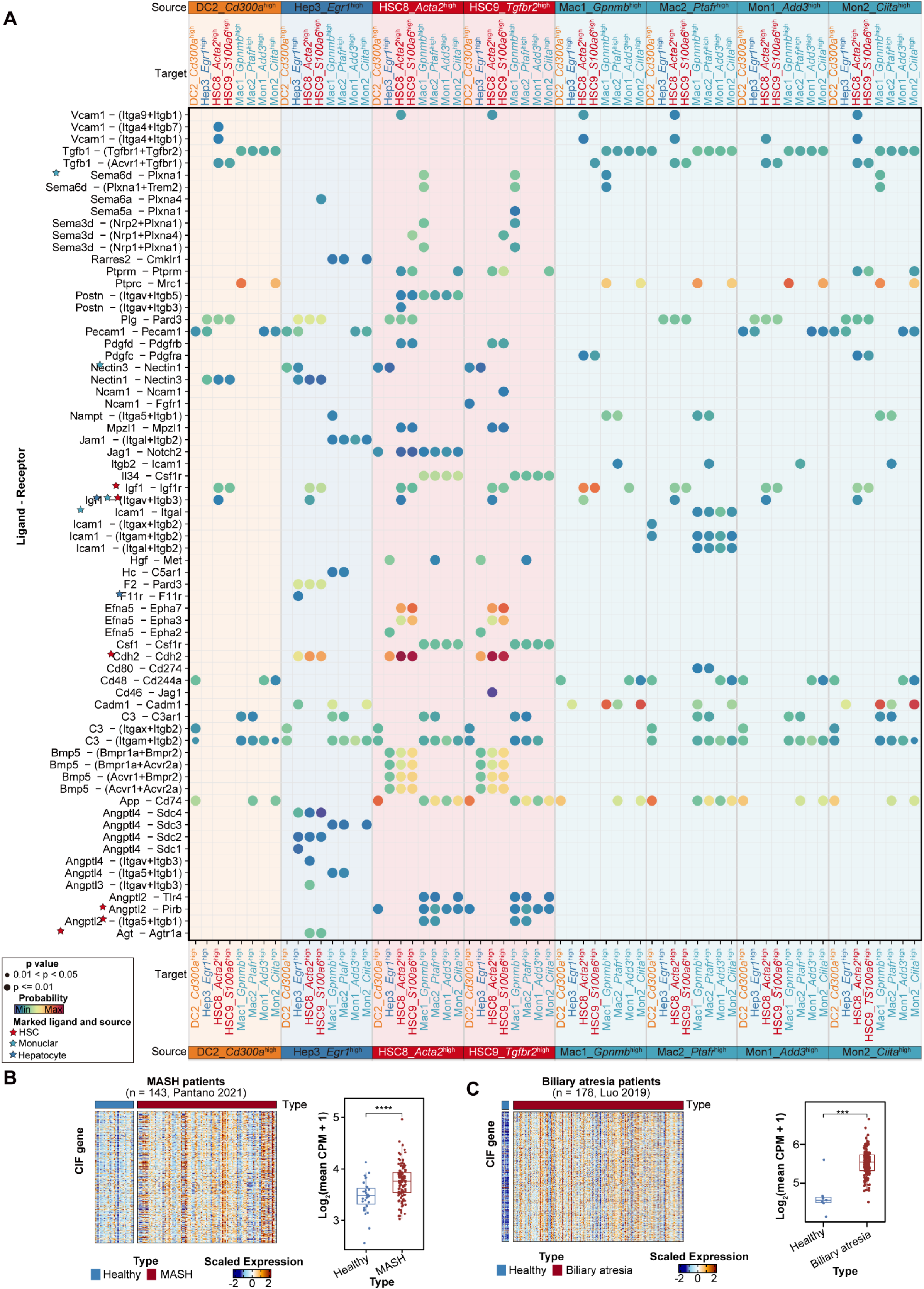
Intercellular communication networks and clinical validation of core inflammation-fibrosis signature. **(A)** Comprehensive ligand-receptor interaction analysis within CM01 module populations (excluding Cho3_*Apoc3*⁺ and HSC4_*Kdr*^high^). Dot plot displaying significant ligand-receptor pairs between source and target cell types. **(B)** Core inflammation-fibrosis gene signature validation in MASH patients (Pantano 2021). Left: Heatmap showing expression of 531-gene core inflammation-fibrosis gene signature across MASH patients. Samples ordered by disease severity with patient metadata indicated. Right: Quantification demonstrating significant elevation of CIF signature in MASH versus healthy controls (log₂ mean CPM + 1). **(C)** CIF signature validation in biliary atresia patients (Luo 2019). Left: Heatmap displaying CIF gene expression in atresia samples. Patients stratified by disease status with clinical annotations. Right: Box plot showing significantly elevated CIF signature in biliary atresia compared to healthy controls (log₂ mean CPM + 1). Abbreviations: Cho (Cholangiocytes), Hep (Hepatocytes), DC (Dendritic cells), B (B cells), T (T cells), NK (Natural killer cells), Kup (Kupffer cells), Mon (Monocytes), Mac (Macrophages), Fib (Fibroblast), Endo (Endothelial). Statistical significance is denoted as: ns, p > 0.05; *, p < 0.05; **, p < 0.01; ***, p < 0.001; ****, p < 0.0001 (unpaired two side Wilcoxon test).

## Supplemental Table Legend

Table S1.The list of datasets used in this study

Table S2.Enriched terms of markers in each HSC subtype

Table S3.Summary of cell type annotation in mouse HSC atlas

Table S4.Regulation modules related to HSC state transitions along the QAA trajectory

Table S5.The list of antibodies used in this study

Table S6.Primers of genes for qPCR

